# A Proximity-Driven Functional Screening Platform for the Discovery of Noninhibitory USP7 Ligand Enabling Targeted Protein Stabilization

**DOI:** 10.64898/2026.09.10.750156

**Authors:** Kate E. Young, Janitha Kumarathunga, Hiruni Kandambi, Vitoria Fernandes Ferro, Weijun Gui

**Affiliations:** Department of Chemistry, Syracuse University, 111 College Place. CST 1-133, Syracuse, NY 13244

**Keywords:** DUBTACs, USP7, chemically induced proximity

## Abstract

Targeted protein stabilization by deubiquitinase-targeting chimeras (DUBTACs) has emerged as a promising therapeutic strategy for preserving the function of essential proteins. DUBTACs are bifunctional molecules that recruit deubiquitinating enzymes (DUBs) to remove ubiquitin chains from substrate proteins, thereby preventing their proteasomal degradation. A key requirement for DUBTACs development is the identification of ligands that bind DUBs without impairing their catalytic activity. However, such noninhibitory ligands are exceedingly rare. Here, we report a novel proximity-driven on-bead functional screening platform for the discovery of noninhibitory DUB ligands. Using this approach, we discovered a previously unreported noninhibitory ligand for USP7. Furthermore, we generated DUBTACs by conjugating this USP7 ligand to a CFTR-binding ligand. Our lead compound, WJ045, effectively stabilized CFTR protein levels in cells, demonstrating the potential of this platform for discovering DUB recruiters and facilitating the development of targeted protein stabilization therapeutics.

## Introduction

Chemically induced proximity has transformed drug discovery by enabling the manipulation of protein function through induced interactions between two biomolecules.^1–6^ Among these modalities, proteolysis-targeting chimeras (PROTACs) represent the most successful example, recruiting E3 ubiquitin ligases to catalyze ubiquitination and subsequent proteasomal degradation of target proteins.^7^ The clinical advancement of targeted protein degradation by PROTACs and molecular glue degraders, has established induced protein degradation as a promising therapeutic modality for targeting disease-associated proteins that are inaccessible to conventional small-molecule inhibitors.^8–11^ The great success of PROTACs has inspired the development of other proximity-induced strategies such as lysosome-targeting chimera (LYTAC),^12^ regulated induced proximity targeting chimeras (RIPTACs),^13^ transcriptional/epigenetic chemical inducers of proximity (TCIPs),^14, 15^ and targeted protein posttranslational modifications.^16^

An emerging complementary application of chemically induced proximity is targeted protein stabilization by deubiquitinase-targeting chimeras (DUBTACs).^17–19^ Aberrant ubiquitination of essential proteins is a driver of many types of diseases and cancers.^20–22^ Deubiquitinating enzymes (DUBs) counteract the ubiquitination process by removing ubiquitin chains from substrate proteins.^23, 24^ Hijacking the enzymatic function of DUBs, DUBTACs rescue protein abundance by recruiting a DUB to remove ubiquitin chains from substrate proteins, thereby preventing proteasomal degradation. The first demonstration of this concept was reported in 2022 using OTUB1 as the recruited enzyme.^17^ Since then, additional DUBs, including USP7, USP28, and USP1, have been explored as potential recruiters.^25–28^ In parallel, DUBTACs have been applied to stabilize a growing number of therapeutically relevant proteins, including CFTR, Wee1, FOXO3A, p53, IRF3, AMPK, cGAS, PPARγ, UTX, PRCP, VHL and KEAP1.^17, 25–27, 29–33^

Despite these advances, the discovery of suitable DUB recruiters remains a major challenge. An ideal DUBTAC recruiter should bind a DUB while preserving its catalytic activity to enable target deubiquitylation. However, the vast majority of reported DUB ligands function as catalytic inhibitors, rendering them not ideal for targeted protein stabilization. This limitation largely stems from current ligand discovery strategies. Activity-based high-throughput screening (HTS) campaigns identify compounds that modulate DUB activity but cannot distinguish nonbinding molecules from ligands that bind without altering catalytic function. Conversely, affinity-based screening readily identifies DUB binders but provides no information on the functional consequences of ligand binding and therefore cannot differentiate noninhibitory recruiters from catalytic inhibitors. Consequently, neither approach is well suited for the discovery of functional DUB recruiters that both engage DUBs and preserve their enzymatic activity.

Here we report a proximity-driven, on-bead functional screening platform for the direct discovery of noninhibitory DUB ligands. Unlike conventional affinity-based screening approaches, our method selectively identifies ligands that both bind DUBs and preserve catalytic activity by coupling ligand recruitment to on-bead deubiquitylation. Using USP7 as a model system, we validate the platform with peptide ligands of known affinity, demonstrate its ability to distinguish noninhibitory binders from catalytic inhibitors, and discover a peptidomimetic USP7 ligand. Finally, we show that the ligand can be used to generate bifunctional molecules that stabilize the cystic fibrosis transmembrane conductance regulator (CFTR), establishing this platform as a general strategy for discovering functional DUB recruiters for targeted protein stabilization.

## Results and Discussion

### Establishment of a Proximity-Driven On-Bead Deubiquitylation Assay

We first sought to establish a functional assay capable of selectively identifying noninhibitory DUB ligands. Building on our previously reported proximity-driven on-bead assay for the discovery of noninhibitory E3 ubiquitin ligase ligands,^34^ we adapted this strategy for DUBs. The assay is based on the premise that a ligand that binds a DUB while preserving its catalytic activity will recruit the enzyme to the TentaGel bead surface, where it catalyzes the cleavage of a co-immobilized ubiquitin substrate (**Figure 1A**). In contrast, molecules that fail to bind the DUB or inhibit its catalytic activity should not induce ubiquitin hydrolysis.

To validate this concept, we constructed a model system that exploits the affinity interaction between His-tagged proteins and Ni–NTA as a surrogate noninhibitory ligand.^35, 36^ TentaGel beads were functionalized with Ni–NTA and co-displayed with 1% ubiquitin through Cu(I)-catalyzed azide–alkyne cycloaddition,^37, 38^ yielding TG-Ub-NTA-Ni beads (**Figure 1B**). As a negative control, analogous beads lacking Ni² (TG-Ub-NTA) were prepared. The synthetic procedures of the TG-Ub-NTA-Ni and TG-Ub-NTA beads are described in **Scheme S1&S2**.

Purified His-tagged full length USP7 was then incubated with the beads for 6 hours at room temperature, followed by washing, staining with Alexa Fluor 647-conjugated anti-ubiquitin antibody, and flow cytometric analysis. Cleavage of bead-bound ubiquitin decreases antibody labeling and therefore reduces fluorescence intensity. TG-Ub-NTA-Ni beads exhibited a clear, concentration-dependent reduction in fluorescence following incubation with His-USP7 (**Figure 1C**), whereas TG-Ub-NTA beads showed no detectable response over the same concentration range (**Figure 1D**). These results demonstrate that enzyme recruitment through a noninhibitory interaction efficiently promotes proximity-driven deubiquitylation.

**Figure 1:**
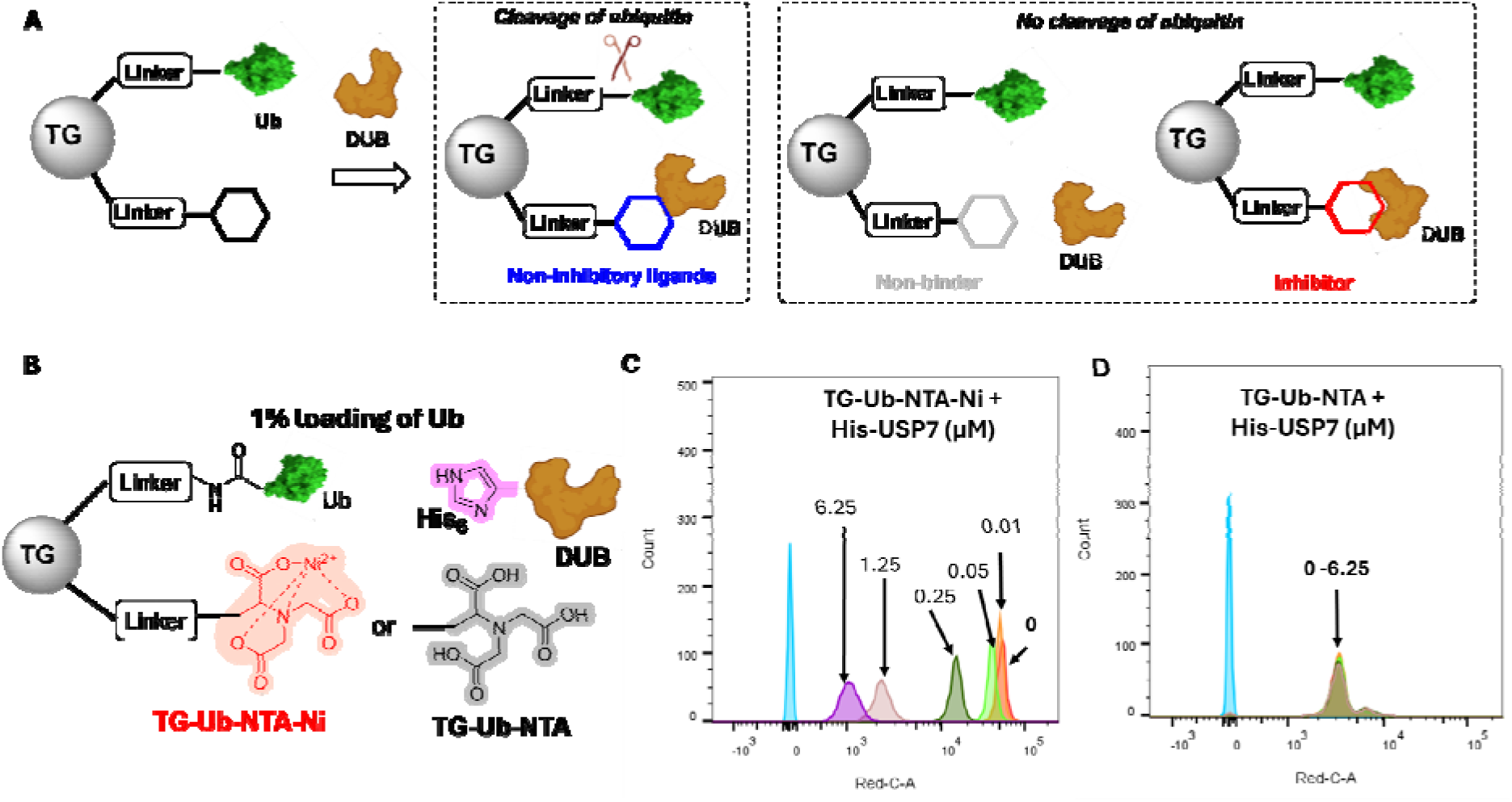
Establishment of a proximity-driven on-bead deubiquitylation assay. (**A**) Schematic illustration of the proximity-driven functional screening platform for the discovery of noninhibitory DUB ligands. (**B**) The chemical structure of TG-Ub-NTA-Ni and TG-Ub-NTA beads. TentaGel beads were functionalized with Ni–NTA or NTA and co-displayed with 1% ubiquitin to generate TG-Ub-NTA-Ni and TG-Ub-NTA, respectively. (**C&D**) Flow cytometry analysis of Alexa Fluor 647 fluorescence (Red C channel with emission wavelength of 670/14 nm) of TG-Ub-NTA-Ni (**C**) and TG-Ub-NTA (**D**) beads containing 1% ubiquitin after incubation with the indicated concentrations of full-length USP7 (USP7 FL) for 6 hours at room temperature, followed by staining with an Alexa Fluor 647-conjugated anti-ubiquitin antibody.

To evaluate the dynamic range of the assay, we reduced the ubiquitin loading from 1% to 0.2%. Despite the lower substrate density, TG-Ub-NTA-Ni beads continued to exhibit a robust concentration-dependent fluorescence decrease (**Figure S1**), indicating that the assay remains sensitive over a broad range of substrate loading. When the ubiquitin loading was further reduced to 0.04%, a concentration-dependent decrease in fluorescence intensity was still observed following incubation with USP7. However, the NTA beads exhibited weak staining at this low loading level, limiting signal quality (**Figure S1**).

We next examined whether this strategy could be generalized to additional DUB families. Similar concentration-dependent responses were observed with USP7 catalytic domain (USP7CD), UCHL1, UCHL3, USP14, and OTUB1 (**Figure S2-S4**), demonstrating that the platform is broadly applicable to mechanistically diverse DUBs.

### Validation of the Proximity-Driven on-Beads Deubiquitylation Assay Using Known USP7 Ligands

Having established the assay, we next investigated whether it could distinguish authentic noninhibitory ligands from catalytic inhibitors. The human proteome encodes approximately 100 deubiquitinating enzymes (DUBs), which exhibit diverse substrate and ubiquitin linkage specificities.^23^ Among these, K48-linked polyubiquitin chains serve as the canonical signal for proteasomal degradation.^39, 40^ We selected USP7 as a model system for several reasons. First, USP7’s activity to cleave K48-linked ubiquitin chains is well-characterized.^41^ Second, USP7 contains an N-terminal TRAF domain that is unique among the DUB family,^42^ providing an attractive site for selective ligand recruitment without directly targeting the catalytic domain. Third, the TRAF domain of USP7 recognizes peptide ligands containing the consensus P/AxxS motif, including the well-characterized PSTS peptide, which binds USP7 with a dissociation constant (Kd) of approximately 10 μM.^43, 44^ This well-defined, noninhibitory interaction provides an ideal positive control for validating the screening platform.

To validate the assay, TentaGel beads co-displaying 1% ubiquitin and the PSTS peptide were synthesized (**Figure 2A**) and incubated with increasing concentrations of full-length USP7 under the same assay conditions. The synthetic detail is described in **Scheme S3**. Following antibody staining and flow cytometric analysis, a concentration-dependent decrease in fluorescence was observed (**Figure 2B**), indicating efficient recruitment of USP7 to the bead surface and subsequent cleavage of immobilized ubiquitin.

**Figure 2:**
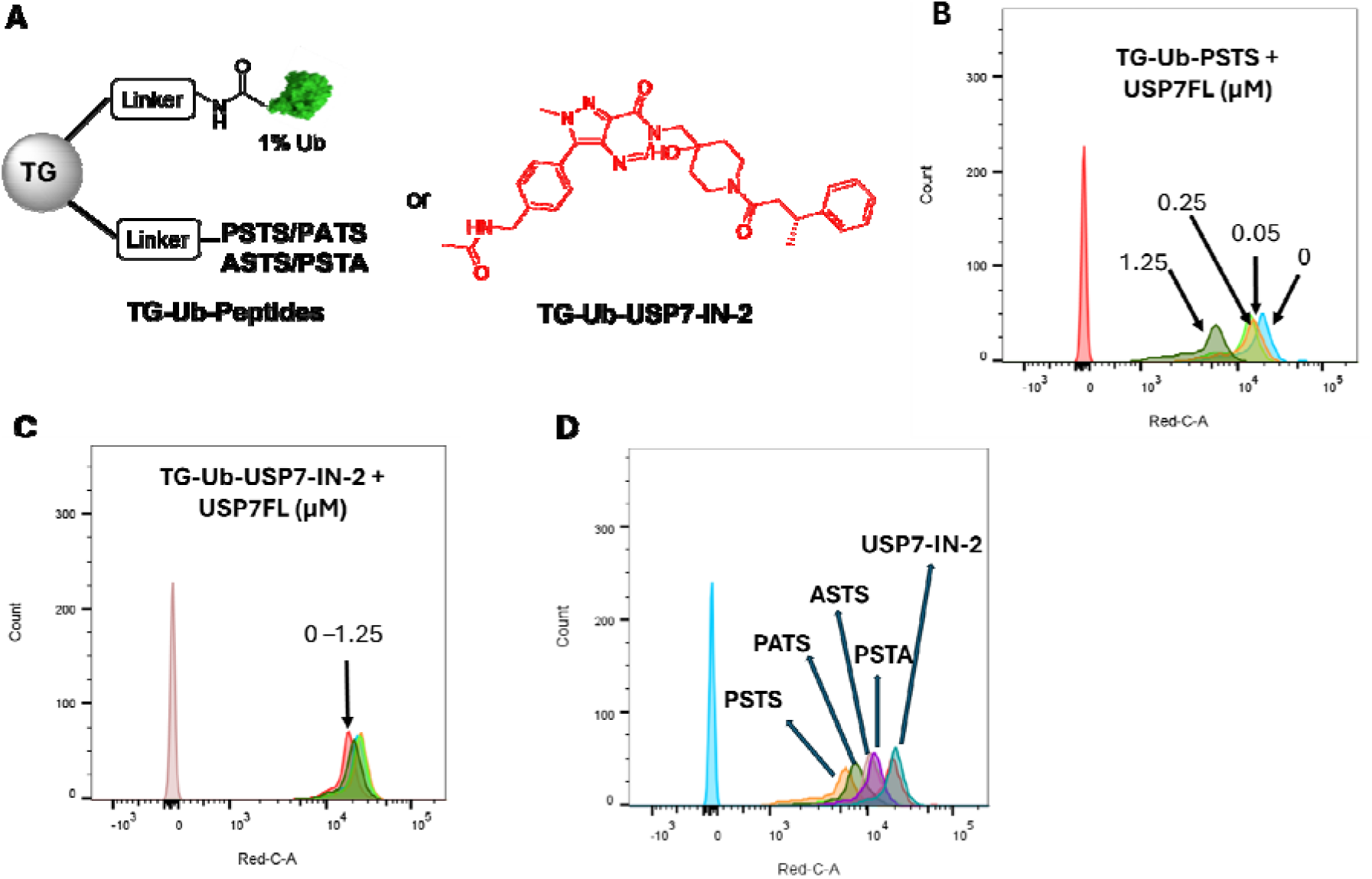
Validation of the proximity-driven on-bead deubiquitylation assay using known USP7 ligands. (**A**) TentaGel beads were functionalized with USP7 peptide ligands or inhibitor USP-IN-2 and co-displayed with 1% ubiquitin. (**B**&**C**) Flow cytometry analysis of Red C signal of beads TG-Ub-PSTS (B) and TG-Ub-USP7-IN-2 (C) with 1% loading of ubiquitin after incubation with different concentrations of USP7 FL as indicated at room temperature for 6 hours, stained with Alexa Fluor 647 labeled anti-ubiquitin. (**D**) Flow cytometry analysis of Red C signal of beads different USP7 ligands and 1% loading of ubiquitin after incubation with 1 μM of USP7 FL at room temperature for 6 hours, stained with Alexa Fluor 647 labeled anti-ubiquitin.

To determine whether catalytic inhibition abolishes this response, we synthesized beads displaying the USP7 inhibitor, USP7-IN-2, and 1% ubiquitin.^45^ In contrast to PSTS, no measurable decrease in fluorescence was detected after incubation with USP7 FL (**Figure 2C**), demonstrating that enzyme binding alone is insufficient to generate a positive signal when catalytic activity is blocked. These findings establish that the assay selectively identifies ligands that preserve DUB activity rather than simply bind the enzyme.

We next examined whether the extent of on-bead deubiquitylation reflects ligand binding affinity. Thus, we synthesized beads displaying three mutant peptides (PATS, ASTS, and PSTA). All three mutant peptides retained measurable activity but produced substantially smaller decreases in fluorescence after incubation with 1 μM USP7 FL compared with the parent PSTS sequence (**Figure 2D**). Mutations of the N-terminal proline (ASTS) or the C-terminal serine (PSTA) caused the greatest loss of activity, consistent with previous reports identifying these residues as the principal determinants of USP7 recognition.^43^ As expected, beads displaying USP7-IN-2 remained inactive.

Together, these results demonstrate that the extent of proximity-driven deubiquitylation is directly related to ligand affinity while simultaneously requiring an intact catalytic mechanism. The assay therefore provides a functional readout that integrates both DUB binding and catalytic competence, enabling the selective discovery of noninhibitory DUB ligands.

### Discovery of Noninhibitory USP7 Ligands Using the On-Bead Functional Assay

Having established the proximity-driven deubiquitylation assay, we next explored its utility for the discovery of noninhibitory USP7 ligands suitable for DUBTACs development. Although the PSTS peptide efficiently recruits the USP7 TRAF domain, its peptide nature renders it susceptible to intracellular proteolysis and therefore unsuitable as a DUBTAC recruiter.

Peptidomimetic scaffolds, which preserve peptide-like recognition features while offering significantly enhanced permeability, stability, and drug-like properties.^46, 47^ We therefore sought to identify peptidomimetic ligands that retain productive USP7 binding while exhibiting improved chemical stability.

Previous structural studies, together with our mutagenesis results, identified the serine residue within the PSTS motif as a critical determinant of USP7 recognition.^42, 43, 48^ Guided by these observations, we designed a focused peptidomimetic library by systematically modifying the residues flanking the essential serine while preserving the key recognition elements required for TRAF-domain binding.

We first investigated substitutions on the N-terminal side of the serine by synthesizing a one-bead-one-well library containing 71 distinct capping groups in 96-well plate (**Figure 3A**). The chemical structures of the 71 capping acids are shown in **Figure S5**. Each compound was co-displayed with 1% ubiquitin on TentaGel beads and screened against 1.25 μM full-length USP7 using the proximity-driven deubiquitylation assay. Parallel control experiments performed in the absence of USP7 established the baseline fluorescence for each bead population, allowing the extent of ubiquitin cleavage to be quantified as the percentage decrease in fluorescence intensity.

Among the 71 analogues evaluated, the ligand incorporating 3-indoleacetic acid produced the strongest activity, resulting in a 16% decrease in fluorescence intensity (**Figure 3B**). To further examine the structural importance of the serine residue, we synthesized additional analogues in which the serine was replaced with D-serine, O-methyl serine, or homoserine while maintaining the optimal N-terminal substituent (**Figure 3B**). Each modification reduced assay activity, yielding fluorescence decreases of only 12%, 10%, and 9%, respectively. These findings further establish the central role of the native serine residue in productive recognition of the USP7 TRAF domain.

**Figure 3:**
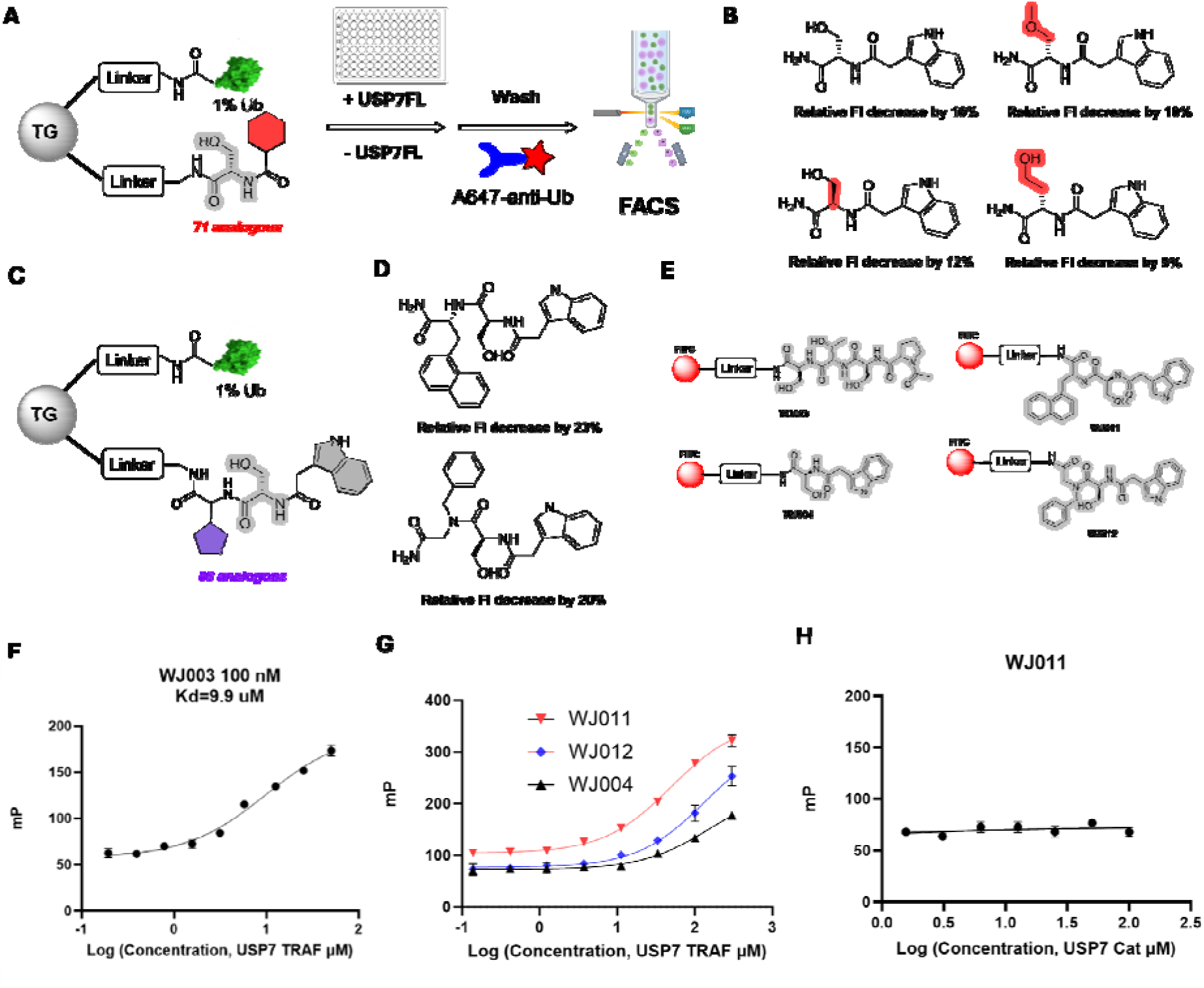
Discovery of noninhibitory USP7 ligands using the on-bead functional assay. (**A**) The screening platform to discover noninhibitory ligands for USP7 FL. (**B**) The chemical structure of the hit ligand and its derivatives. The relative fluorescent intensity decrease of the respective beads after incubation with 1 μM of USP7 FL at room temperature for 6 hours, stained with Alexa Fluor 647 labeled anti-ubiquitin. (**C**) The representative chemical structure of the beads displaying 88 different analogous on the C-terminal of serine. (**D**) The chemical structure of the ligands displayed on the beads and the activity to decrease the relative fluorescent intensity of the beads after incubation with 1 μM of USP7 FL at room temperature for 6 hours, stained with Alexa Fluor 647 labeled anti-ubiquitin. (**E**) The chemical structure of the fluorescent probes. (**F**&**G**&**H**) fluorescence polarization assay to measure the binding affinity of the probe to the protein as indicated.

To further optimize ligand affinity, we next diversified the C-terminal side of the serine by constructing a second focused library containing 88 unnatural amino acids while retaining the 3-indoleacetic acid substituent identified in the initial screen (**Figure 3C**). The chemical structure of 88 unnatural amino acids is shown in **Figure S6.** Screening of this library identified top two lead compounds (**Figure 3D**) that produced fluorescence decreases of 23% and 20% respectively, representing a marked improvement over the first-generation analogues and demonstrating that iterative optimization of both sides of the pharmacophore effectively enhances USP7 recruitment.

Because the on-bead assay provides a functional rather than direct binding readout, we next sought to independently validate ligand engagement using fluorescence polarization assay. Four fluorescein-labeled probes (**WJ003**, **WJ004**, **WJ011**, and **WJ012**) were synthesized by conjugating fluorescein isothiocyanate (FITC) to the corresponding ligands (**Figure 3E**). The synthesis and characterization of fluorescent probes are described in **Scheme S4**. Consistent with previous reports,^43^ the parent PSTS peptide bound the USP7 TRAF domain with a dissociation constant of 9.9 μM (**Figure 3F**). Among the optimized ligands, **WJ011** displayed the highest affinity with a Kd around 49 μM, followed by **WJ012** and **WJ004** (**Figure 3G**). Notably, this affinity ranking closely paralleled the activities observed in the on-bead deubiquitylation assay, supporting the ability of the screening platform to identify authentic USP7 binders.

To confirm the binding specificity of the optimized ligands, **WJ011** was evaluated against the isolated catalytic domain of USP7 (USP7CD), which lacks the N-terminal TRAF domain. No measurable binding was detected (**Figure 3H**), confirming that ligand recognition is mediated specifically through the TRAF domain rather than the catalytic domain.

### Lead USP7 Ligand WJ021 Preserves USP7 Catalytic Activity

In order to evaluate whether the lead ligand inhibits the catalytic activity of USP7. **WJ021** as shown in **Figure 4A** was synthesized (**Scheme S5)**. We first characterized the enzymatic activities of full-length USP7 (USP7 FL) and USP7 catalytic domain (USP7 CD) using the ubiquitin-rhodamine fluorogenic substrate. Following incubation of 1 μM ubiquitin-rhodamine with increasing concentrations of USP7 FL or USP7 CD for 3 hours at room temperature, fluorescence was measured to quantify substrate hydrolysis (**Figure 4B**). Both USP7 FL and USP7 CD exhibited concentration-dependent hydrolysis of the ubiquitin-rhodamine substrate, with full-length USP7 displaying higher catalytic activity than the catalytic domain.

**Figure 4:**
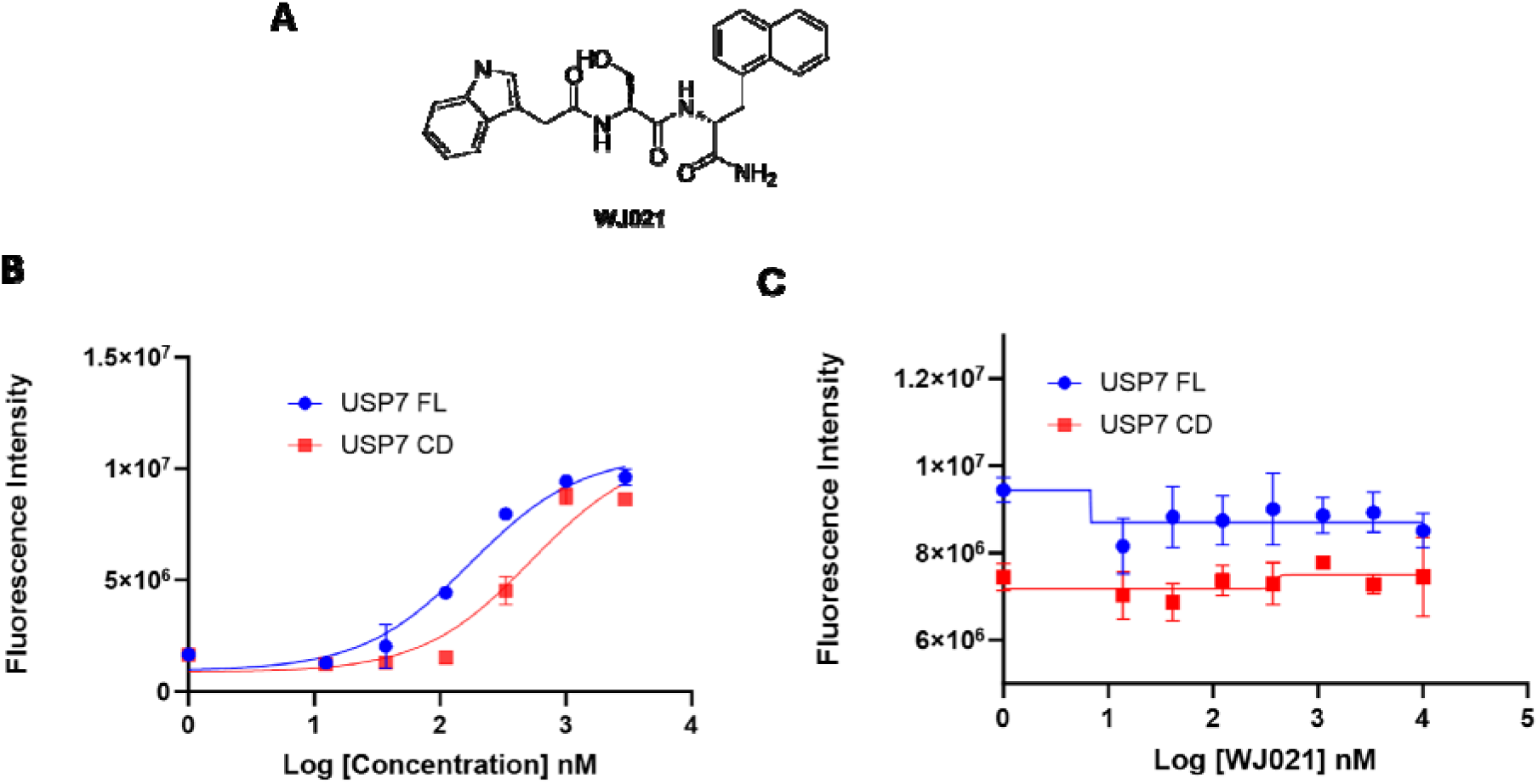
WJ021 preserves USP7 catalytic activity. (**A**) The chemical structure of WJ021. (**B**) Characterization of the enzymatic activities of USP7 FL and USP7 CD using the ubiquitin-rhodamine fluorogenic substrate. (**C**) Measurement of influence of WJ021 activity in USP7 FL or USP7CD hydrolyzing ubiquitin-rhodamine fluorogenic substrate.

We next assessed the effect of **WJ021** on USP7 activity by incubating 1 μM USP7 FL or USP7 CD with increasing concentrations of **WJ021** at room temperature, followed by measurement of ubiquitin-rhodamine hydrolysis. **WJ021** did not measurably inhibit the activity of either USP7 FL or USP7 CD under any of the conditions tested (**Figure 4C**), demonstrating that the optimized ligand preserves the catalytic activity of USP7. These results establish **WJ021** as a noninhibitory USP7 recruiter suitable for DUBTAC development.

Collectively, these results demonstrate that the proximity-driven on-bead assay enables the functional discovery and optimization of noninhibitory USP7 ligands. Furthermore, the strong correlation between on-bead deubiquitylation and independent binding measurements validates the assay as a robust platform for identifying DUB recruiters suitable for targeted protein stabilization.

### Development of USP7-Recruiting DUBTACs for Targeted Stabilization of CFTR

Having identified a noninhibitory USP7 ligand, we next investigated whether this ligand could function as USP7 recruiter in bifunctional molecules for targeted protein stabilization. As a proof of concept, we selected the cystic fibrosis transmembrane conductance regulator (CFTR) as the target protein because its degradation contributes to the pathogenesis of cystic fibrosis and restoration of CFTR stability represents a clinically validated therapeutic strategy.^49, 50^

A series of DUBTACs were synthesized by conjugating the USP7 ligand (**WJ020**) to a reported CFTR-binding ligand through linkers of varying length and composition (**Figure 5A**). The synthesis scheme of **WJ020** and bifunctional DUBTACs are described in **Scheme S6&S7**.

The resulting compounds were evaluated for their ability to stabilize CFTR in cells by Western blot analysis. The ΔF508-CFTR mutant protein in the CFBE41o4.7 ΔF508-CFTR expressing human cystic fibrosis bronchial epithelial cells were treated with the compounds at 10 μM concentration for 24 hours. DMSO and NJH-2-056 were used as negative and positive controls respectively.^17^ Among the analogues examined, the most potent compound **WJ045** produced 11.6 fold of increase of CFTR protein abundance compared to the DMSO control, whereas compounds containing suboptimal linkers exhibited substantially weaker activity (**Figure 5B**). As comparison, **WJ021** did not show any detectable increase of CFTR level, which confirms that CFTR stabilization resulted from USP7-mediated deubiquitylation rather than nonspecific effects. Moreover, USP7-based CFTR DUBTAC **WJ045** stabilized CFTR in a concentration dependent manner (**Figure 4D**). Treatment of CFBE41o4.7 ΔF508-CFTR with 20 μM **WJ045** for 24 hours increased CFTR protein levels by 29.1-fold compared with the DMSO control. Consistent with a concentration-dependent effect, lower concentrations of **WJ045** resulted in progressively reduced CFTR stabilization. **WJ045** also increased CFTR protein levels in a time-dependent manner. Treatment with 20 μM **WJ045** for 8 hours increased CFTR levels by 5.7-fold, whereas 24 hours of treatment resulted in a 21.2-fold increase compared with the DMSO control.

**Figure 5:**
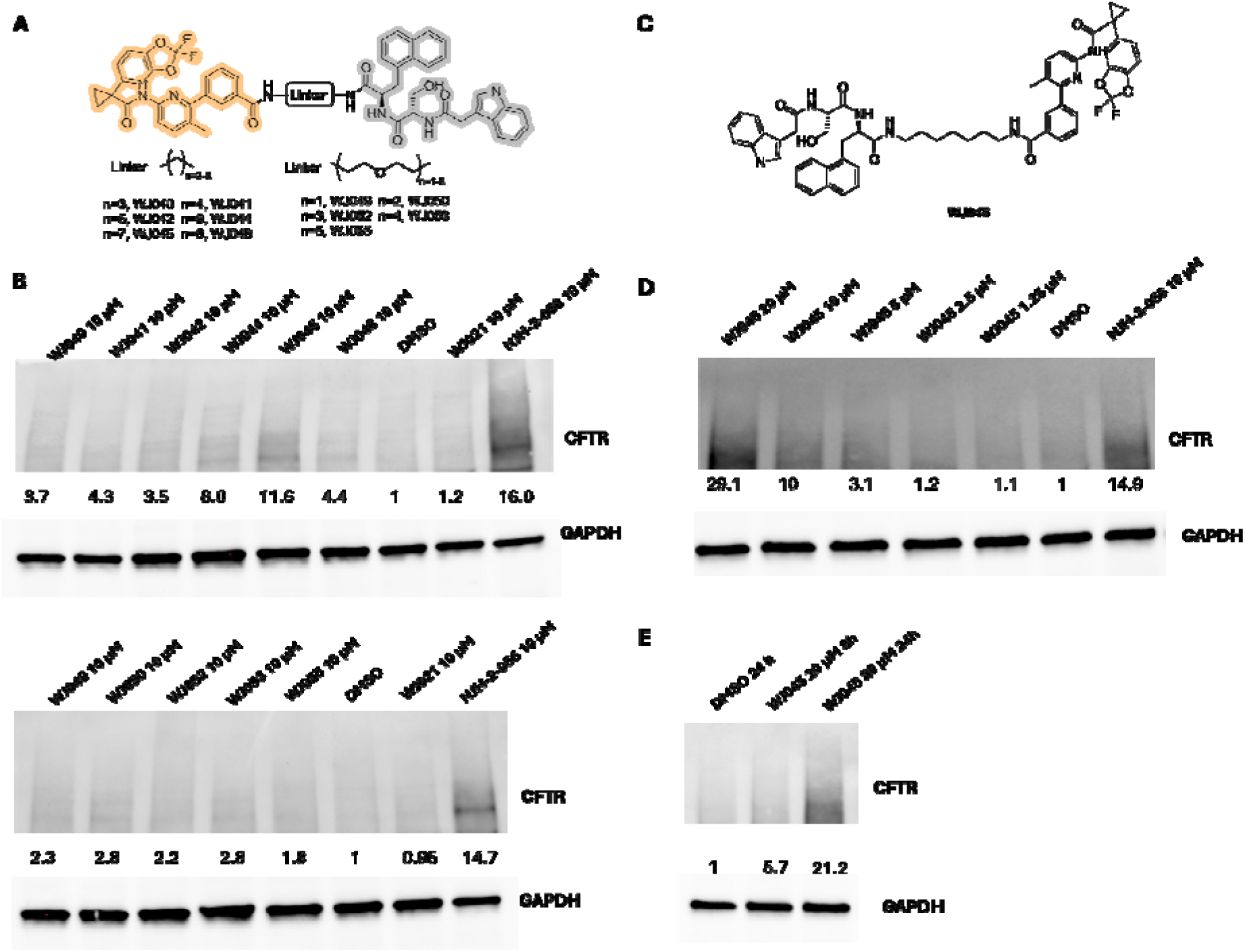
USP7-recruiting DUBTACs for targeted stabilization of CFTR. (**A**) The representative chemical structure of USP7-based CFTR DUBTACs. (**B**) The human cystic fibrosis bronchial epithelial CFBE41o-4.7 ΔF508-CFTR cells were treated with 10 μM of the indicated compounds for 24 hours, with NJH-2−057 as a positive control, followed by Western blot analysis. (**C**) The chemical structure of WJ045. (**D**) CFBE41o-4.7 ΔF508-CFTR cells were treated with the indicated concentrations of WJ045 for 24 hours, followed by Western blot analysis. (**E**) CFBE41o-4.7 ΔF508-CFTR cells were treated with 20 μM WJ045 or DMSO for the indicated hours, followed by Western blot analysis.

Collectively, these studies demonstrate that the noninhibitory USP7 ligand discovered through our functional screening platform can be successfully incorporated into bifunctional molecules that induce targeted protein stabilization. More broadly, these results provide proof of concept that proximity-driven functional screening can directly identify DUB recruiters suitable for DUBTAC development, thereby overcoming one of the principal challenges limiting the expansion of targeted protein stabilization technologies.

## Discussion

Chemically induced proximity has rapidly expanded the scope of drug discovery by enabling the selective modulation of protein function through induced protein–protein interactions. Although targeted protein degradation has emerged as a broadly applicable therapeutic modality, its complementary strategy, targeted protein stabilization, remains comparatively underdeveloped. DUBTACs, which harness DUBs to remove ubiquitin chains from target proteins, represent a promising approach for achieving targeted protein stabilization. However, a major challenge in DUBTACs development is the limited availability of DUB ligands that engage their target without compromising catalytic activity.

Although DUBTACs constructed using reversible DUB inhibitors have been shown to promote target protein stabilization in cells.^25–28^ A possible explanation is that a reversible inhibitor, when incorporated into a DUBTAC, can recruit and locally concentrate the DUB near the target protein while still permitting sufficient catalytic activity to remove ubiquitin chains.^25^ Consistent with this concept, DUBTACs incorporating weak USP1 inhibitors have been reported to outperform those containing more potent inhibitors,^27^ highlighting the importance of preserving DUB catalytic function. Thus, an ideal DUBTAC ligand should efficiently engage the DUB while minimally perturbing its enzymatic activity.

In this work, we addressed this challenge by developing a proximity-driven, on-bead functional screening platform that directly selects for noninhibitory DUB ligands. Unlike traditional affinity-based screening methods, our assay couples ligand binding with enzyme function, allowing only catalytically competent DUBs to generate a positive signal. This functional selection distinguishes productive recruiters from inactive binders or catalytic inhibitors and therefore provides a direct solution to a central bottleneck in DUBTAC development.

The design of the assay is conceptually simple yet broadly applicable. By co-displaying ubiquitin and candidate ligands on the same bead, productive DUB recruitment creates a local increase in enzyme concentration that promotes cleavage of the immobilized ubiquitin substrate. The resulting decrease in fluorescence provides a quantitative functional readout of ligand activity. Importantly, this readout integrates two essential properties required for DUBTAC development—enzyme binding and preservation of catalytic activity—within a single screening assay. This dual selection represents a significant advantage over conventional biophysical approaches, which typically evaluate binding and enzymatic activity independently.

Several observations support the robustness of the platform. First, the assay successfully distinguished known noninhibitory peptide ligands from a catalytic USP7 inhibitor, demonstrating excellent functional specificity. Second, systematic mutagenesis of the USP7-binding peptide revealed a strong correlation between ligand affinity and on-bead deubiquitylation efficiency, indicating that the fluorescence signal reflects productive molecular recognition rather than nonspecific enzyme adsorption. Finally, fluorescence polarization measurements independently validated the binding affinities of the optimized ligands and closely paralleled the activities observed in the functional assay. Together, these findings establish that proximity-driven deubiquitylation serves as a reliable surrogate for productive DUB recruitment.

An important feature of this strategy is that it enables functional ligand discovery rather than merely affinity optimization. Most screening approaches identify compounds that bind the target protein, after which extensive secondary assays are required to determine whether binding perturbs protein function. In contrast, our assay incorporates catalytic competence into the primary selection step, substantially reducing the likelihood of advancing inhibitors as lead compounds. This feature is particularly advantageous for enzymes whose catalytic activity must be preserved for therapeutic applications.

The utility of the platform is further demonstrated by the successful development of USP7-recruiting DUBTACs capable of stabilizing CFTR in cells. The observation that optimized bifunctional molecule increased CFTR abundance supports the central premise that noninhibitory USP7 ligands identified by our screening strategy can function as productive DUB recruiters in a cellular context. More broadly, these results establish a direct workflow from functional ligand discovery to DUBTAC development, providing an efficient strategy for expanding the repertoire of DUB recruiters available for targeted protein stabilization.

Although this study focuses on USP7, the assay design is not inherently restricted to a single DUB. We demonstrate compatibility with multiple members of the USP, UCH, and OTU families, suggesting that the platform should be broadly adaptable across mechanistically diverse deubiquitinases. Because the assay relies only on enzyme-mediated substrate cleavage, it should also be readily modified for high-throughput screening of genetically encoded libraries, one-bead-one-compound collections, or DNA-encoded chemical libraries. Such adaptations could substantially accelerate the discovery of functional DUB recruiters for both established and previously unexplored deubiquitinases.

Despite these advances, several limitations warrant consideration. First, the best compound **WJ045** is not as potent as positive control NJH-2-056. But our result demonstrated that engaging the pocket outside the catalytic domain of USP7 can generate DUBTACs. Further optimization of the ligands will very likely increase the potency. Moreover, the current platform primarily evaluates ligand function using purified proteins under defined in vitro conditions and therefore does not directly account for intracellular factors such as protein localization, conformational dynamics, or competition with endogenous binding partners. Furthermore, successful DUBTAC activity depends not only on recruiter affinity but also on productive ternary complex formation, linker architecture, and target accessibility. Future studies integrating structural characterization, quantitative ternary complex analysis, and cellular proteomics will provide deeper insight into the molecular determinants governing efficient targeted protein stabilization.

In summary, we describe a functional screening strategy that directly identifies noninhibitory DUB ligands suitable for targeted protein stabilization. By integrating enzyme recruitment and catalytic activity into a single assay, this platform overcomes a major limitation of existing ligand discovery approaches and establishes a general framework for developing next-generation DUBTACs. Importantly, the optimized noninhibitory USP7 ligand was successfully incorporated into bifunctional DUBTACs that stabilized CFTR in cells, providing proof of concept that functional DUB recruiters identified through this platform can be directly translated into targeted protein stabilization. Beyond USP7, the assay is readily adaptable to other deubiquitinases and should facilitate the discovery of functional recruiters for expanding the DUBTAC toolbox. More broadly, this work establishes a general paradigm for functional ligand discovery in chemically induced proximity, in which catalytic competence is integrated into the primary screening process. We anticipate that this strategy will accelerate the development of next-generation targeted protein stabilization technologies and broaden the therapeutic potential of chemically induced proximity.

## ASSOCIATED CONTENT

## AUTHOR INFORMATION

### Author Contributions

The manuscript was written through contributions of all authors. All authors have given approval to the final version of the manuscript.

