## Supporting Information for "A Proximity-Driven Functional Screening Platform for the Discovery of Noninhibitory USP7 Ligand Enabling Targeted Protein Stabilization"

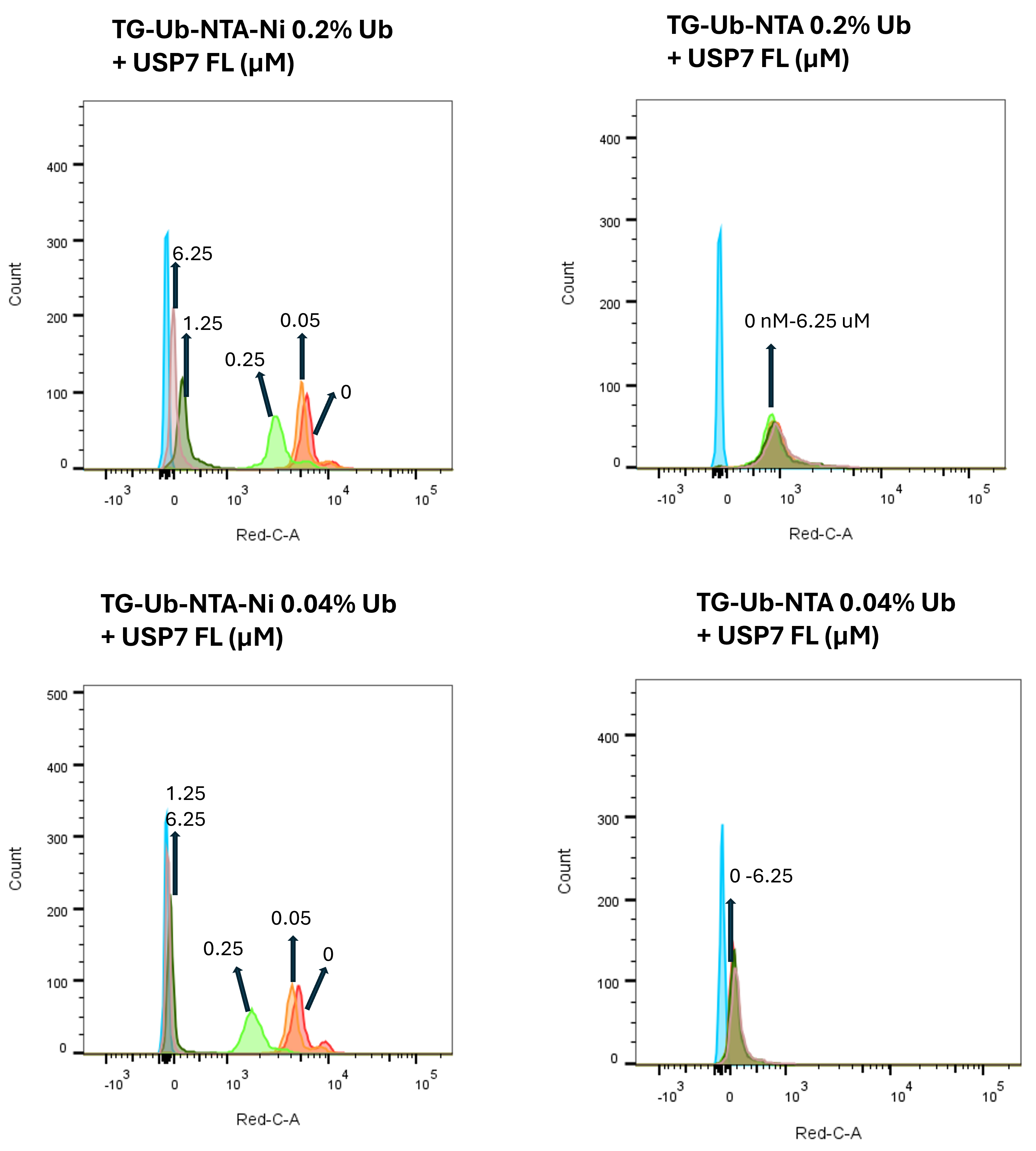

**Figure S1**: Flow cytometry analysis of Red C signal (Red C channel with emission wavelength of 670/14 nm) of beads (TG-Ub-NTA-Ni or TG-Ub-NTA) with different loading of ubiquitin (0.2%, 0.04%) after incubation with different concentrations of USP7 FL indicated at room temperature for 6 h, stained with Alexa Fluor 647 labeled anti-ubiquitin.

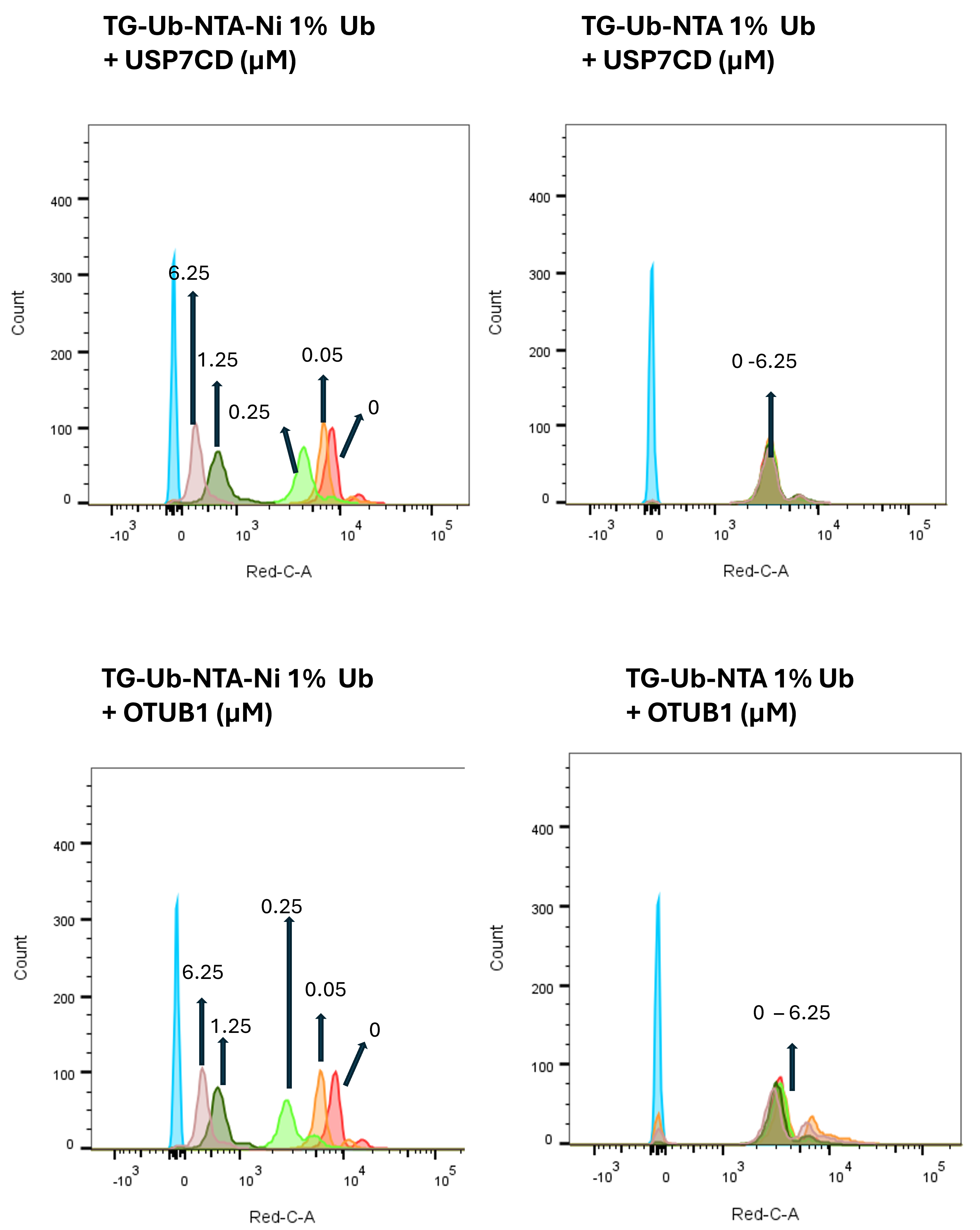

**Figure S2**: Flow cytometry analysis of Red C signal (Red C channel with emission wavelength of 670/14 nm) of beads (TG-Ub-NTA-Ni or TG-Ub-NTA) with 1% loading of ubiquitin after incubation with different concentrations of USP7 CD or OTUB1as indicated at room temperature for 6 h, stained with Alexa Fluor 647 labeled anti-ubiquitin.

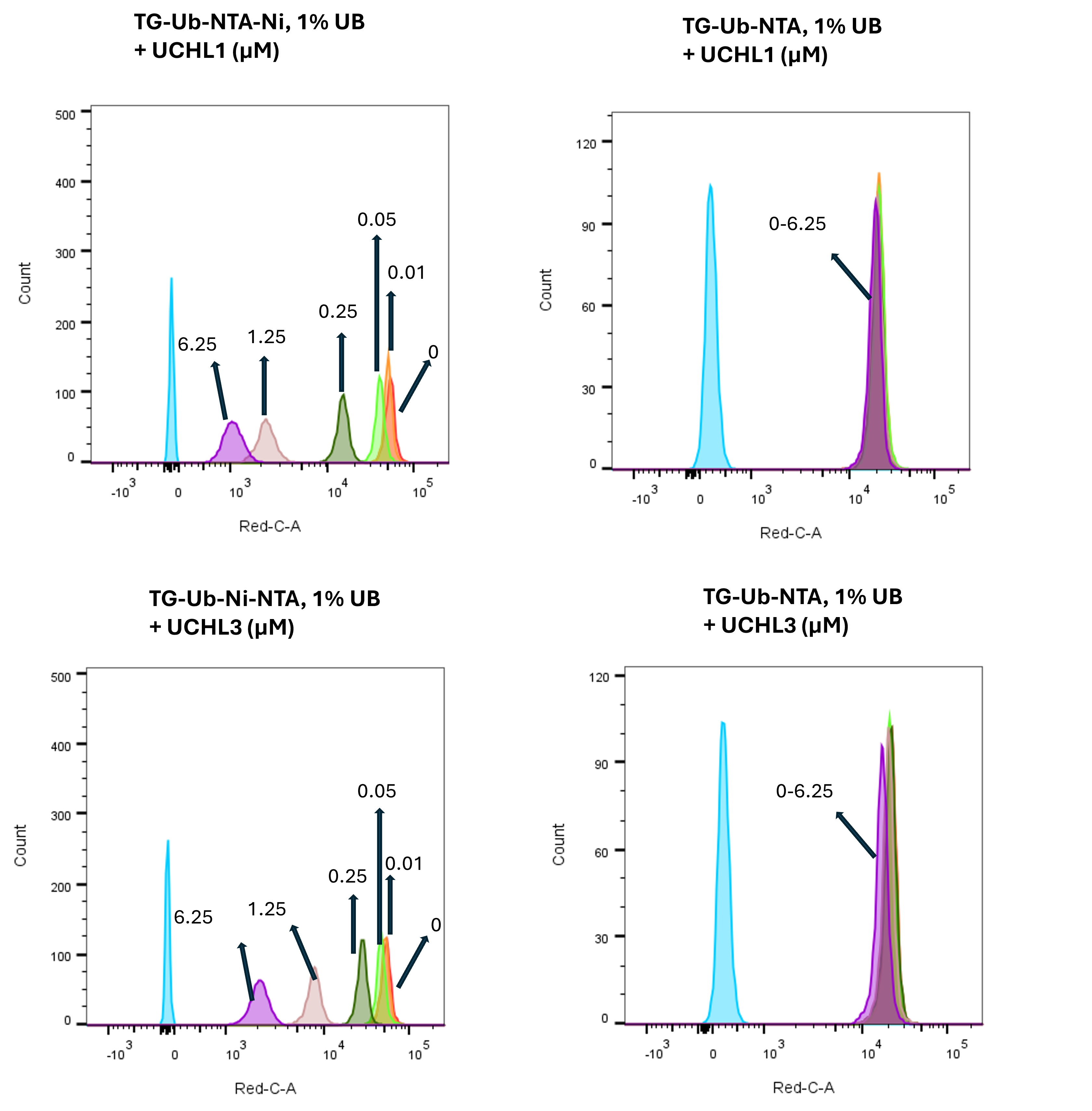

**Figure S3**: Flow cytometry analysis of Red C signal (Red C channel with emission wavelength of 670/14 nm) of beads (TG-Ub-NTA-Ni or TG-Ub-NTA) with 1% loading of ubiquitin after incubation with different concentrations of UCHL1 or UCHL3 as indicated at room temperature for 6 h, stained with Alexa Fluor 647 labeled anti-ubiquitin.

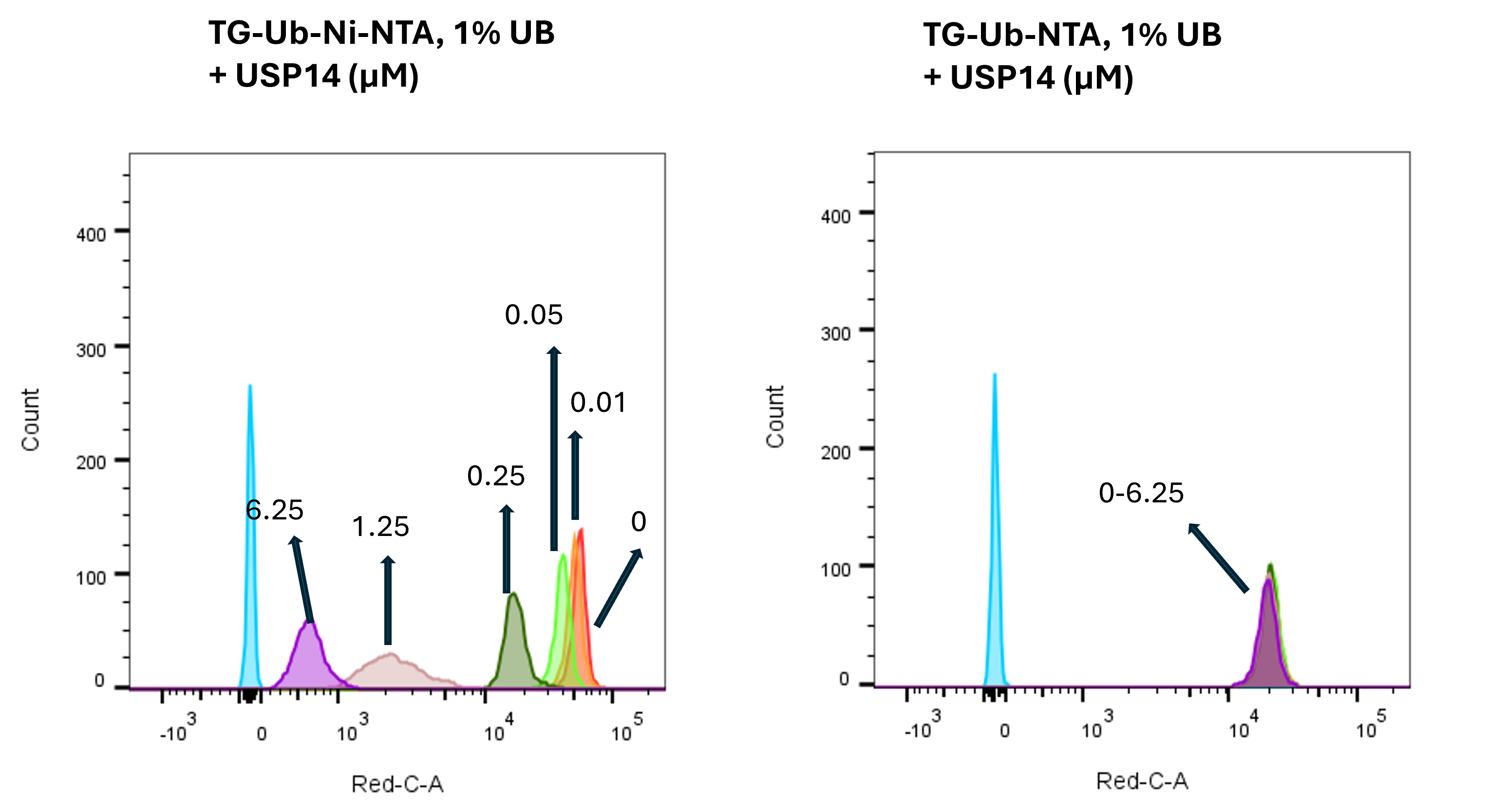

**Figure S4**: Flow cytometry analysis of Red C signal (Red C channel with emission wavelength of 670/14 nm) of beads (TG-Ub-NTA-Ni or TG-Ub-NTA) with 1% loading of ubiquitin after incubation with different concentrations of USP14 as indicated at room temperature for 6 h, stained with Alexa Fluor 647 labeled anti-ubiquitin.

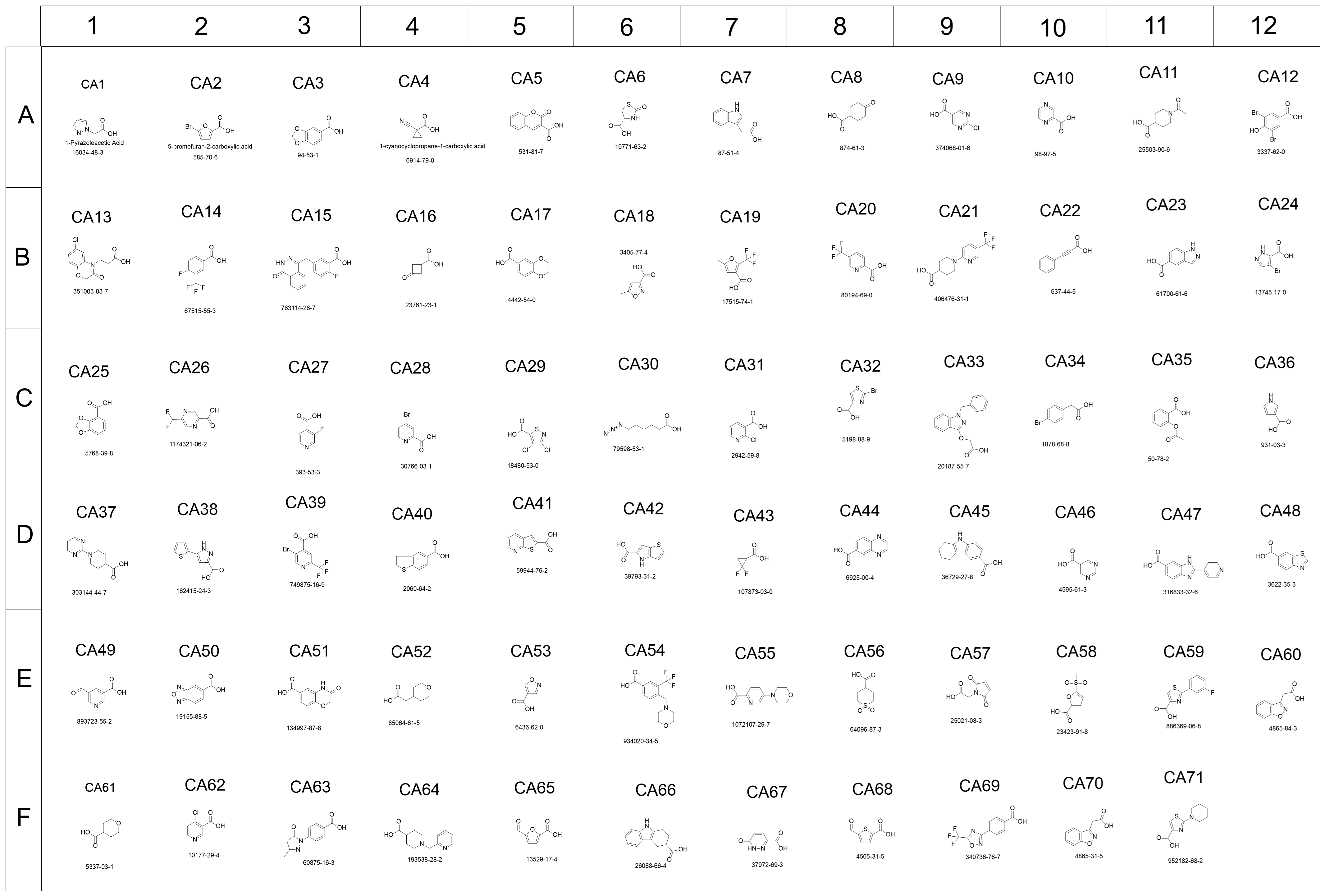

**Figure S5**: The chemical structure of 71 capping acids.

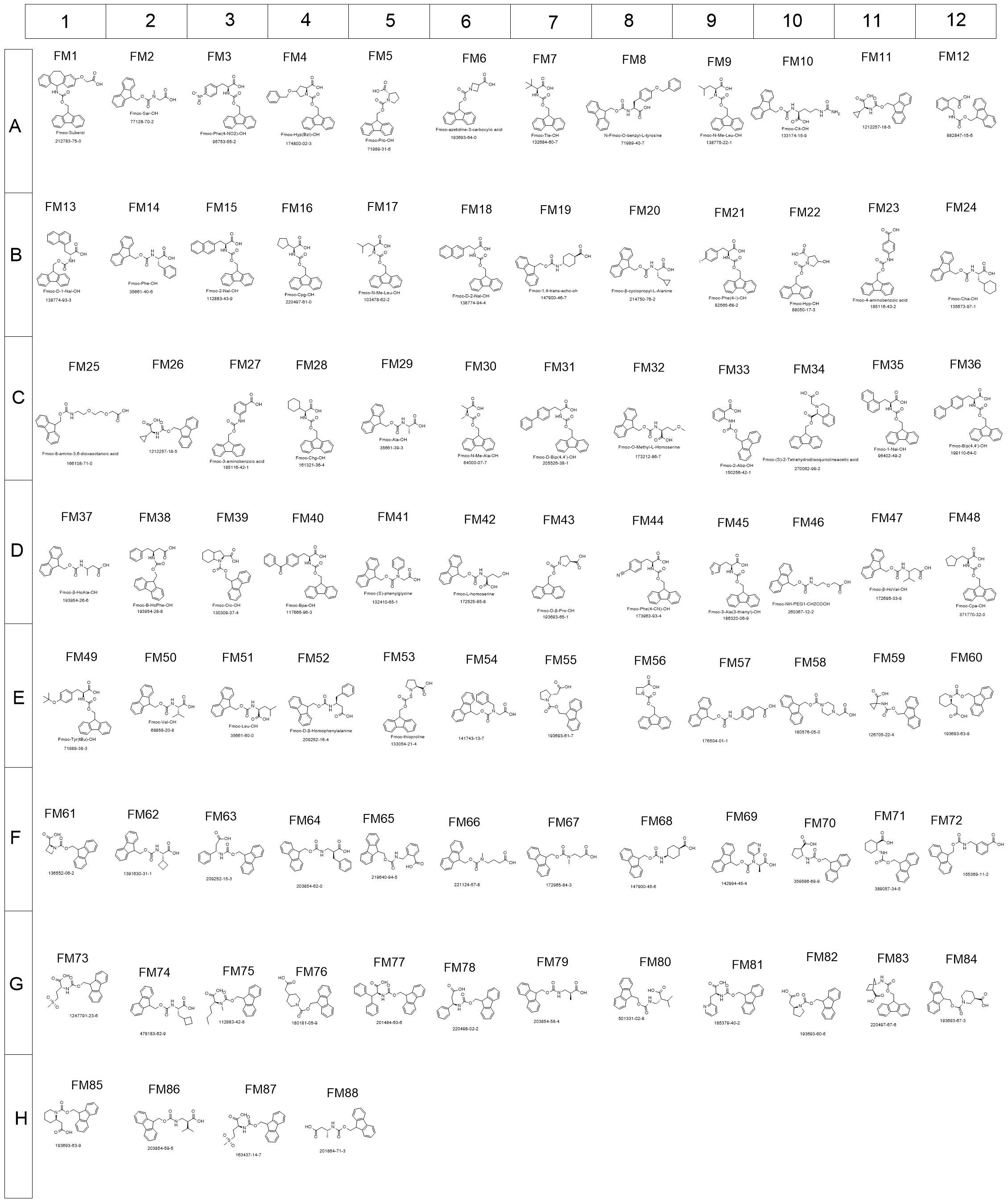

**Figure S6**: The chemical structure of 88 unnatural amino acids.

**General information.**

Chemical reagents were purchased from VWR, Millipore Sigma, ChemScene and Ambeed. ^1^H and ^13^C NMR spectra were obtained with Bruker AV400 NMR Spectrometer with a CryoProbe. Chemical shifts are reported in δ (ppm) units and ^13^C and ^1^H signals from deuterated solvents were used as references. 60 F254 (Merck) silica plate was used in thin layer chromatography (TLC) analysis.

ThermoFisher Scientific Vanquish Flex Binary 105 series HPLC equipped with ThermoFisher-C18 column (Part number: Acclaim 120, 3.0*100 mm. 3 µm) and an electrospray ionization (ESI) source (Thermo scientific ISQEM-EM) was used to obtain mass spectra for small molecules. The following methods were used to analyze the purity of the samples in the LC-MS instrument. Solvent A contains 100% water (0.1% formic acid) and solvent B contains 100% acetonitrile (0.1% formic acid).

| Time (min) | A [%] | B [%] | Flow [mL/Min] |
| --- | --- | --- | --- |
| 0.0 | 95.0 | 5.0 | 0.6 |
| 1.0 | 95.0 | 5.0 | 0.6 |
| 10.0 | 5.0 | 95.0 | 0.6 |
| 10.1 | 95.0 | 5.0 | 0.6 |
| 12.0 | 95.0 | 5.0 | 0.6 |

Small molecule column chromatography was conducted on silica gel (230-400 mesh) or a Biotage sfar Bio C18 column (300Å, 20 µm, 10 g, Biotage) or Biotage sfar Bio C18 column (300Å, 20 µm, 50 g, Biotage). The UV absorbance at 214, 254 nm were used to monitor the purification process. Water and acetonitrile were used as solvent A and B respectively. The following methods were used to purify the samples.

**General solid phase synthesis protocol:**

Compounds were synthesized on TentaGel resin using general Solid Phase Synthesis protocol described below. The compounds displayed on the beads were synthesized on 10 µm TentaGel M NH2 resin (Rapp-Polymere ,0.33 mmol/g) mixed with 10% of 300 µm TentaGel MB RAM resin (Rapp-Polymere, 0.19 mmol/g) for mass analysis. The beads were transferred to a fritted spin-column (Mobicol Classic, large filter, 10 µm pore size) and swelled in DMF (1 h, RT). FMOC was removed with 20% piperidine in DMF (3x resin volume after swelling, 500 µL) at room temperature for 15 minutes, twice. It was then washed thoroughly with DMF, EtOAc and DMF, and successively coupled with either amino acids or peptoid units. In general, acids (3 times of resin capacity, 1.0 mmol) were pre-activated at RT for 5 minutes with DIC/Oxyma/Collidine (1.4/1.0/1.0 mmol) in DMF, added to the resin, incubated at 37°C for one hour, and washed with DMF, EtOAc (5 times each, 5x resin volume). FMOC was removed where applicable.

**Supplemental Methods:**

**On-beads cleavage assay and flow** **cytometry analysis:** 10 µm TentaGel beads were stored in PBST (0.2 g/L KCl, 0.24 g/L KH_2_PO_4_, 8 g/L NaCl, 1.44 g/L Na_2_HPO_4_ anhydrous, 0.05% Tween-20, pH 7.4) after equipping with the chemicals indicated. Multiscreen® 96 well Plate equipped with hydrophilic PVDF membrane (MSBN12 from Millipore Sigma) was activated by EtOH and washed by PBST buffer. 10^5^ beads were transferred into each well of the plate and washed three times with PBST, incubated with 150 μL PBST at 4 °C for three hours, washed three times with PBST, and the beads were blocked by 150 μL StartingBlock (PBS) blocking buffer (ThermoFisher, Cat: 37538) overnight. Then, the beads were washed three times with deubiquitination buffer (20 mM Tris, 250 mM NaCl, 2 mM DTT, pH 8.0). Then the beads were incubated with the indicated concentrations of DUB in 100 μL deubiquitination buffer at room temperature for 6 hours. After incubation, the beads were washed with PBST for 6 times, and incubated with 1 μL Alexa Fluor® 647 anti-Ubiquitin Antibody (Biolegend, Cat: 838710, 0.5 mg/mL) in 150 μL PBST buffer in each well over night at 4 °C with shaking. Then the beads were washed with PBST for 6 times and analyzed by flow cytometry (BD Fortessa). The Red C channel was selected for the analysis. The signals were plotted using FlowJo_V10.10.0.

**DUBs expression and purification**

His tag labeled USP7 full length (Cat: 227214), USP7 catalytic domain (Cat: 227201), OTUB1 (Cat: 25149) plasmids were purchased from addgene. His tag labeled USP14, UCHL1, UCHL3, USP7 TRAF plasmids were purchased from GeneScript and transformed into BL21 cells and cultured in Terrific Broth at 37 °C until OD600 reached 0.3-0.4. The incubation temperature decreased to 22 °C until OD600 reached 0.6. Then, protein expression was induced with 1 mM IPTG. Cells were cultured for an additional 24 h at 22 °C after the induction and harvested by centrifugation at 5,000 rpm for 15 min at 4 °C. For 6 L of cell culture, the cells were lysis by 200 mL lysis buffer (50 mM sodium phosphate; 300 mM NaCl, 1.5 mM imidazole, pH 8.0). Then the lysate was centrifuged at 3000 × g for 30 minutes at 4 °C to pellet the cellular debris. Transfer the supernatant to a fresh tube and incubate with 8 mL HisPur Ni-NTA resin (Thermo Scientific, 88221) for 2 hours at 4°C with shaking. Then the HisPur Ni-NTA resin was washed with 200 mL wash buffer (50 mM sodium phosphate; 300 mM NaCl; 1.5 mM imidazole, pH 8.0). Then Gradient elution of the protein by 5 mL 100 mM imidazole in Wash Buffer, 5 mL 250 mM imidazole in Wash Buffer, 5 mL 500 mM imidazole in Wash Buffer, 5 mL 1000 mM imidazole in Wash Buffer. Collect each elution separately. The desire protein product was buffer exchanged using a PD-10 column (Cytiva, 17085101) using deubiquitinating buffer (20 mM Tris, 250 mM NaCl, 2 mM DTT, pH 8.0).

**His-UCHL1 amino acids sequence:**

MGSSHHHHHHSSGLVPRGS

MQLKPMEINPEMLNKVLSRLGVAGQWRFVDVLGLEEESLGSVPAPACALLLLFPLTAQHENFRKKQIEELKGQEVSPKVYFMKQTIGNSCGTIGLIHAVANNQDKLGFEDGSVLKQFLSETEKMSPEDRAKCFEKNEAIQAAHDAVAQEGQCRVDDKVNFHFILFNNVDGHLYELDGRMPFPVNHGASSEDTLLKDAAKVCREFTEREQGEVRFSAVALCKAA

**His-UCHL3 amino acids sequence:**

MGSSHHHHHHSSGLVPRGS

MEGQRWLPLEANPEVTNQFLKQLGLHPNWQFVDVYGMDPELLSMVPRPVCAVLLLFPITEKYEVFRTEEEEKIKSQGQDVTSSVYFMKQTISNACGTIGLIHAIANNKDKMHFESGSTLKKFLEESVSMSPEERARYLENYDAIRVTHETSAHEGQTEAPSIDEKVDLHFIALVHVDGHLYELDGRKPFPINHGETSDETLLEDAIEVCKKFMERDPDELRFNAIALSAA

**His-USP14 amino acids sequence:**

MGSSHHHHHHSSGLVPRGS

MPLYSVTVKWGKEKFEGVELNTDEPPMVFKAQLFALTGVQPARQKVMVKGGTLKDDDWGNIKIKNGMTLLMMGSADALPEEPSAKTVFVEDMTEEQLASAMELPCGLTNLGNTCYMNATVQCIRSVPELKDALKRYAGALRASGEMASAQYITAALRDLFDSMDKTSSSIPPIILLQFLHMAFPQFAEKGEQGQYLQQDANECWIQMMRVLQQKLEAIEDDSVKETDSSSASAATPSKKKSLIDQFFGVEFETTMKCTESEEEEVTKGKENQLQLSCFINQEVKYLFTGLKLRLQEEITKQSPTLQRNALYIKSSKISRLPAYLTIQMVRFFYKEKESVNAKVLKDVKFPLMLDMYELCTPELQEKMVSFRSKFKDLEDKKVNQQPNTSDKKSSPQKEVKYEPFSFADDIGSNNCGYYDLQAVLTHQGRSSSSGHYVSWVKRKQDEWIKFDDDKVSIVTPEDILRLSGGGDWHIAYVLLYGPRRVEIMEEESEQ

**His-USP7 TRAF (1-205) amino acids sequence:**

MHHHHHHSSGLVPRGS

MNHQQQQQQQKAGEQQLSEPEDMEMEAGDTDDPPRITQNPVINGNVALSDGHNTAEEDMEDDTSWRSEATFQFTVERFSRLSESVLSPPCFVRNLPWKIMVMPRFYPDRPHQKSVGFFLQCNAESDSTSWSCHAQAVLKIINYRDDEKSFSRRISHLFFHKENDWGFSNFMAWSEVTDPEKGFIDDDKVTFEVFVQADAPHGVAW

**Mammalian Cell Culture**. The commercially available human cystic fibrosis bronchial epithelial CFBE41o- 4.7 ΔF508-CFTR cells were obtained from Millipore Sigma (SCC159). CFBE41o- 4.7 ΔF508-CFTR cells were derived from parental CFBE41o- cells by introducing the ΔF508-CFTR expressing construct. CFBE41o- 4.7 ΔF508-CFTR cells were cultured in α-MEM media (Sigma. #M2279) supplemented with 10% fetal bovine serum (FBS), 2 mM-glutamine, 300 μg/mL hygromycin B, 100 U/mL penicillin, and 100 μg/mL streptomycin, 37 °C, 5% CO_2_.

**Immunoblotting Analysis of CFTR protein level**. For treatment with indicated compounds, the CFBE41o- 4.7 ΔF508-CFTR cells were plated in a 6-well plate precoated with a coating mixture 10 μg/mL Fibronectin (Sigma, #F2006), 30 μg/mL PureCol Collagen (Sigma, #5006), and 100 μg/mL BSA (Sigma. #126575)), and treated with the indicated compounds when the cells reached 60∼70% confluency. Post compound treatment, cells were washed 2x with DPBS (14190250, Thermo Fisher Scientific), and harvested by TrypLE™ Express Enzyme (1X), phenol red (12605010, Thermo Fisher Scientific). Harvested cells were used immediately for the following experiments or stored at -80°C. Cells were lysed with 20 μL Mammalian Permeabilization Reagent (MPER) (78501, Thermo Fisher Scientific) and incubated on ice for 30 minutes. Soluble protein supernatant was clarified by centrifugation at 12,000g for 10 minutes at 4 °C in a benchtop microcentrifuge (Eppendorf). Protein concentration was determined by protein A280 by Nanodrop. 40 μg cell lysate was loaded onto 4-15% gradient gels (Cat#4561083, BioRad). SDS-Page gels were run at 130V for 60 minutes, and transferring to nitrocellulose membrane was executed according to manufacturer’s instructions (1704270, Bio-Rad). Membranes were blocked in 5% Blot-Quickblocker modified milk protein (786-011, G-Biosciences) in Tris buffered saline containing 0.1% Tween-20 (TBST) buffer at room temperature for 1h. Then, the membranes were incubated with primary antibodies CFTR (#78335, 1:1,000, Cell Signaling Technologies) or anti-GAPDH (60004-1-1g, proteintech, 1:100,000) at 4 °C overnight. After washing five times with TBST, the membranes were further incubated with indicated secondary antibodies: anti-mouse secondary antibody (A4416, 1:3000, Sigma) or anti-rabbit secondary antibody (A-4914, 1:3000, Sigma) for another 1 h at room temperature. Then the blots were stained with 10 mL Immobilon Western Chemiluminescent HRP Substrate (WBKLS0050, Sigma) for 10 min. Then, the blots were imaged by Bio-Rad ChemiDoc Touch Imaging System. Western blots were quantified using ImageJ (RRID: SCR_003070). The quantification of CFTR level is normalized to DMSO control.

**Fluorescence Polarization Assay**. A serial dilution of recombinant USP7 TRAF domain in 20 mM Tris, 250 mM NaCl, 2 mM DTT, pH 8.0. was prepared and incubated with 100 nM fluorescent probe at room temperature for 60 minutes with slow shaking.

The fluorescent polarization signal was obtained in Molecular Devices FilterMax F5 plate reader with excitation at wavelength 485 nm (bandwidth 10 nm) and detection of emitted light at wavelength 528 nm (bandwidth 10 nm).

**Ubiquitin-Rhodamine Activity Assay**. DUB activity was tested with a fluorogenic, ubiquitin-rhodamine 110 (UbRho) assay in black 96 well plate. Different concentrations of enzymes were incubated with 1 µM UbRho in buffer 20 mM Tris, 250 mM NaCl, 2 mM DTT, pH 8.0. The fluorescent intensity was obtained in Molecular Devices FilterMax F5 plate reader at excitation: 485 nm and emission: 535 nm.

**Scheme S1: Ubiquitin 1-76-Propargylamide generation**

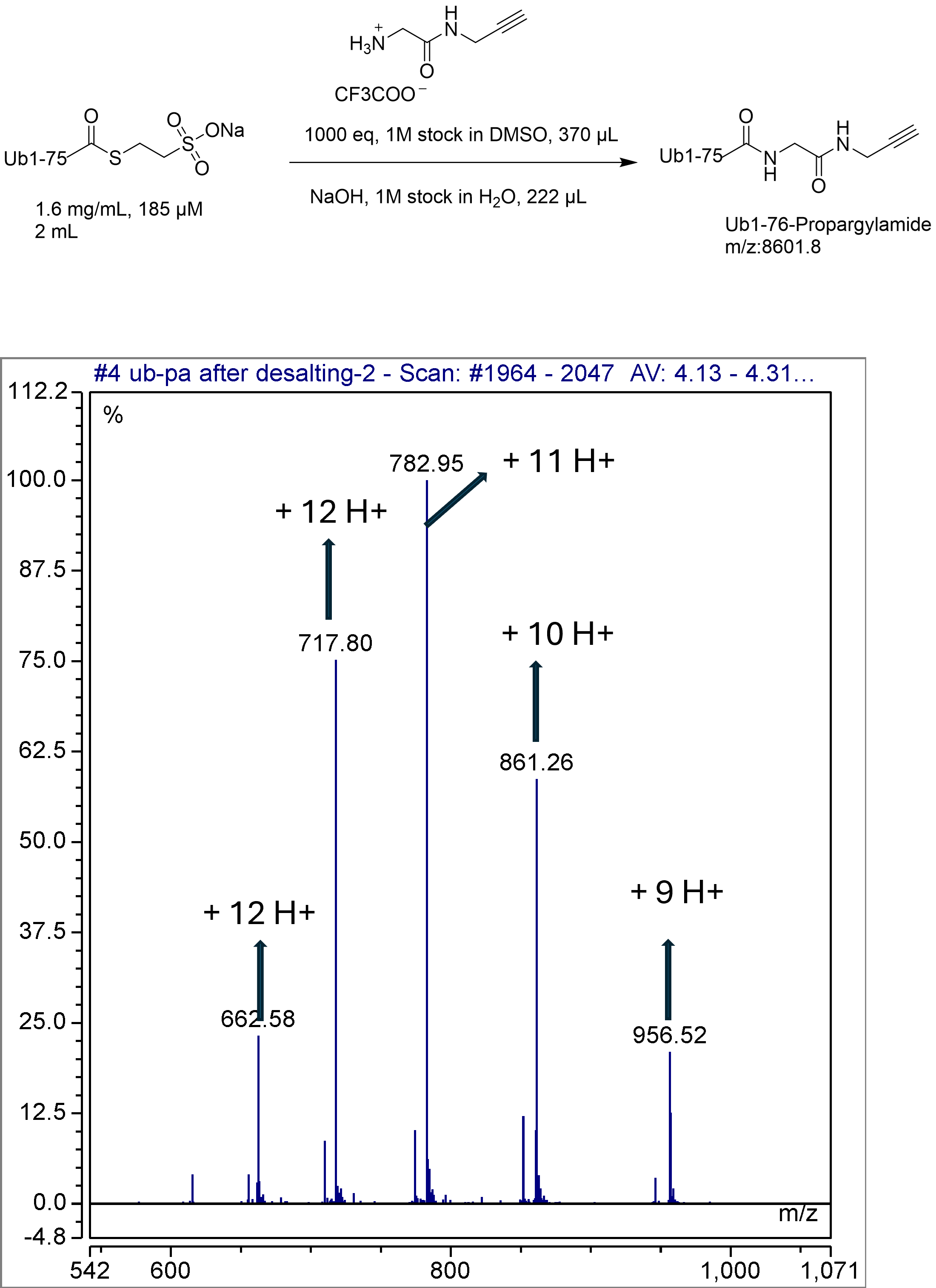

Scheme S1: Generation of TG-Ub-NTA-Ni with different loading of ubiquitin on 10 μm TentaGel beads.

Ubiquitin 1-75 Mesna was generated as previously reported.^1^ Gly-Propargylamide (1000 eq, 1M in DMSO, 370 µL) and NaOH (600 eq, 1M in H_2_O, 222 µL) were mix together and Ubiquitin1-75 Mesna (1.6 mg/mL, 2 mL) was added to the solution and incubate at 37 °C for 16 h. Then, the product Ub1-76-Propargylamide was concentrated with Amicon Ultra column with a 3K centrifugal filter. Then the product was buffer exchanged by PD-10 column (Cytiva 17085101) using 1M GuHCl buffer. And characterized by LC-MS. The calculated mass is 8601.8. LC-MS detected multiple charge states as indicated above.

**Scheme S2: Generation of TG-Ub-NTA-Ni with different loadings of ubiquitin on 10 μm TentaGel beads.**

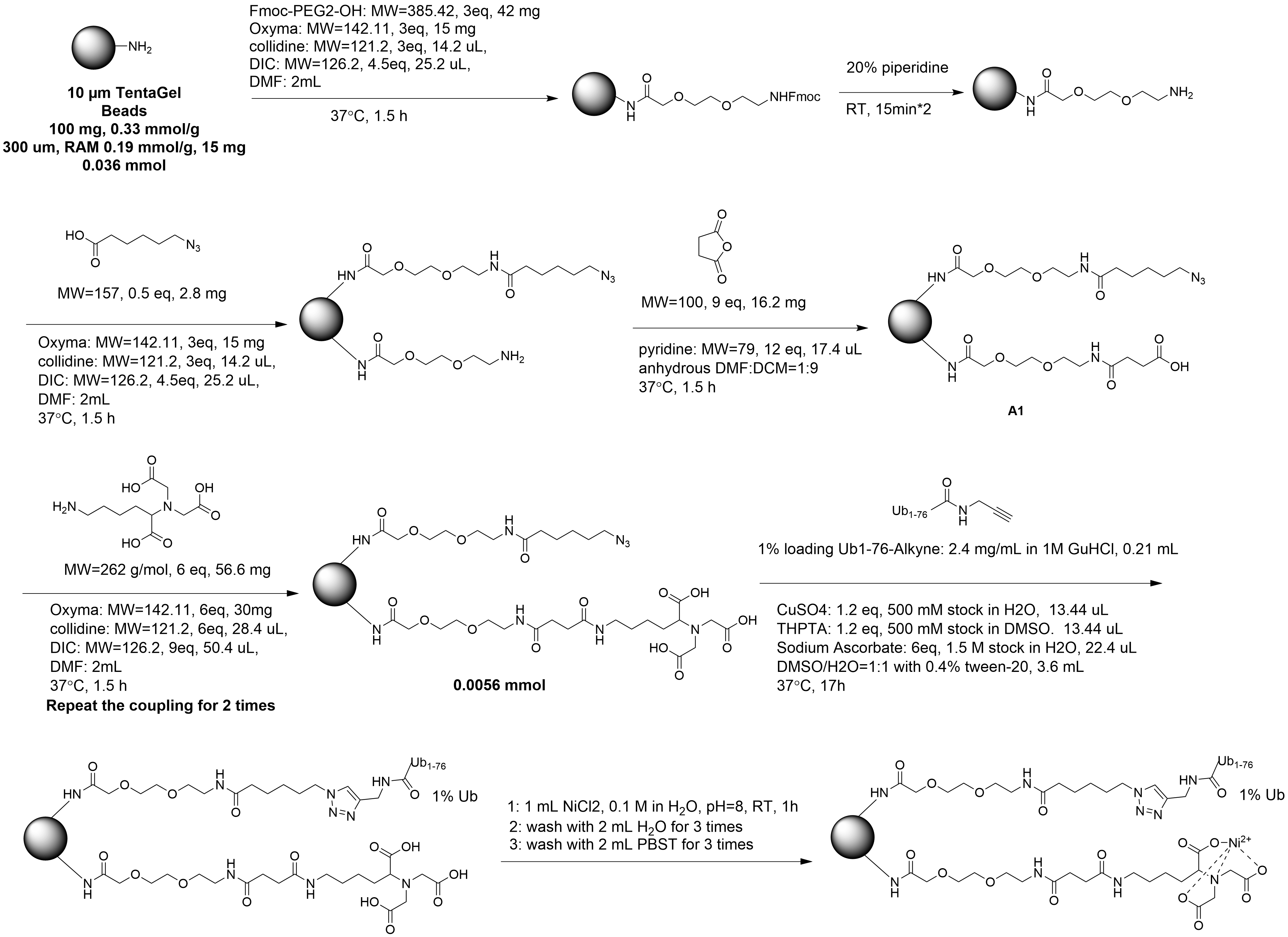

Intermediate **A1** was prepared according to the standard TentaGel solid-phase synthesis protocol described above. To install the nitrilotriacetic acid (NTA) functionality, Oxyma, collidine, and *N,N′*-diisopropylcarbodiimide (DIC) were dissolved in 2 mL of DMF and added to **A1** beads to activate the COOH on the resin for 10 min at room temperature, nitrilotriacetic acid was added, and the reaction mixture was incubated at 37 °C for 1.5 h. The beads were then washed sequentially with EtOAc, MeOH, and DMF. This coupling procedure was repeated twice to ensure complete NTA functionalization.

Ubiquitin loading was controlled by varying the ratio of ubiquitin(1–76)-Propargylamide. Specifically, 0.0056 mmol of NTA-functionalized beads was reacted with mixtures containing ubiquitin(1–76)- Propargylamide at defined molar ratios using a Cu(I)-catalyzed azide–alkyne cycloaddition (CuAAC). The resulting beads contained different surface loadings of ubiquitin. Then the beads were washed with BTPBB buffer (100 mM NaCl, 10 mM EDTA, 1% SDS, 1% Tween 20, 10 mM Bis-Tris, pH=7.6) for three times. And incubate in BTPBB buffer for 3 hours at room temperature and wash with BTPBB buffer again. Then, the beads were washed with Milli-Q water for 3 times.

Then the beads were charged with Ni²⁺ by incubation in 1 mL of 0.1 M NiCl₂ (Milli-Q water, pH 8.0) at room temperature for 1 h. The beads were subsequently washed three times with 2 mL of Milli-Q water, followed by three washes with PBST buffer, and stored in PBST until use.

**Scheme S3: Generation of Tentagel beads displaying USP7 ligands.**

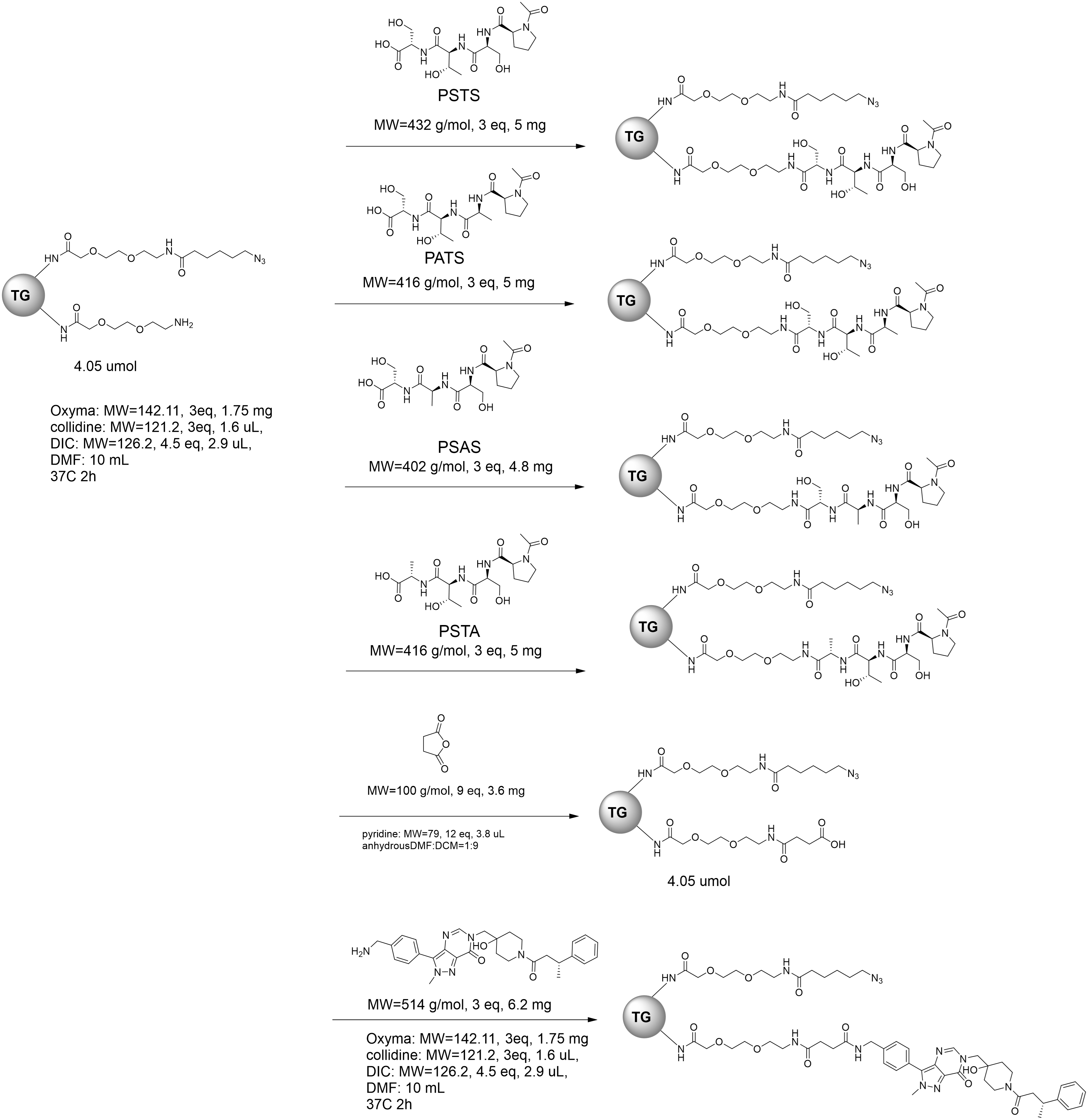

Generation of the beads displaying different USP7 ligands follows the standard TentaGel solid-phase synthesis protocol described above. PSTS, PATS, PSAS, PSTA were purchased from Genescript. The USP7 inhibitor USP-IN-2 was purchased from Sigma. The LC-MS quality control of the USP7-IN-2 beads is shown below.

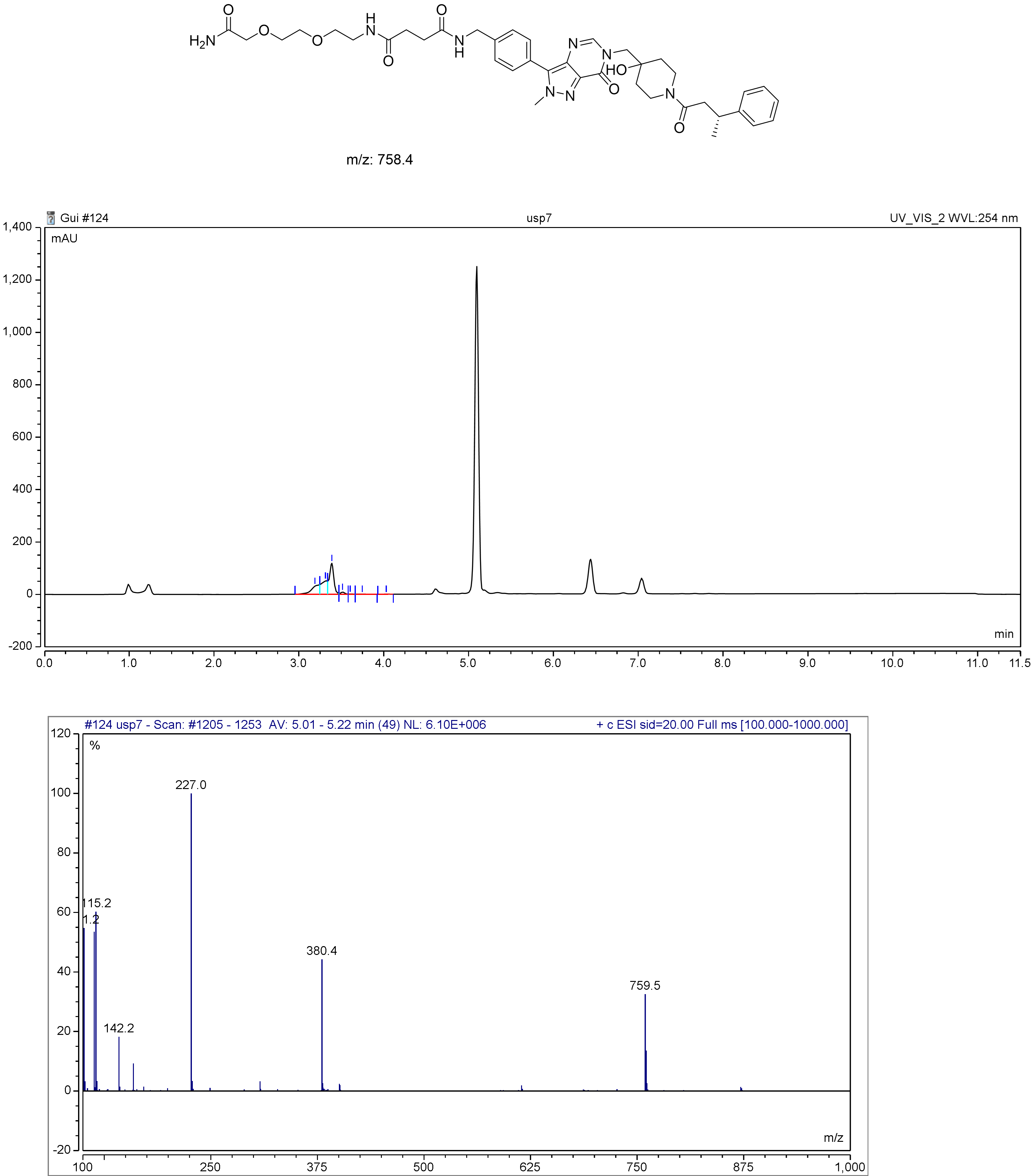

**Scheme S4: Generation of fluorescent probe WJ003.**

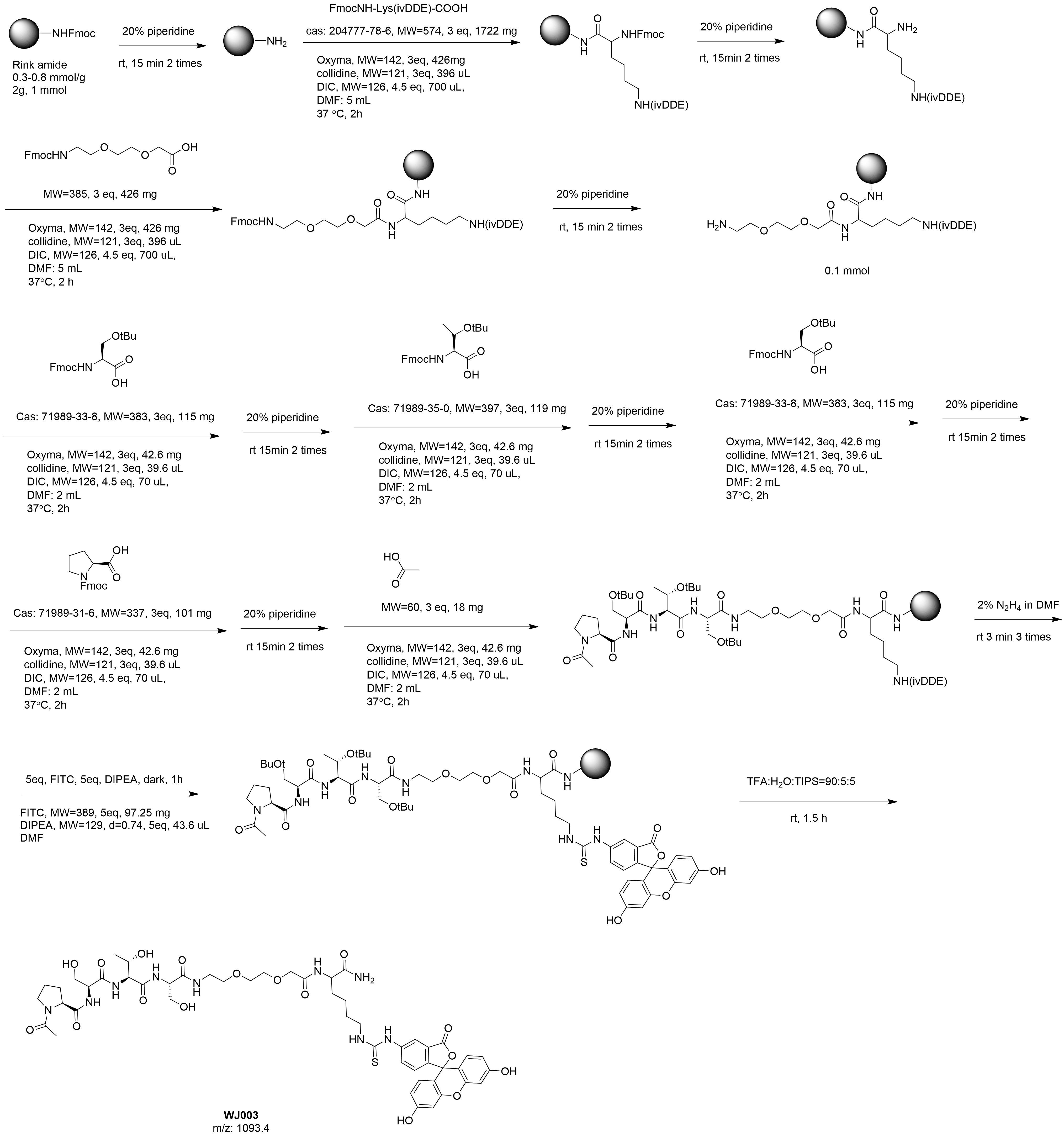

Fluorescent probe **WJ003** was synthesized using standard Fmoc-based solid-phase peptide synthesis (SPPS) on Rink amide resin. Upon completion of the synthesis, the product was cleaved from the resin using a cleavage cocktail consisting of trifluoroacetic acid (TFA)/H₂O/triisopropylsilane (TIPS) (90:5:5, v/v/v). The cleavage solution was concentrated under reduced pressure, and the crude product was purified by reverse-phase C18 preparative HPLC. The purified probe was characterized by LC–MS.

The generation of fluorescent probe **WJ004**, **WJ011** and **WJ012** follow the same protocol as described above. The LC-MS characterization is shown below.

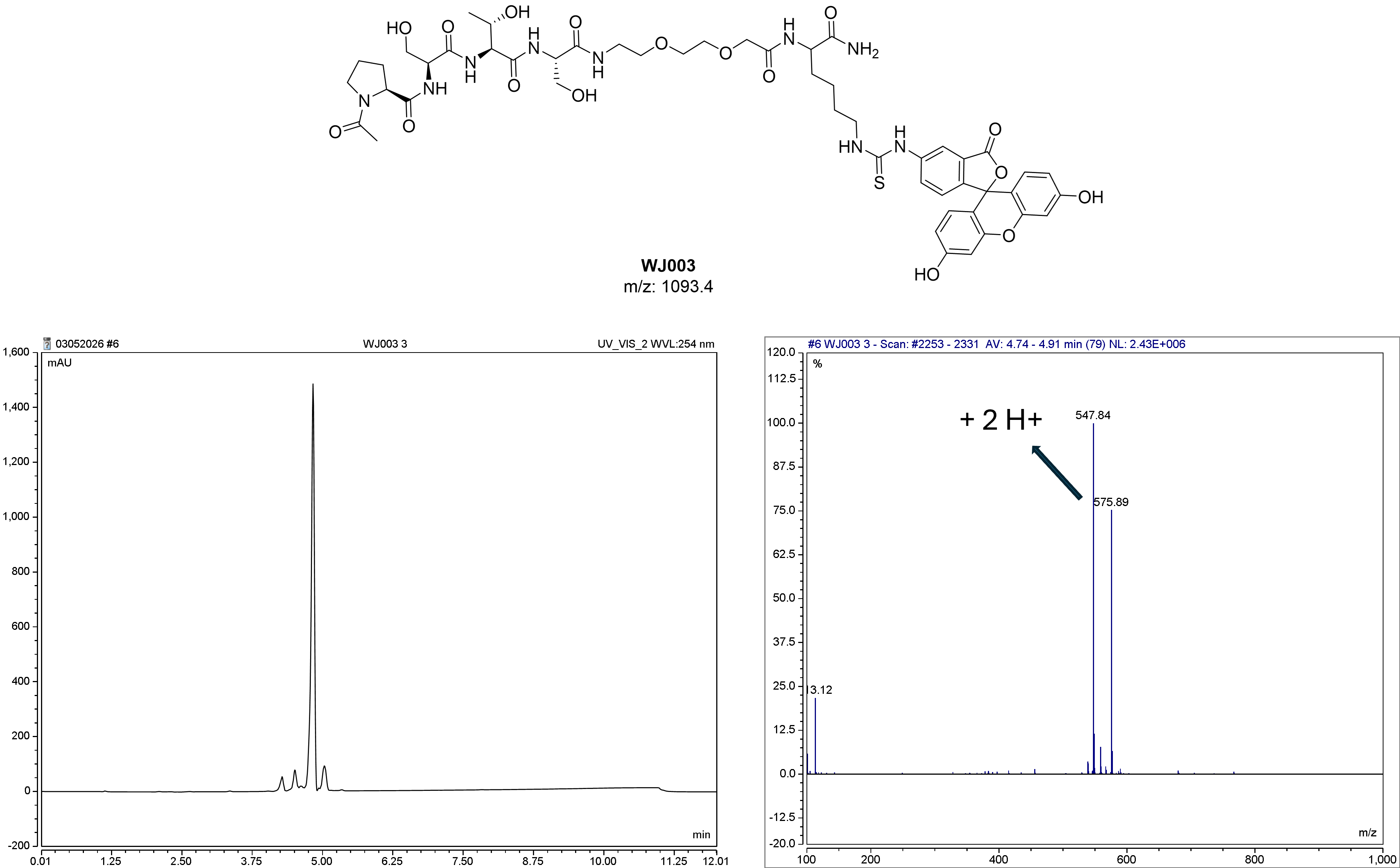

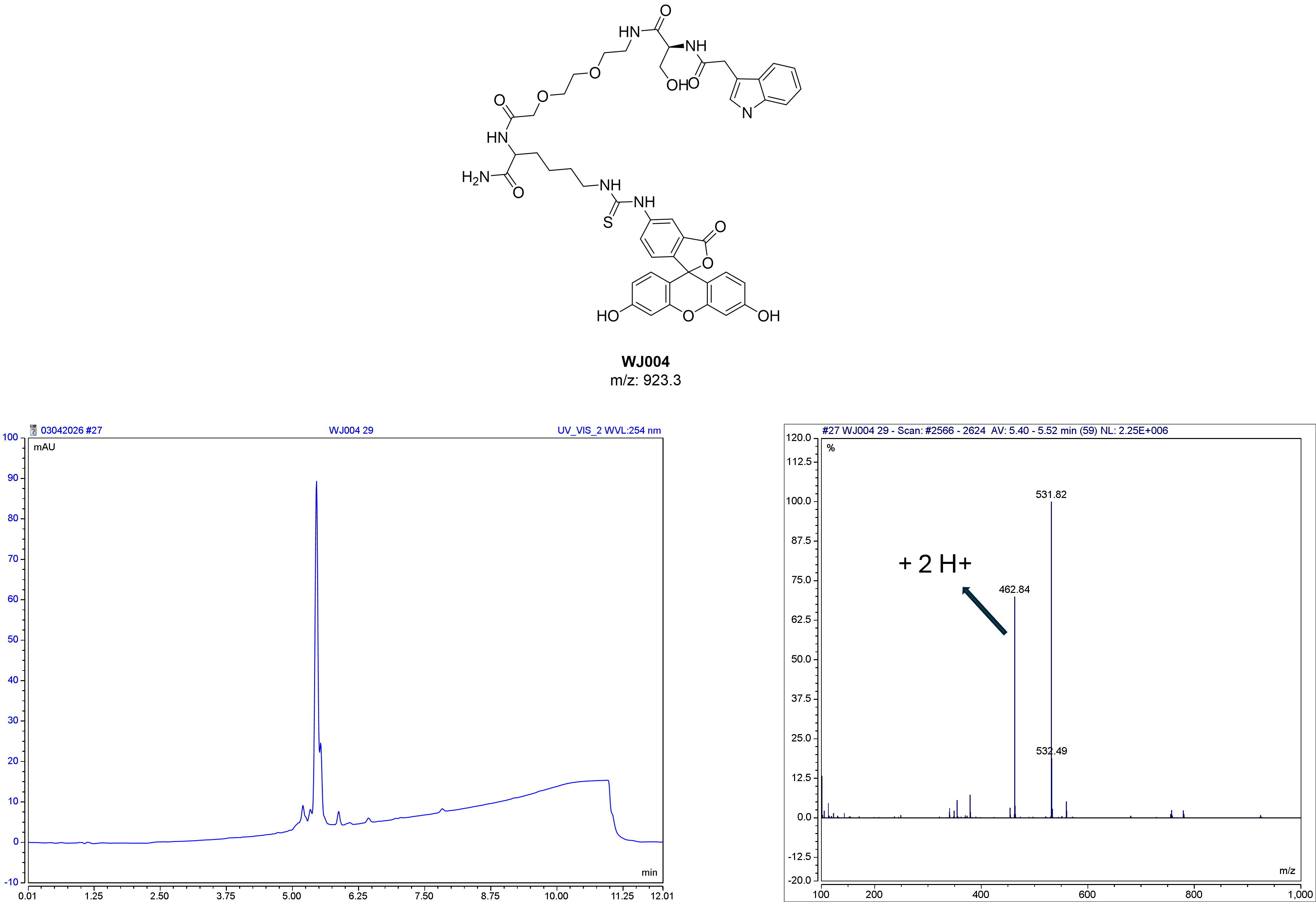

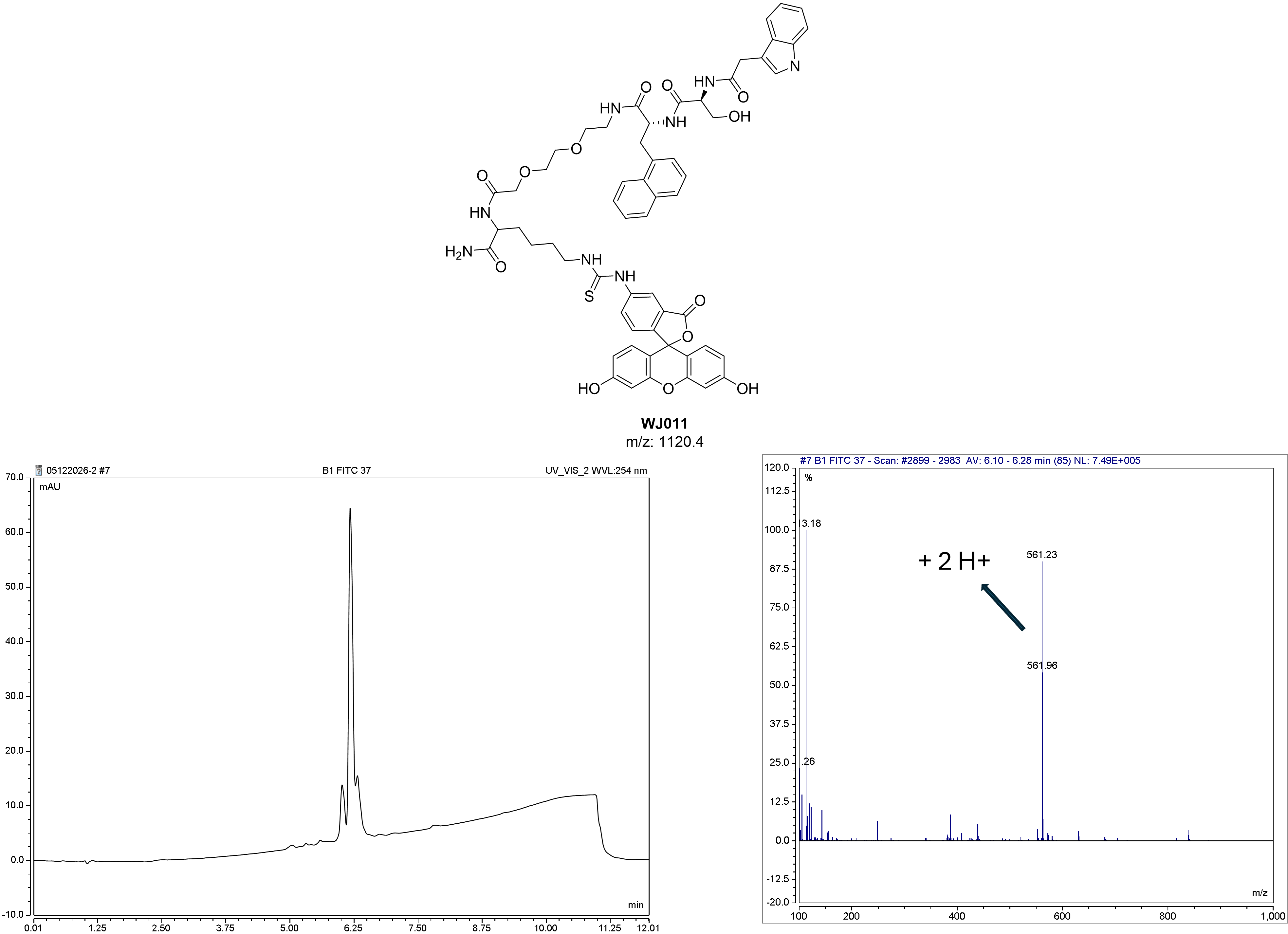

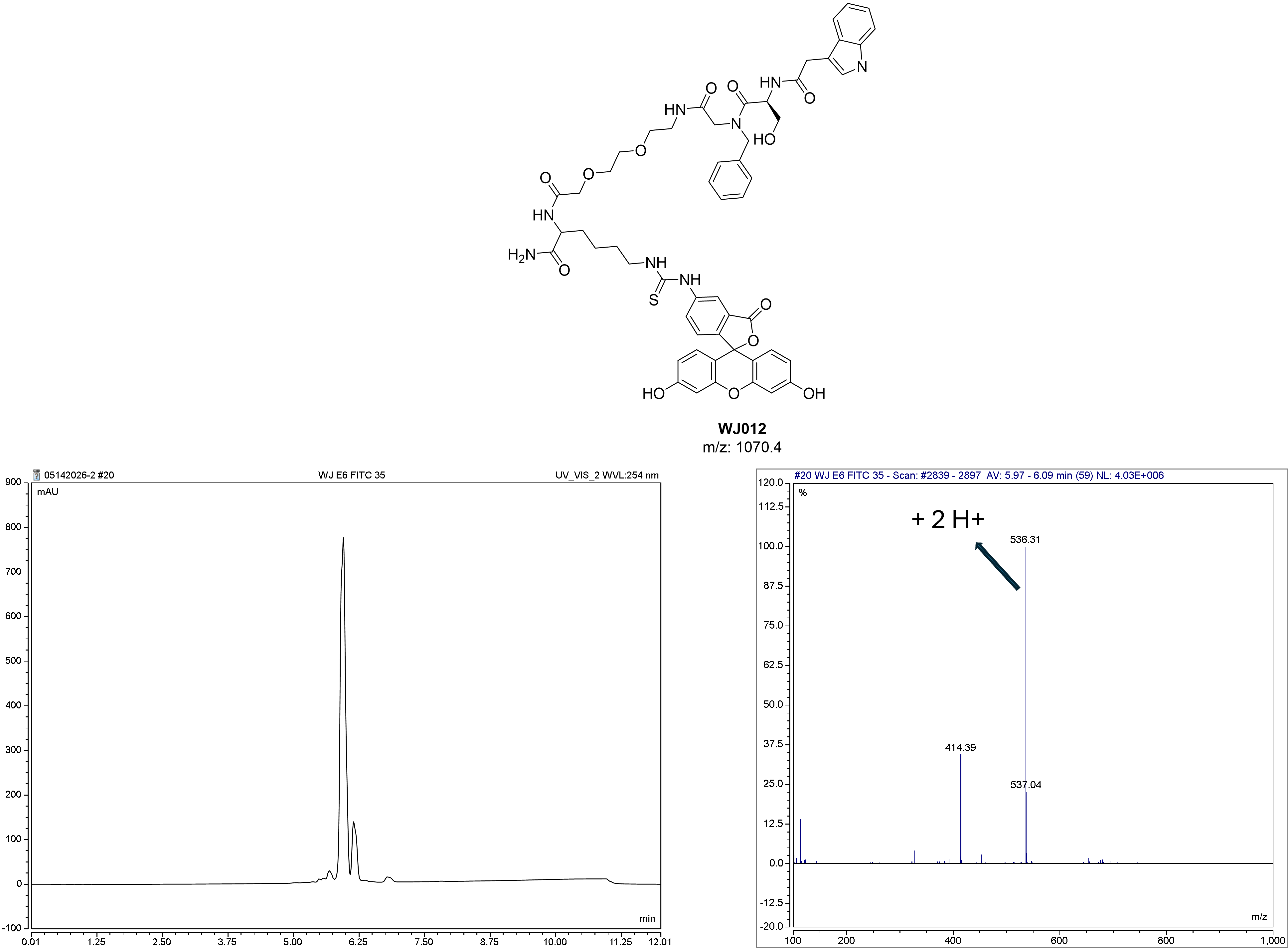

**Scheme S5: Generation of WJ021.**

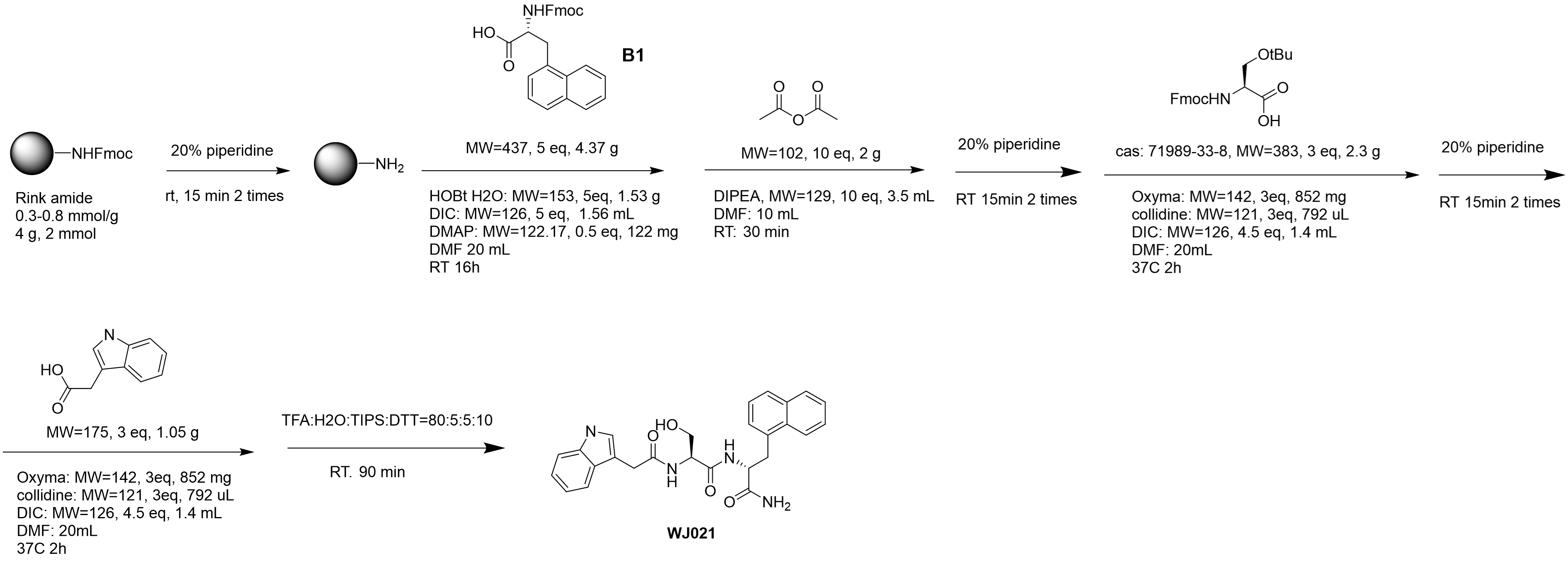

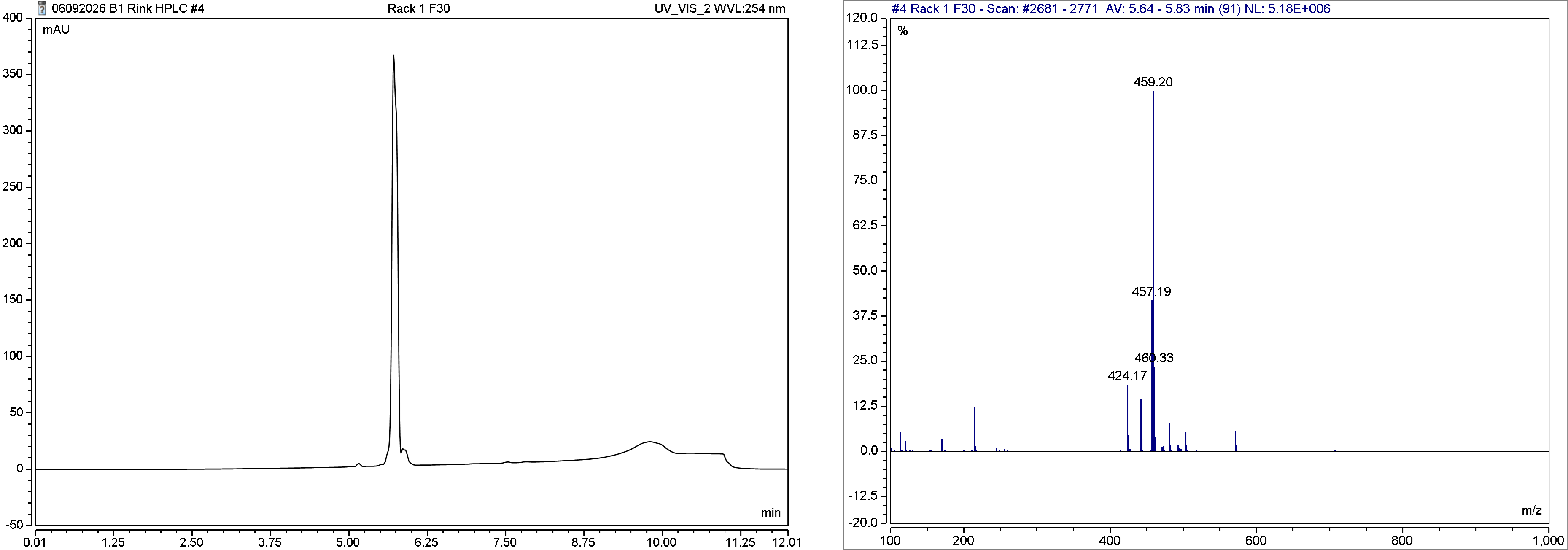

The synthesis of WJ021 follows a standard Rink amide solid phase synthesis. The final product was purified by HPLC and characterized by LC-MS. The calculated mass of WJ021 is 458.20, the mass found is 459.20.

**WJ021**

^1^H NMR (400 MHz, DMSO) δ 10.85 (s, 1H), 8.21 (dd, *J* = 14.0, 8.4 Hz, 2H), 7.99 (d, *J* = 7.0 Hz, 1H), 7.90 (d, *J* = 7.9 Hz, 1H), 7.76 (d, *J* = 7.9 Hz, 1H), 7.60 – 7.47 (m, 3H), 7.37 – 7.26 (m, 4H), 7.20 (d, *J* = 2.3 Hz, 2H), 7.09 – 7.02 (m, 1H), 7.00 – 6.93 (m, 1H), 4.86 (t, *J* = 5.4 Hz, 1H), 4.45 (td, *J* = 9.0, 4.3 Hz, 1H), 4.18 (q, *J* = 6.1 Hz, 1H), 3.68 – 3.55 (m, 3H), 3.44 – 3.35 (m, 2H), 3.11 (dd, *J* = 14.2, 9.7 Hz, 1H).

^13^C NMR (101 MHz, DMSO) δ 173.44, 171.56, 170.62, 136.54, 133.86, 131.97, 129.06, 127.41, 126.56, 125.77, 124.22, 124.05, 121.39, 119.18, 118.77, 111.73, 109.16, 61.95, 56.17, 53.94, 34.82, 32.82.

MS (ESI, positive) m/z calculated for C_26_H_27_N_4_O_4_ [M+H]^+^ : 459.20, found: 459.20.

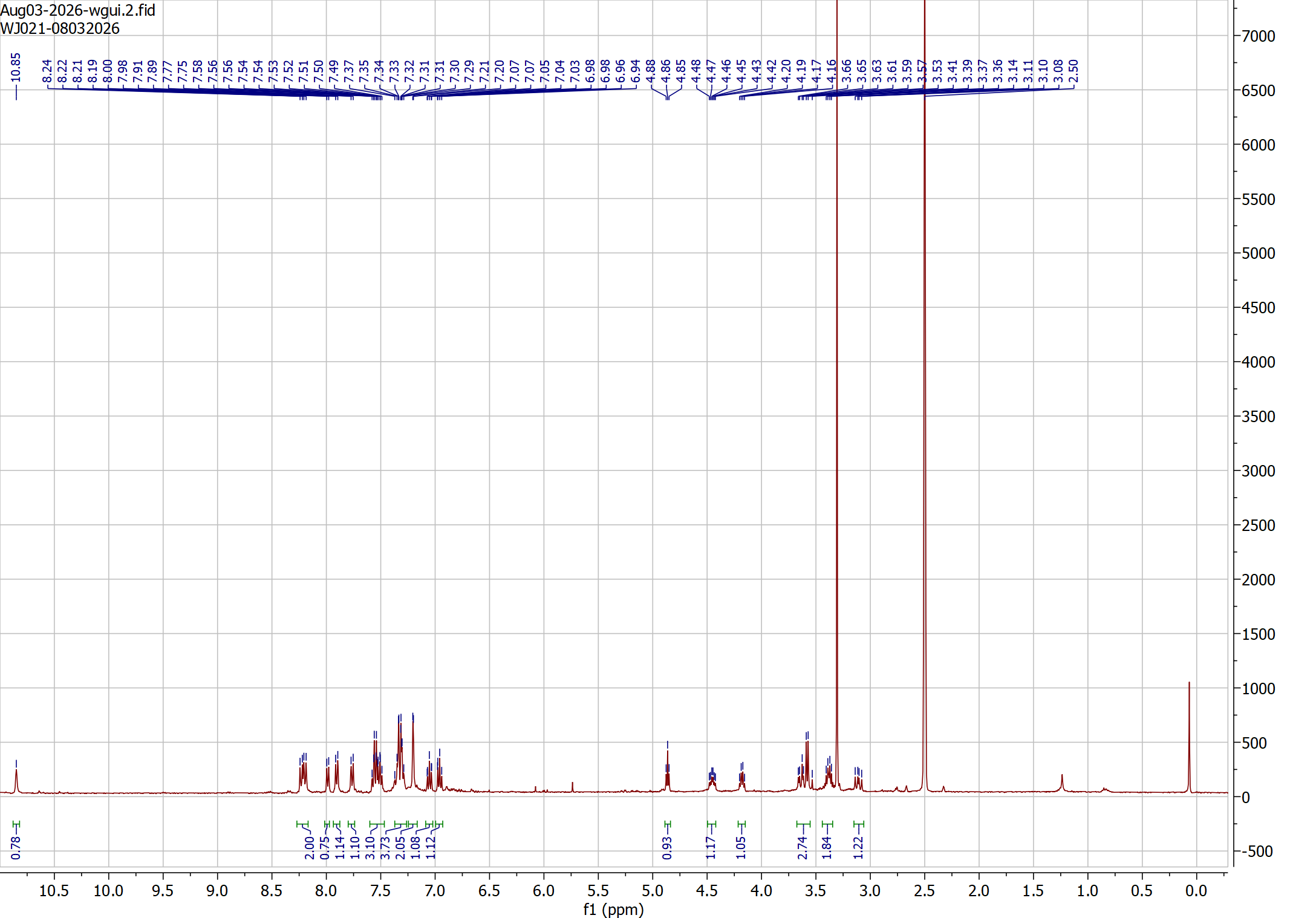

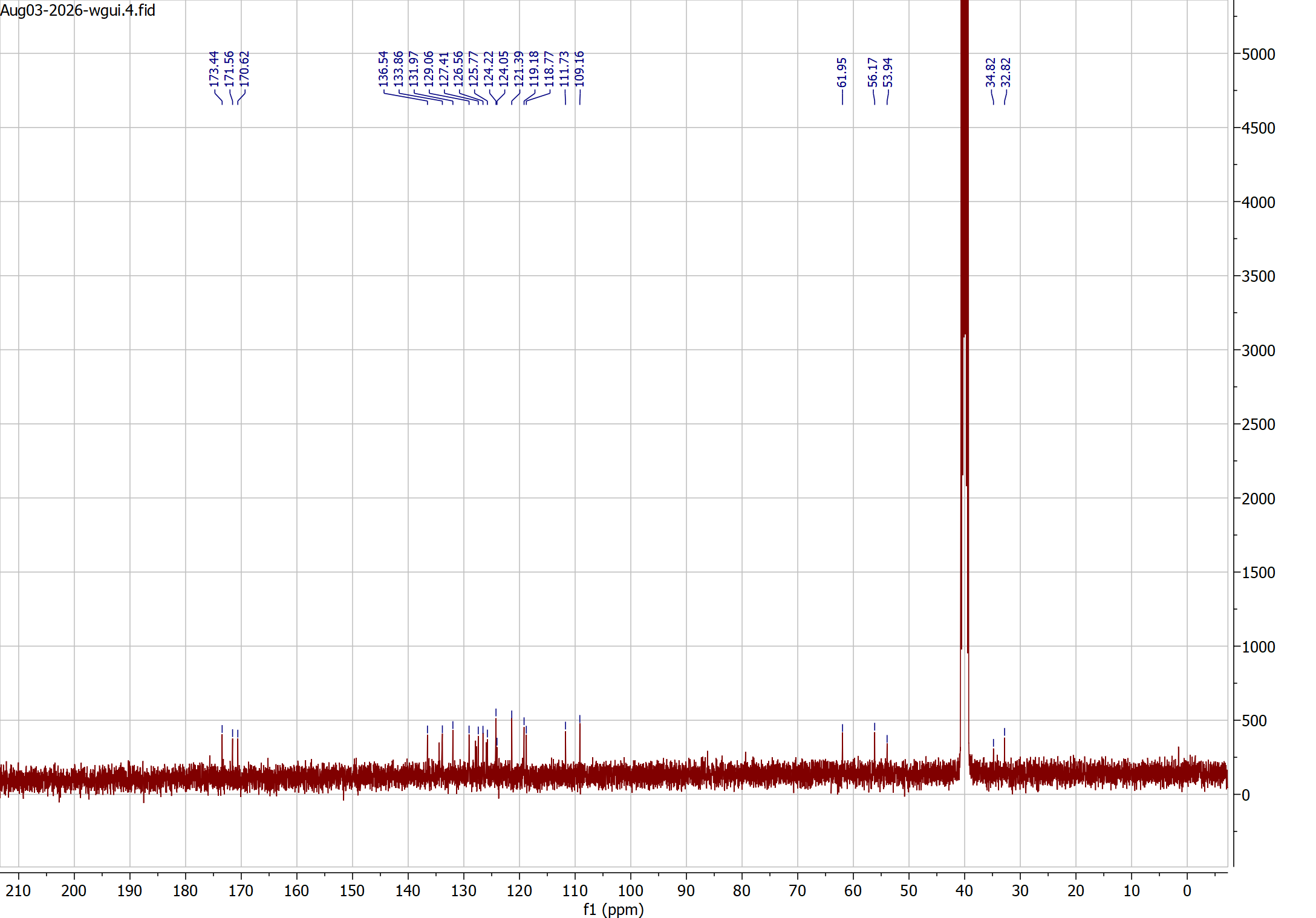

**Scheme S6: Generation of WJ020.**

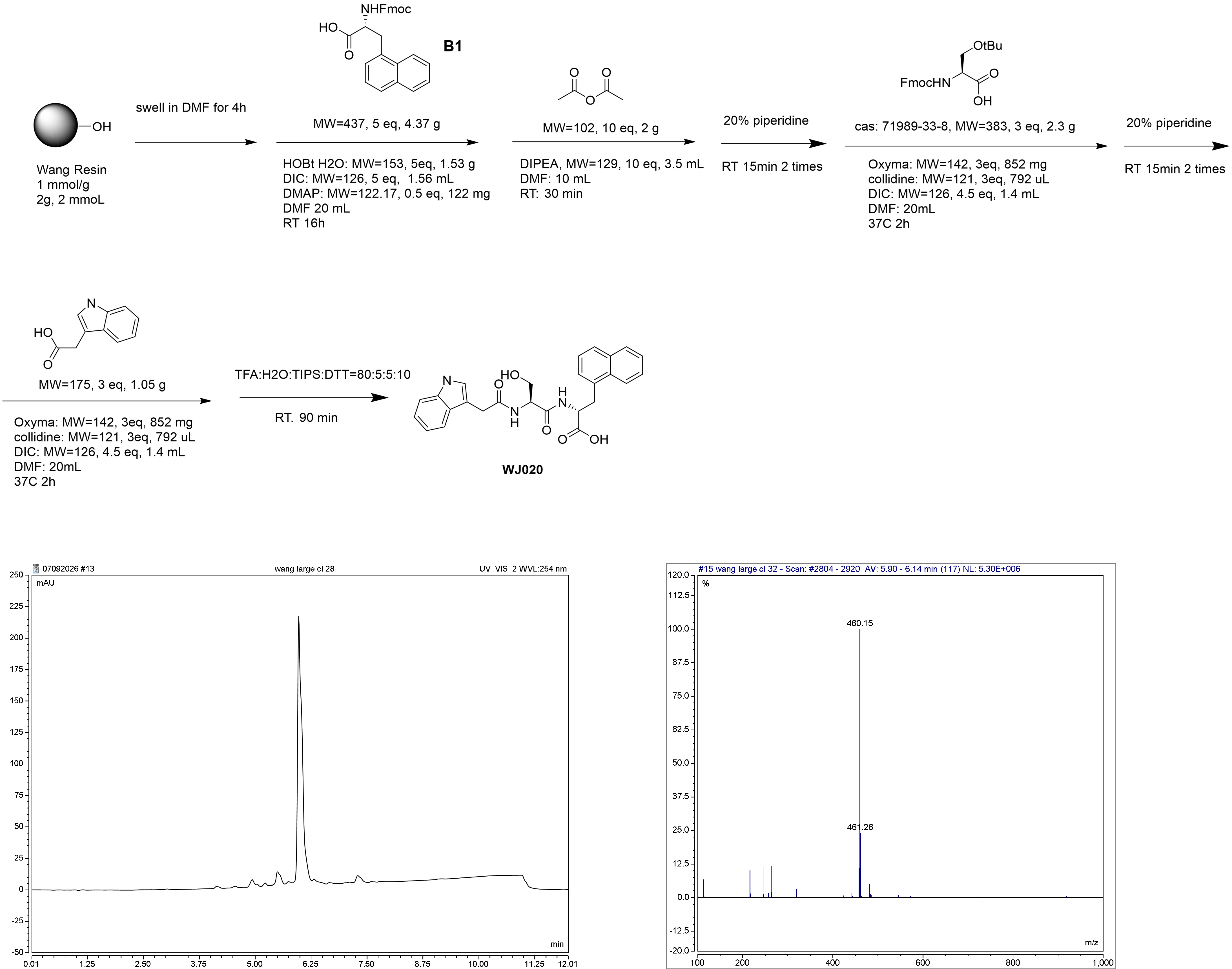

The synthesis of WJ020 follows a standard Wang resin solid phase synthesis. The final product was purified by HPLC and characterized by LC-MS. The calculated mass of WJ020 is 459.18, the mass found is 460.15.

**WJ020**:

^1^H NMR (400 MHz, DMSO) δ 10.85 (d, *J* = 2.5 Hz, 1H), 8.25 (d, *J* = 8.1 Hz, 1H), 8.12 (d, *J* = 8.3 Hz, 1H), 7.92 (dd, *J* = 8.1, 1.5 Hz, 1H), 7.86 (d, *J* = 8.2 Hz, 1H), 7.79 (dd, *J* = 7.8, 1.6 Hz, 1H), 7.62 – 7.47 (m, 4H), 7.38 – 7.29 (m, 3H), 7.21 (d, *J* = 2.3 Hz, 1H), 7.05 (ddd, *J* = 8.2, 6.9, 1.2 Hz, 1H), 6.95 (ddd, *J* = 8.0, 6.9, 1.1 Hz, 1H), 4.54 (td, *J* = 8.4, 5.1 Hz, 1H), 4.34 (dt, *J* = 7.9, 5.8 Hz, 1H), 3.58 – 3.49 (m, 2H), 3.45 – 3.38 (m, 2H), 3.36 – 3.28 (m, 1H), 3.22 (dd, *J* = 14.2, 8.9 Hz, 1H).

^13^C NMR (101 MHz, DMSO) δ 173.35, 171.11, 170.50, 162.77, 136.54, 133.88, 133.75, 131.87, 129.17, 128.07, 127.76, 127.66, 126.69, 126.05, 125.82, 124.28, 123.76, 121.36, 119.23, 118.75, 111.71, 109.23, 62.34, 55.35, 53.35, 36.24, 34.98, 32.89, 31.24.

MS (ESI, positive) m/z calculated for C_26_H_27_N_3_O_5_ [M+H]^+^ : 460.19, found: 460.15.

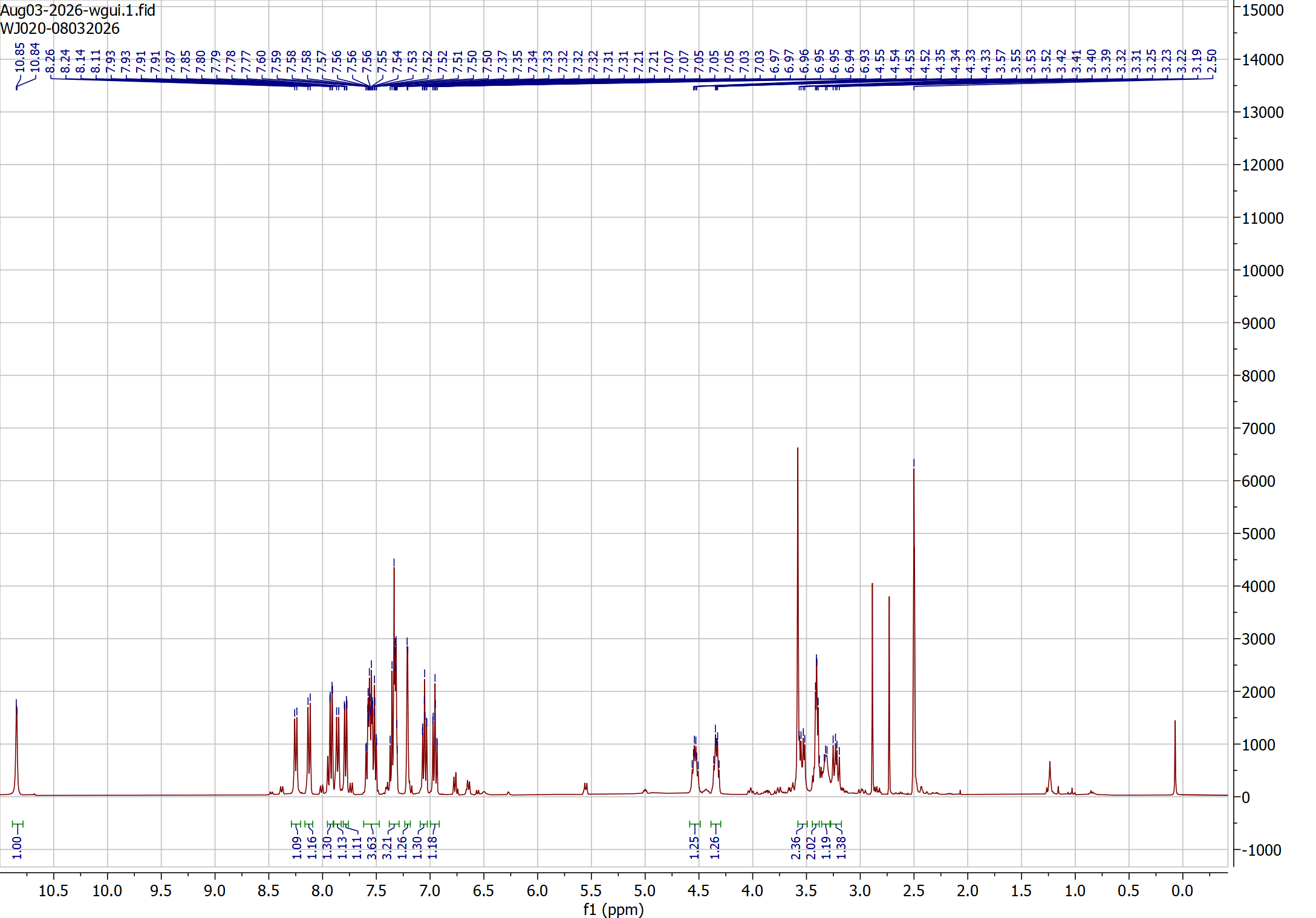

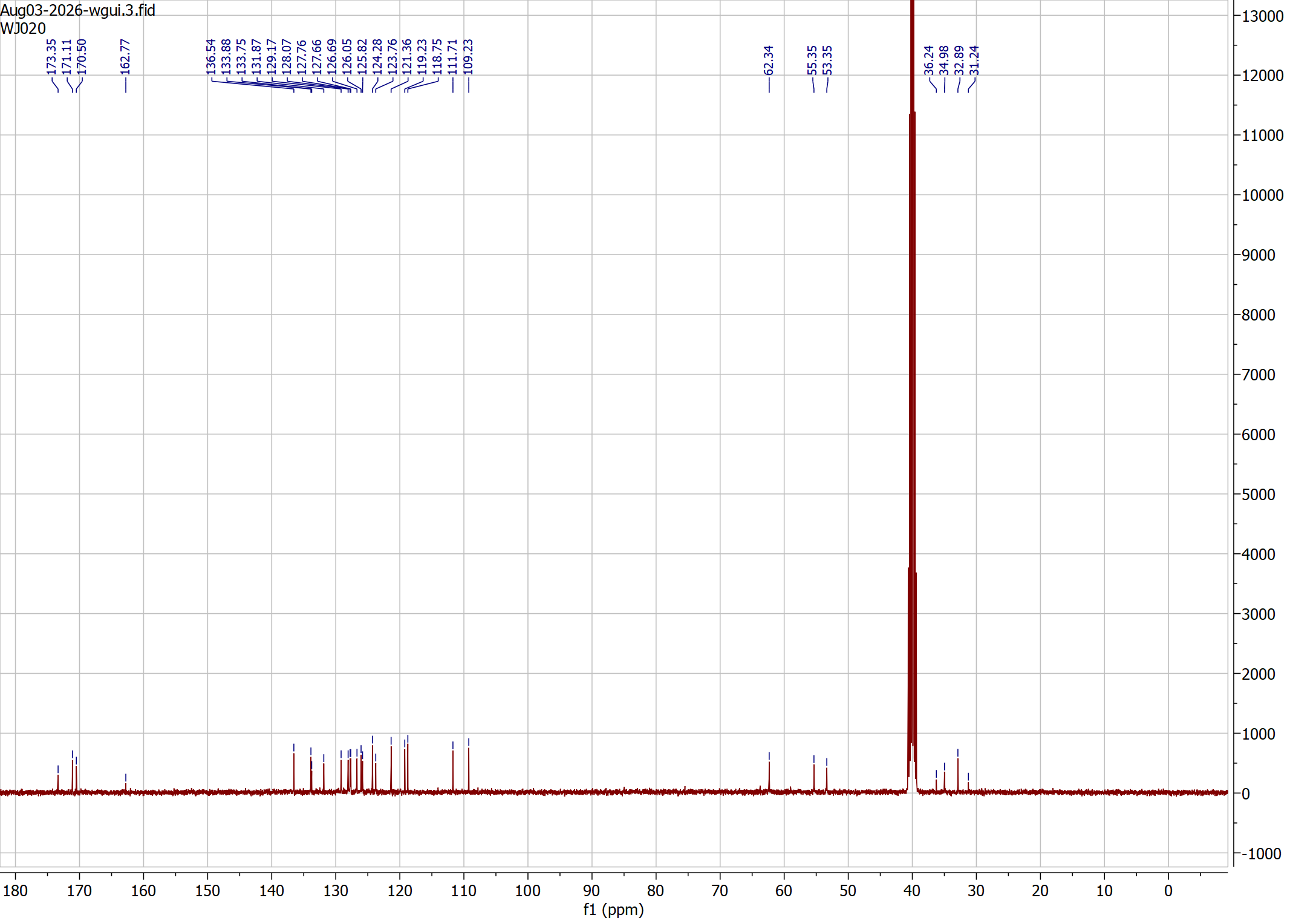

**Scheme S7: Generation of CFTR DUBTACs.**

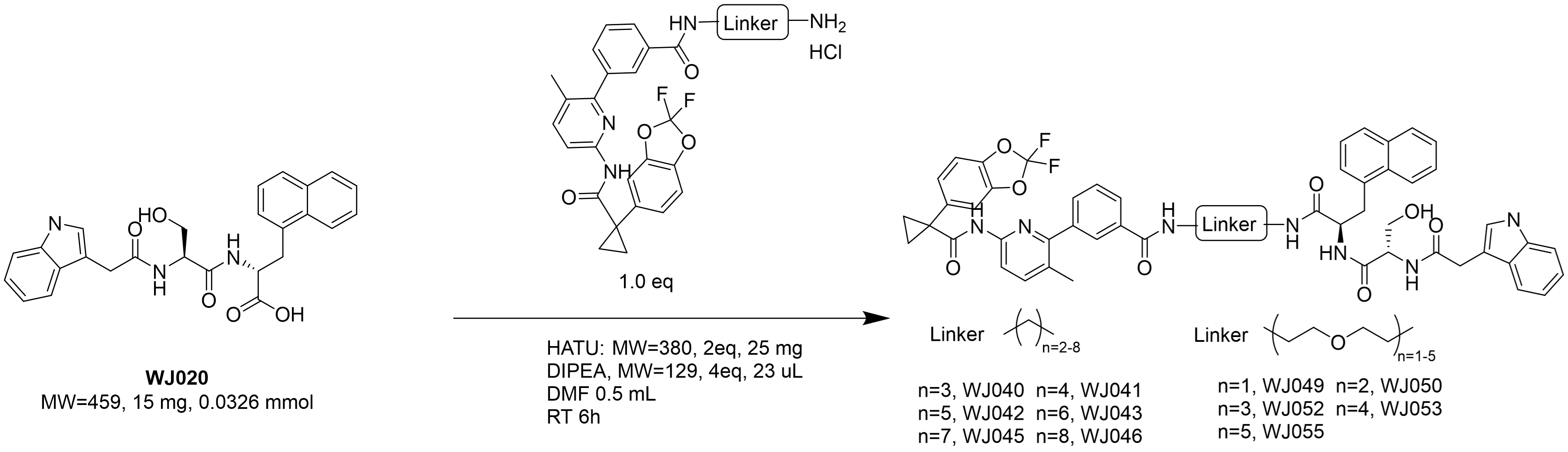

The CFTR ligand with different linkers was generated as previous publication reported.^2^ The coupling of WJ020 to CFTR ligands is by incubation of 15 mg, 0.0326 mmol WJ020 with HATU (2eq, 25 mg), DIPEA (4eq, 23 uL) and 1 equivalent of CFTR-Linker-NH2 in 0.5 mL DMF for 6 hours at room temperature. LC-MS was used to monitor the reaction. After the reaction completed. The reaction mixture was injected into C-18 column and HPLC purification to collect the desire fractions. The final products were characterized using LC-MS and NMR.

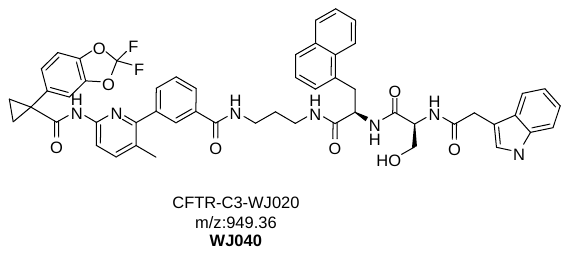

**WJ040**

^1^H NMR (400 MHz, DMSO) δ 10.84 (s, 1H), 8.93 (s, 1H), 8.44 (t, *J* = 5.6 Hz, 1H), 8.22 (dd, *J* = 23.7, 8.2 Hz, 2H), 8.03 (d, *J* = 7.0 Hz, 1H), 7.87 (dt, *J* = 17.0, 9.2 Hz, 5H), 7.73 (t, *J* = 7.3 Hz, 2H), 7.61 – 7.46 (m, 6H), 7.39 – 7.27 (m, 5H), 7.19 (d, *J* = 2.4 Hz, 1H), 7.04 (t, *J* = 7.5 Hz, 1H), 6.94 (t, *J* = 7.3 Hz, 1H), 4.88 (t, *J* = 5.4 Hz, 1H), 4.51 – 4.40 (m, 1H), 4.21 (q, *J* = 6.1 Hz, 1H), 3.56 (dd, *J* = 17.6, 3.9 Hz, 3H), 3.42 (t, *J* = 6.8 Hz, 2H), 3.25 – 3.08 (m, 4H), 2.99 (d, *J* = 6.5 Hz, 2H), 2.21 (s, 3H), 1.50 (q, *J* = 4.2 Hz, 2H), 1.40 – 1.34 (m, 2H), 1.15 (d, *J* = 2.8 Hz, 2H).

^13^C NMR (101 MHz, DMSO) δ 171.69, 155.92, 149.51, 140.04, 136.74, 136.54, 134.85, 128.46, 127.00, 124.20, 118.74, 113.59, 112.67, 110.58, 109.17, 31.81, 26.87, 19.18, 16.14.

MS (ESI, positive) m/z calculated for C_53_H_51_F_2_N_7_O_8_ [M+H]^+^ : 950.37, found: 950.41.

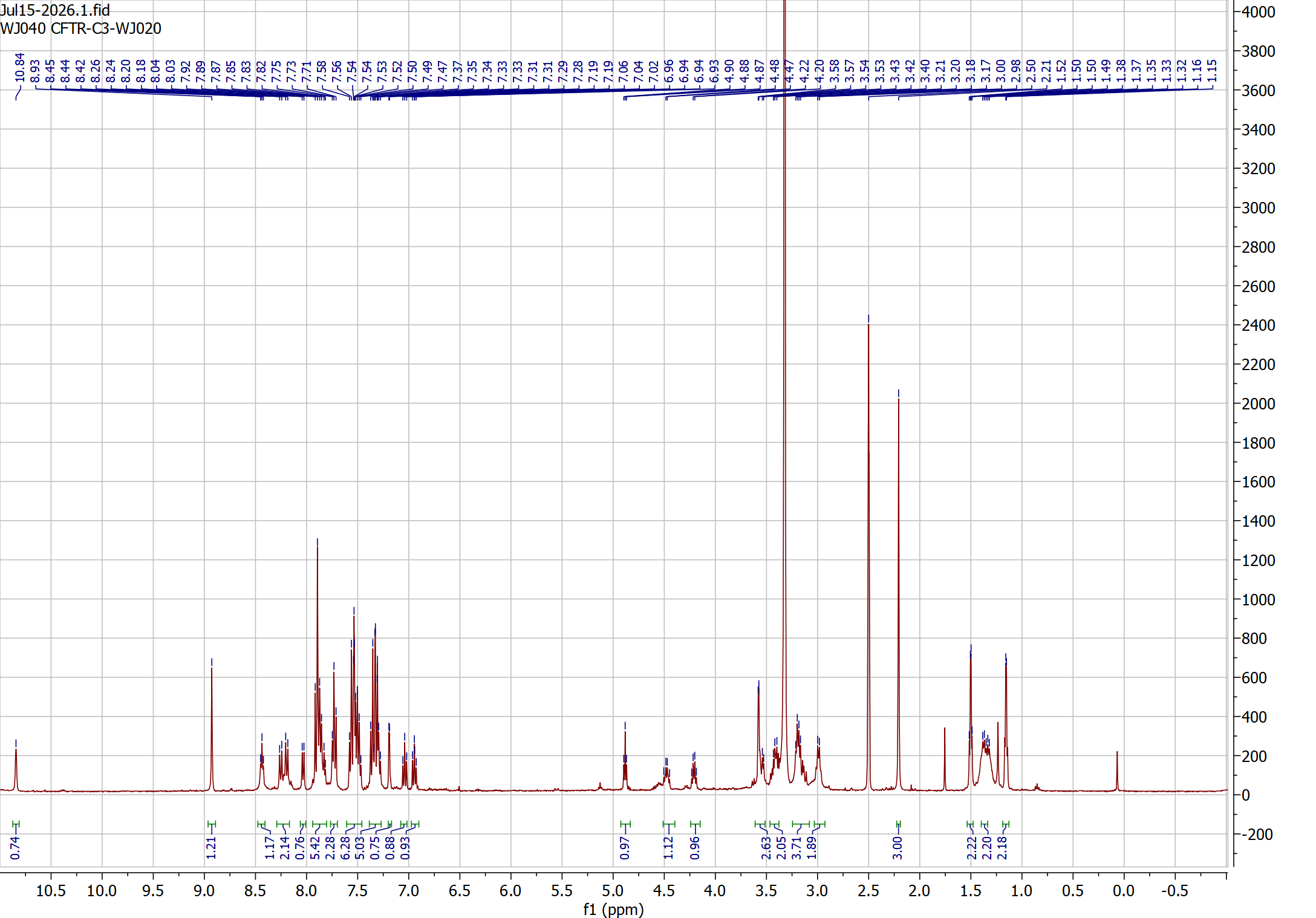

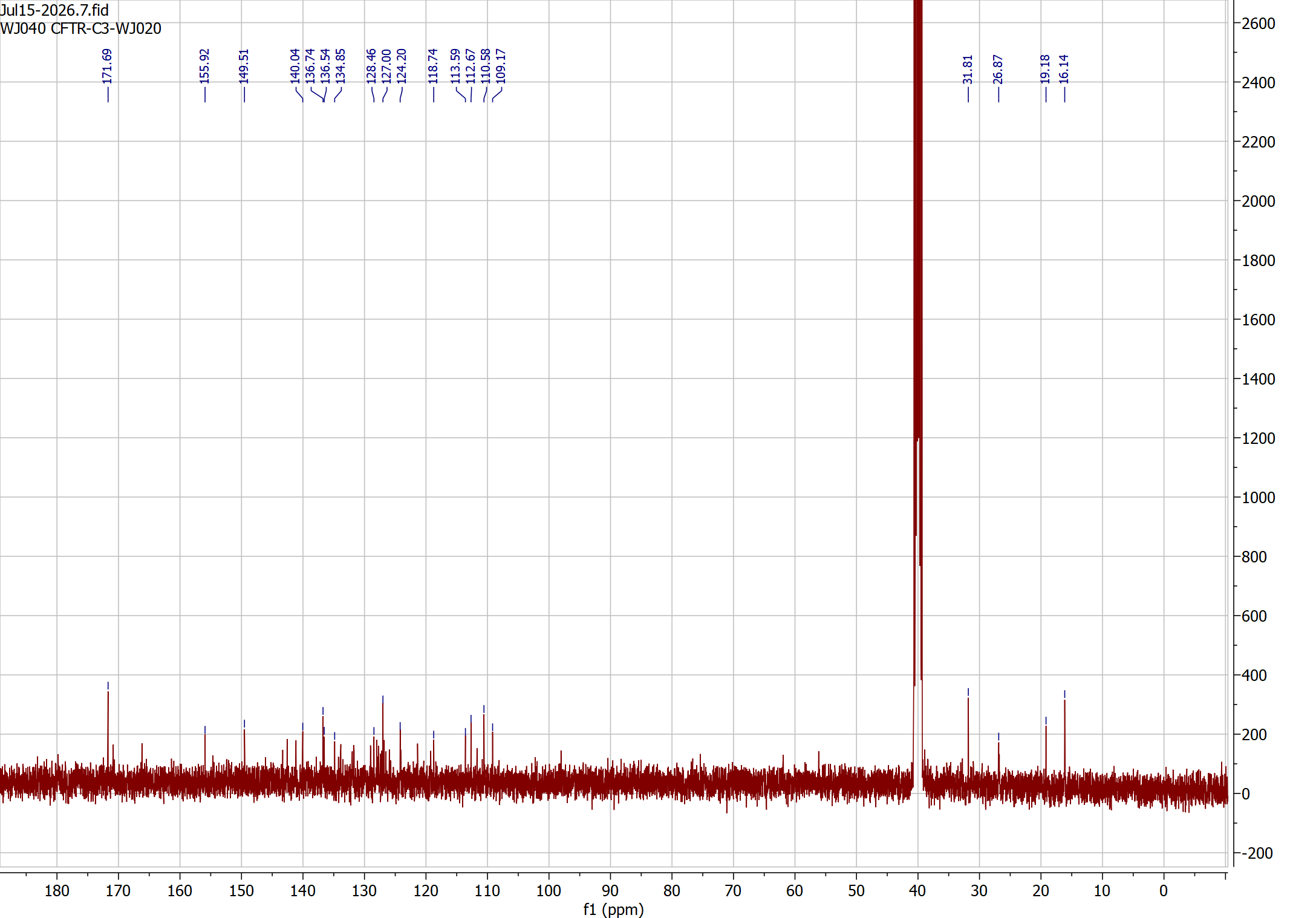

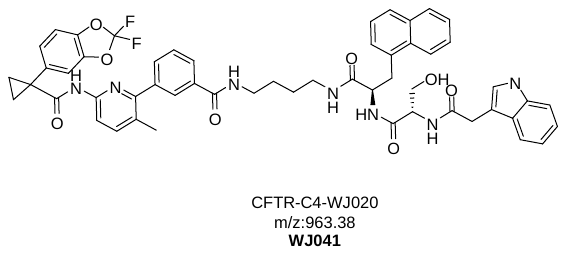

**WJ041**

^1^H NMR (400 MHz, DMSO) δ 10.84 (s, 1H), 8.93 (s, 1H), 8.44 (t, *J* = 5.5 Hz, 1H), 8.23 (dd, *J* = 25.0, 8.2 Hz, 2H), 8.05 (d, *J* = 6.9 Hz, 1H), 7.94 – 7.83 (m, 4H), 7.83 – 7.70 (m, 3H), 7.60 – 7.46 (m, 6H), 7.42 – 7.27 (m, 5H), 7.20 (d, *J* = 2.3 Hz, 1H), 7.07 – 7.02 (m, 1H), 6.94 (ddd, *J* = 8.0, 7.0, 1.1 Hz, 1H), 4.89 (t, *J* = 5.3 Hz, 1H), 4.47 (d, *J* = 5.6 Hz, 1H), 4.21 (d, *J* = 6.4 Hz, 1H), 3.62 – 3.53 (m, 2H), 3.49 – 3.38 (m, 2H), 3.20 (q, *J* = 7.2 Hz, 3H), 3.02 – 2.91 (m, 2H), 2.21 (s, 3H), 1.51 (q, *J* = 4.2 Hz, 2H), 1.48 – 1.39 (m, 2H), 1.28 (d, *J* = 8.5 Hz, 2H), 1.16 (d, *J* = 2.5 Hz, 2H).

^13^C NMR (101 MHz, DMSO) δ 171.69, 170.79, 170.69, 166.16, 155.92, 149.51, 143.31, 142.59, 141.14, 140.02, 136.75, 136.54, 134.88, 134.27, 133.86, 132.00, 131.74, 128.44, 128.00, 127.90, 127.73, 127.45, 127.22, 127.01, 126.79, 126.51, 125.97, 125.75, 124.18, 124.10, 121.37, 119.23, 118.73, 113.56, 112.69, 111.70, 110.59, 109.18, 61.90, 56.19, 54.28, 32.78, 31.82, 29.23, 28.95, 24.27, 19.18, 16.15.

MS (ESI, positive) m/z calculated for C_54_H_52_F_2_N_7_O_8_ [M+H]^+^ : 964.38, found: 964.45.

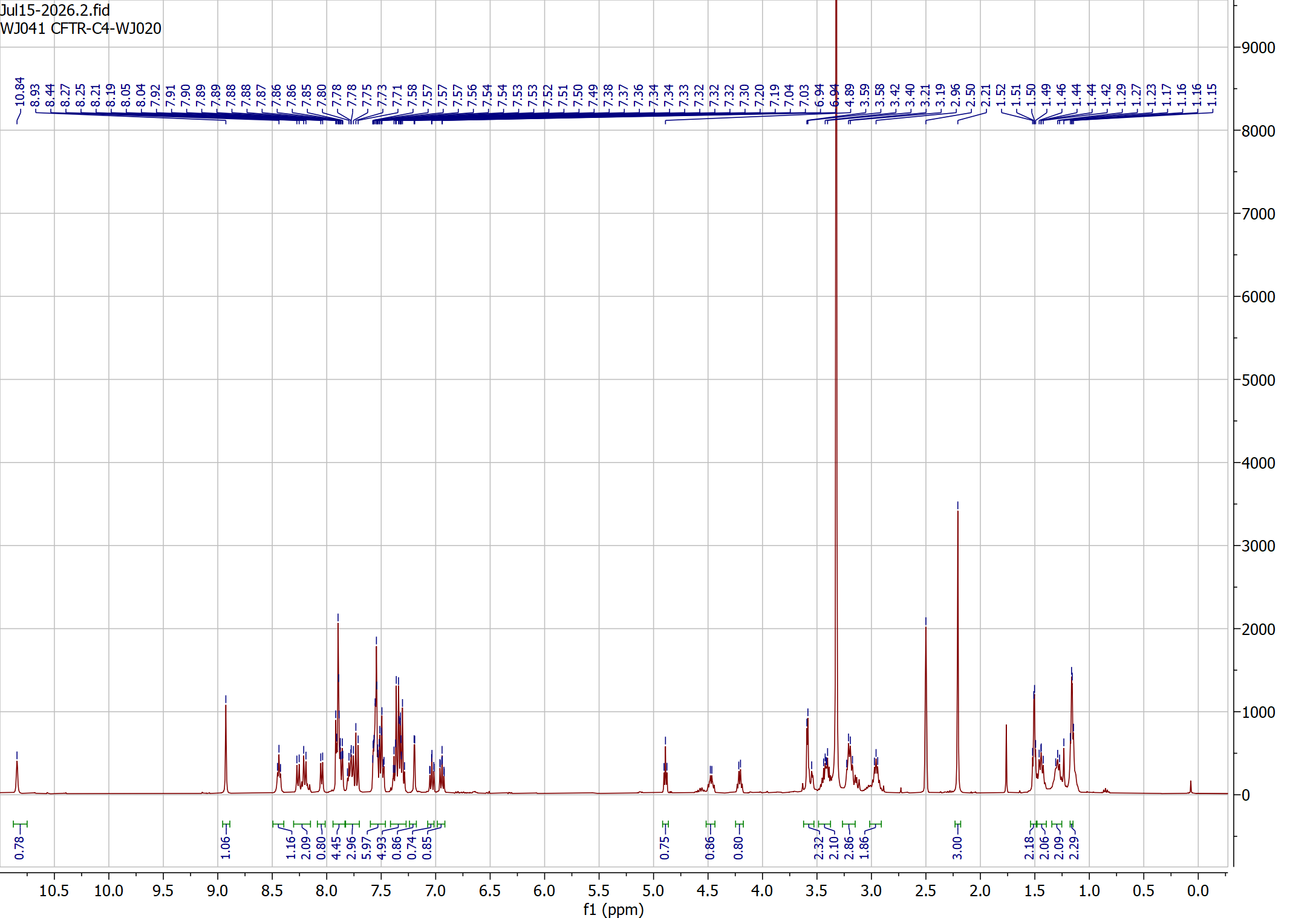

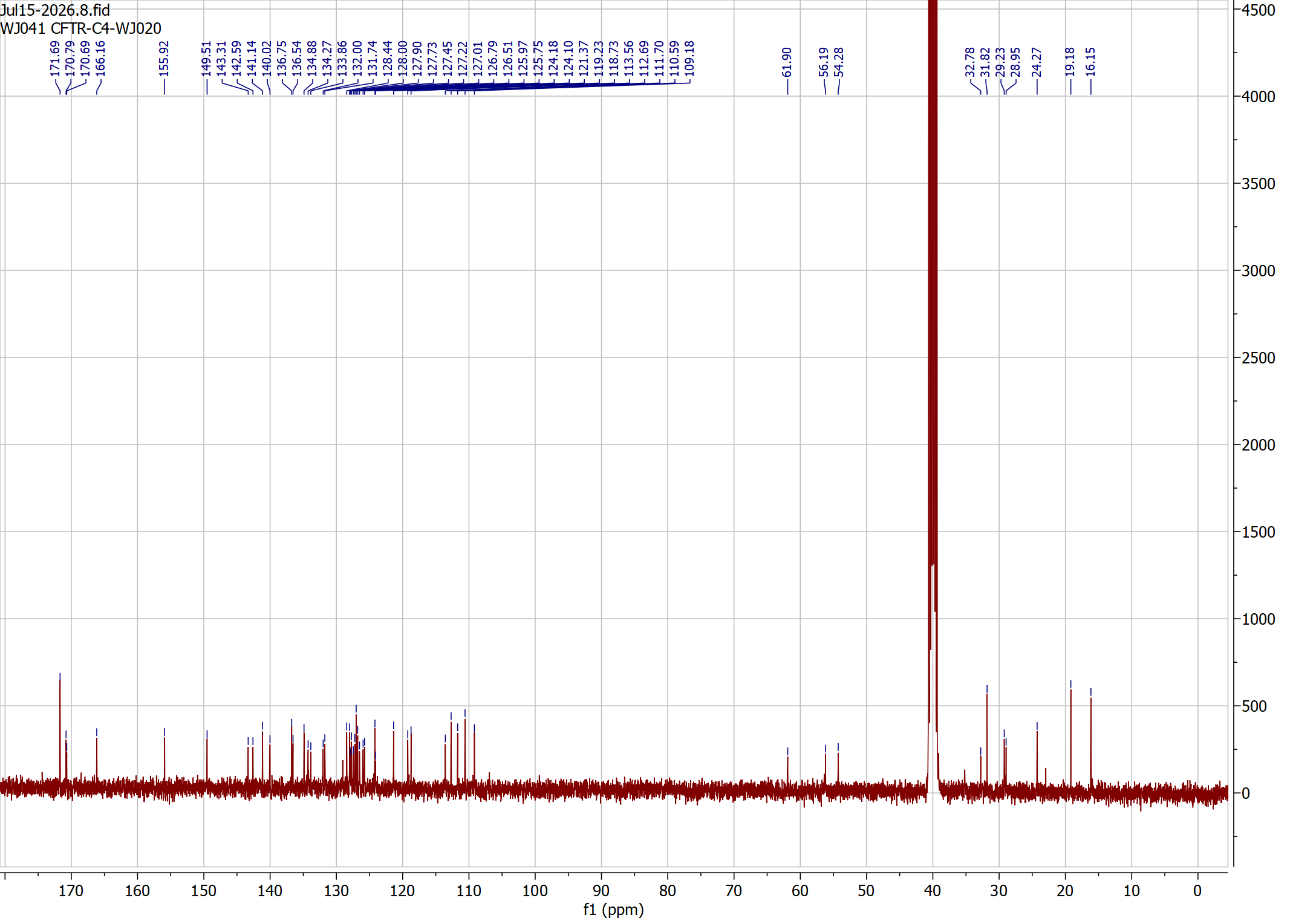

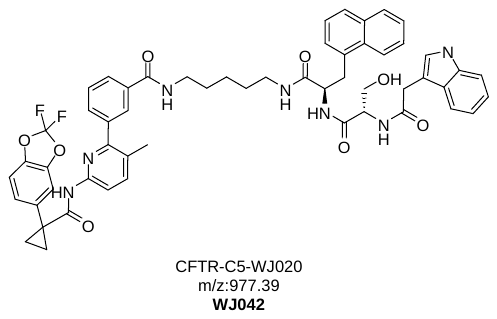

**WJ042**

^1^H NMR (400 MHz, DMSO) δ 10.85 (d, *J* = 2.5 Hz, 1H), 8.93 (s, 1H), 8.46 (t, *J* = 5.6 Hz, 1H), 8.26 (d, *J* = 8.3 Hz, 1H), 8.20 (d, *J* = 8.3 Hz, 1H), 8.05 (d, *J* = 6.9 Hz, 1H), 7.89 (td, *J* = 8.1, 6.1 Hz, 4H), 7.81 – 7.71 (m, 3H), 7.60 – 7.44 (m, 6H), 7.41 – 7.28 (m, 5H), 7.20 (d, *J* = 2.3 Hz, 1H), 7.04 (ddd, *J* = 8.1, 7.0, 1.2 Hz, 1H), 6.94 (ddd, *J* = 8.0, 7.0, 1.1 Hz, 1H), 4.89 (t, *J* = 5.4 Hz, 1H), 4.47 (td, *J* = 8.5, 5.3 Hz, 1H), 4.22 (q, *J* = 6.1 Hz, 1H), 3.62 – 3.53 (m, 2H), 3.48 – 3.36 (m, 2H), 3.23 (q, *J* = 6.7 Hz, 2H), 3.15 (dd, *J* = 14.0, 8.8 Hz, 1H), 2.95 (dq, *J* = 12.8, 6.8 Hz, 2H), 2.21 (s, 3H), 1.54 – 1.41 (m, 4H), 1.27 – 1.13 (m, 6H).

^13^C NMR (101 MHz, DMSO) δ 171.68, 170.75, 170.68, 166.14, 155.92, 149.52, 143.31, 142.58, 141.14, 140.03, 136.74, 136.55, 134.91, 134.27, 133.86, 132.00, 131.75, 129.03, 128.46, 128.00, 127.89, 127.73, 127.43, 127.21, 127.01, 126.78, 126.50, 125.96, 125.74, 124.18, 124.10, 121.37, 119.23, 118.72, 113.56, 112.69, 111.70, 110.58, 109.18, 61.91, 56.19, 54.28, 35.18, 32.78, 31.82, 29.52, 29.18, 26.66, 26.48, 19.19, 16.15.

MS (ESI, positive) m/z calculated for C_55_H_55_F_2_N_7_O_8_ [M+H]^+^ : 978.40, found: 978.41.

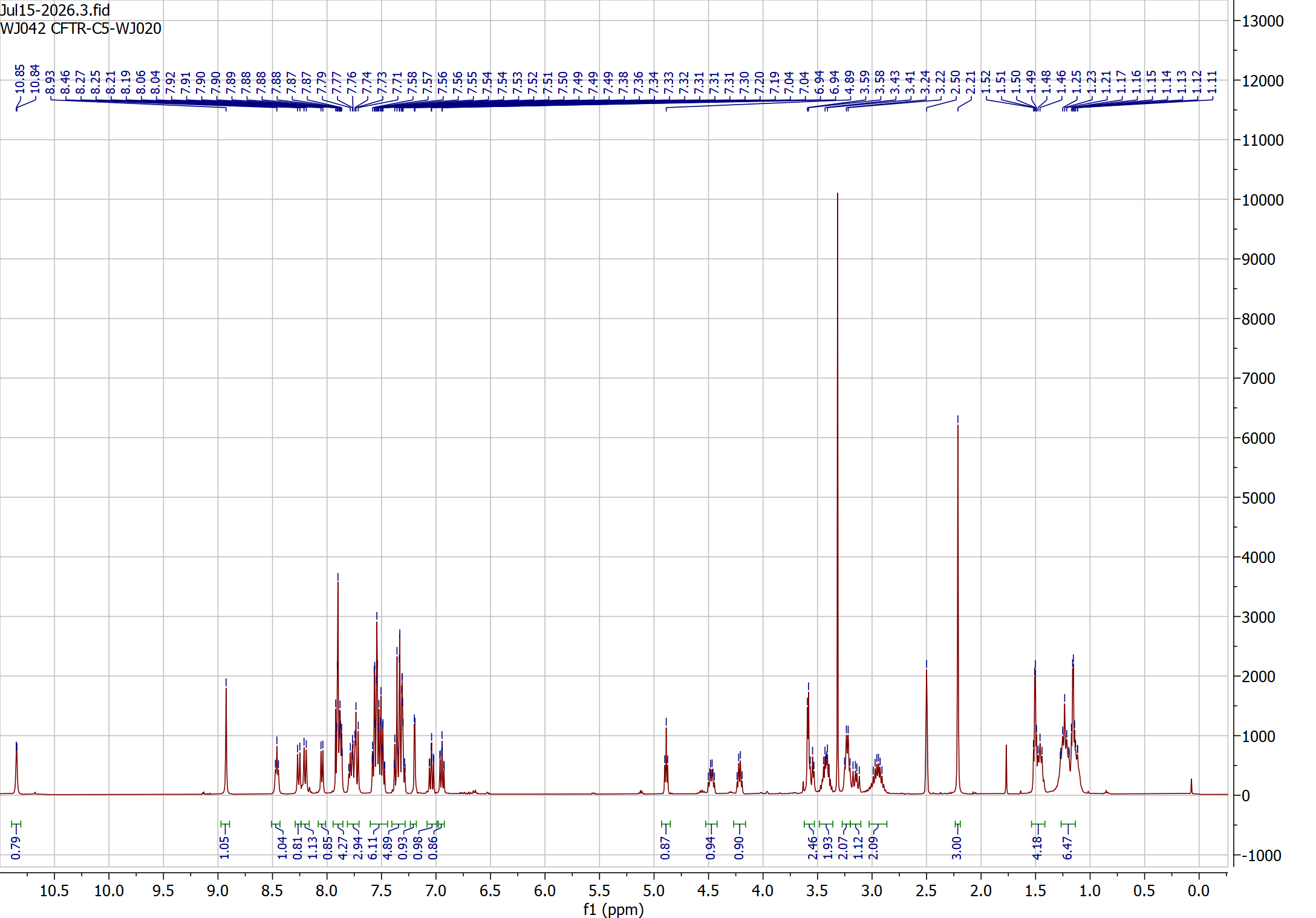

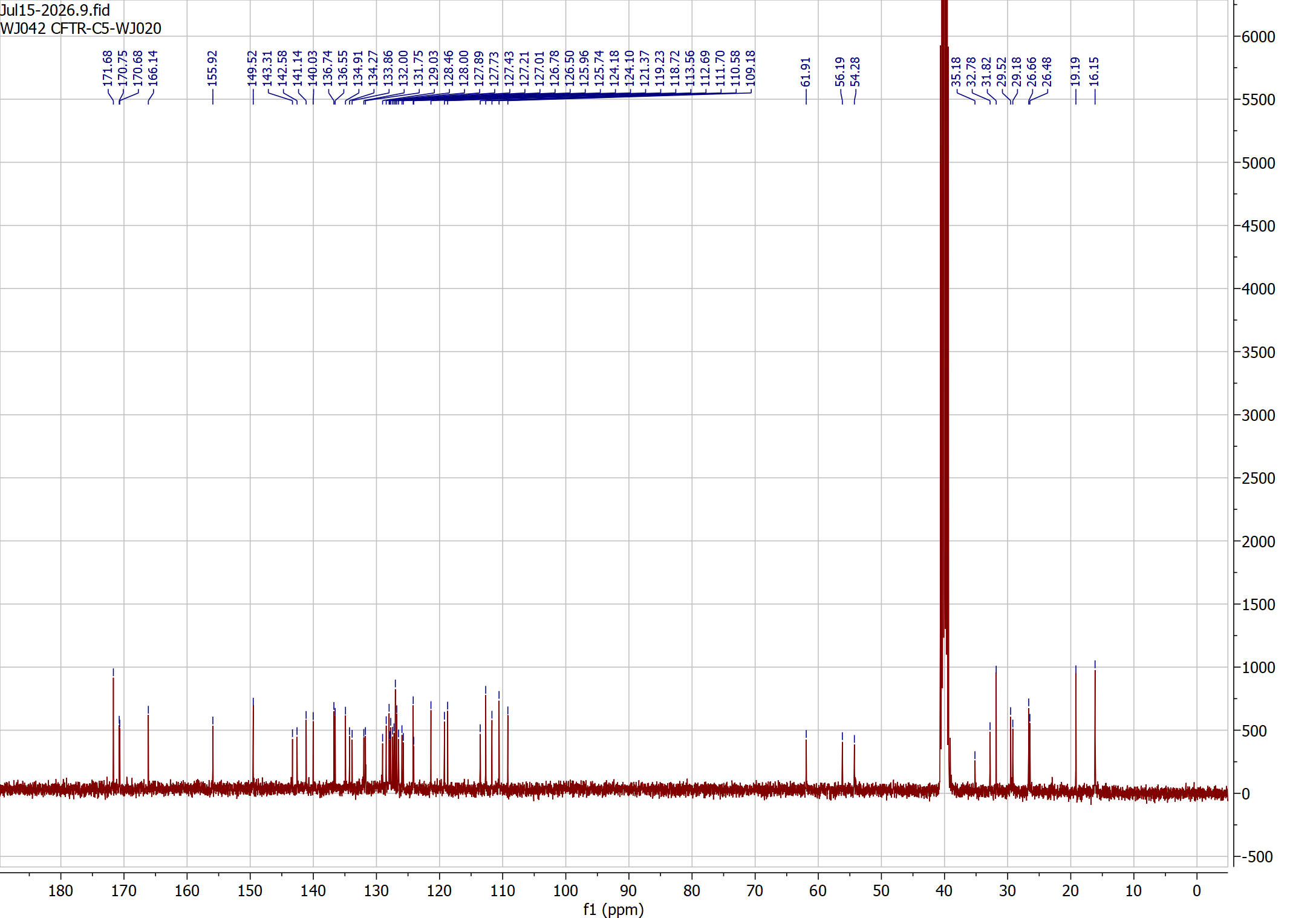

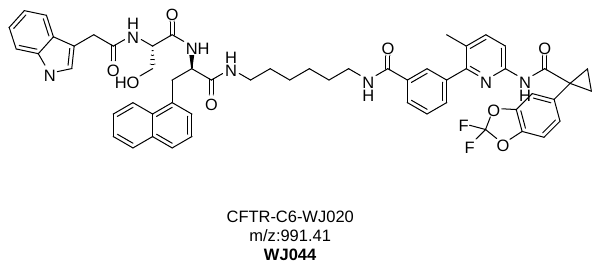

**WJ044**

^1^H NMR (400 MHz, DMSO) δ 10.85 (s, 1H), 8.92 (s, 1H), 8.46 (t, *J* = 5.4 Hz, 1H), 8.23 (dd, *J* = 22.3, 8.1 Hz, 2H), 8.04 (d, *J* = 6.9 Hz, 1H), 7.89 (q, *J* = 7.6 Hz, 4H), 7.82 – 7.70 (m, 3H), 7.61 – 7.44 (m, 6H), 7.34 (ddt, *J* = 12.6, 9.6, 6.0 Hz, 5H), 7.19 (d, *J* = 2.3 Hz, 1H), 7.08 – 7.02 (m, 1H), 6.94 (t, *J* = 7.2 Hz, 1H), 4.88 (t, *J* = 5.4 Hz, 1H), 4.47 (d, *J* = 5.7 Hz, 1H), 4.27 – 4.17 (m, 1H), 3.63 – 3.51 (m, 2H), 3.48 – 3.36 (m, 2H), 3.23 (d, *J* = 6.6 Hz, 2H), 3.14 (dd, *J* = 14.1, 8.8 Hz, 1H), 3.03 – 2.91 (m, 2H), 2.21 (s, 3H), 1.51 (q, *J* = 4.3 Hz, 4H), 1.24 (s, 6H), 1.16 (t, *J* = 3.5 Hz, 2H).

^13^C NMR (101 MHz, DMSO) δ 171.68, 170.75, 170.68, 166.14, 155.92, 149.52, 143.30, 142.58, 141.14, 140.03, 136.74, 136.55, 134.90, 134.27, 133.86, 132.00, 131.75, 129.03, 128.46, 128.00, 127.89, 127.73, 127.43, 127.21, 127.01, 126.78, 126.50, 125.96, 125.74, 124.18, 124.10, 121.37, 119.23, 118.73, 113.56, 112.69, 111.70, 110.58, 109.18, 61.91, 56.19, 54.28, 35.18, 32.78, 31.82, 29.52, 29.18, 26.66, 26.48, 22.96, 19.18, 16.15.

MS (ESI, positive) m/z calculated for C_56_H_57_F_2_N_7_O_8_ [M+H]^+^ : 992.42, found: 992.38.

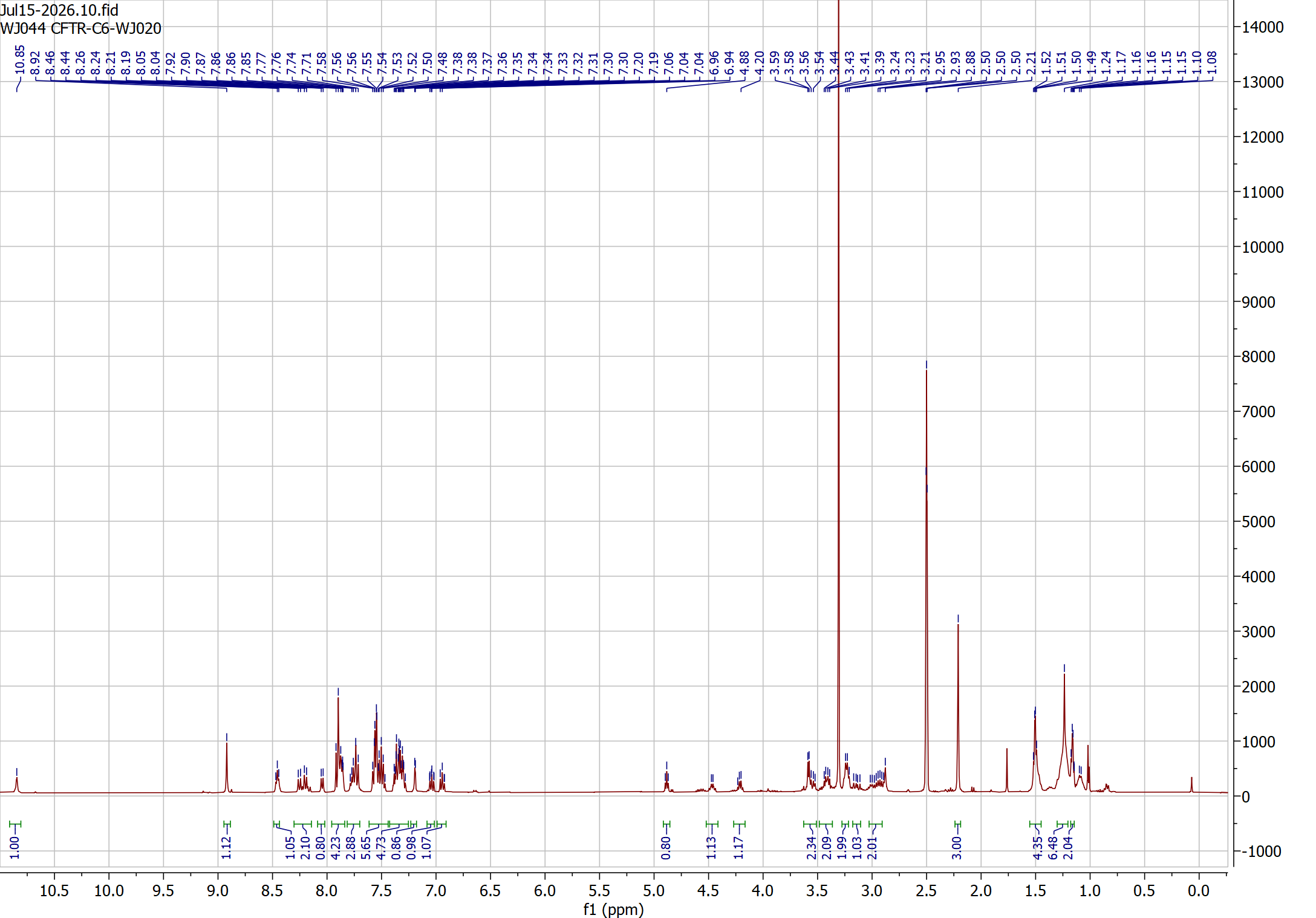

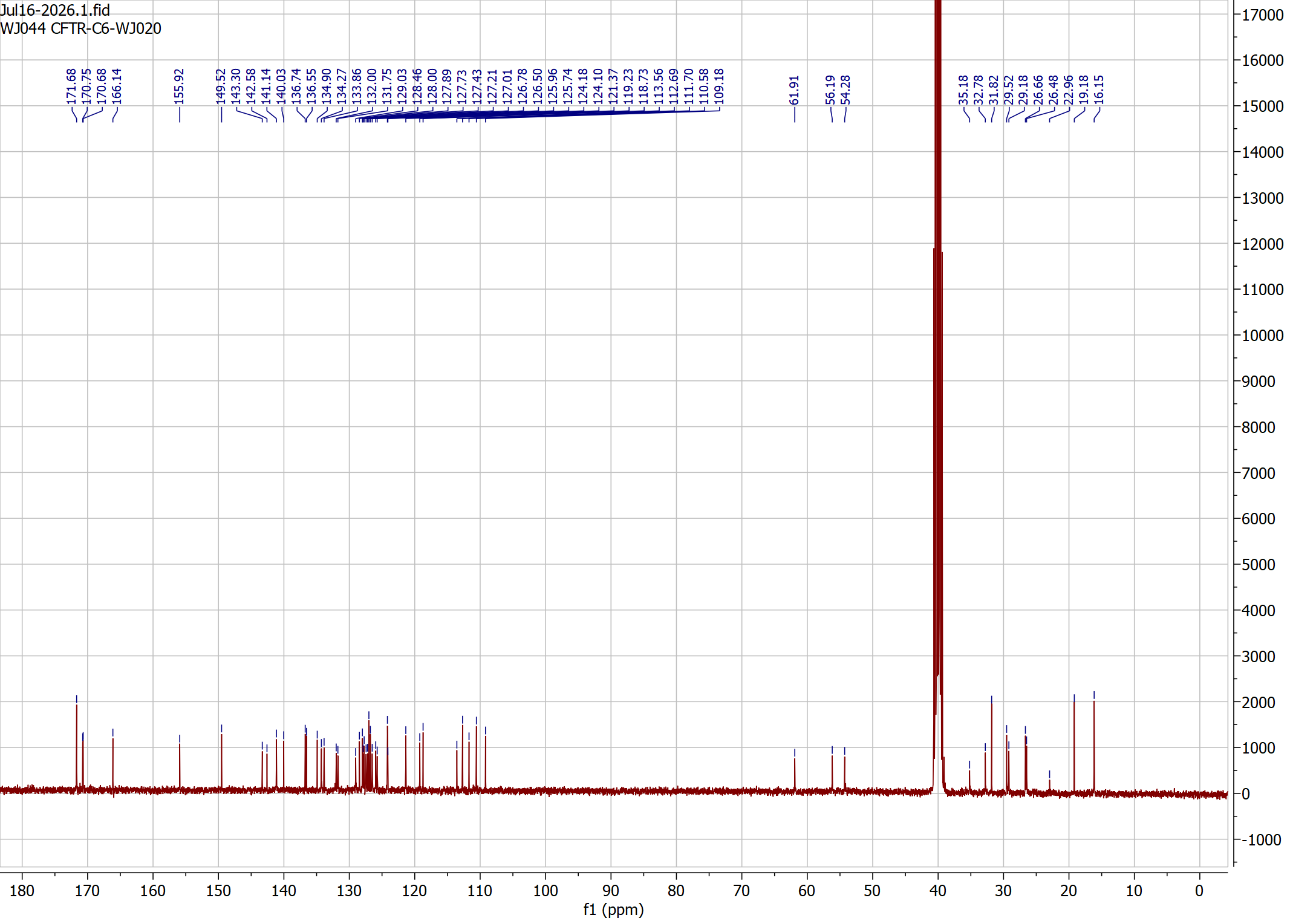

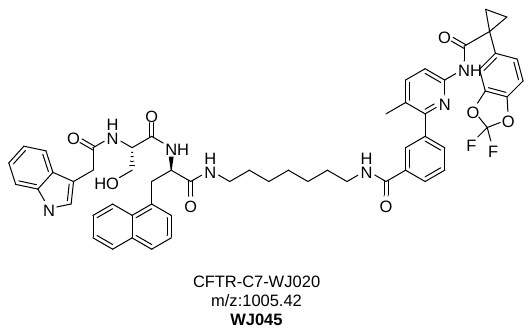

**WJ045**

^1^H NMR (400 MHz, DMSO) δ 10.84 (s, 1H), 8.92 (s, 1H), 8.45 (t, *J* = 5.6 Hz, 1H), 8.23 (dd, *J* = 22.7, 8.2 Hz, 2H), 8.04 (d, *J* = 6.9 Hz, 1H), 7.94 – 7.84 (m, 4H), 7.74 (dd, *J* = 12.8, 8.5 Hz, 3H), 7.61 – 7.46 (m, 6H), 7.40 – 7.27 (m, 5H), 7.19 (d, *J* = 2.4 Hz, 1H), 7.08 – 7.00 (m, 1H), 6.94 (ddd, *J* = 7.9, 7.0, 1.0 Hz, 1H), 4.88 (t, *J* = 5.4 Hz, 1H), 4.47 (d, *J* = 5.5 Hz, 1H), 4.21 (d, *J* = 6.3 Hz, 1H), 3.57 (dd, *J* = 12.5, 6.7 Hz, 2H), 3.41 (dt, *J* = 8.0, 5.5 Hz, 2H), 3.27 – 3.21 (m, 2H), 3.14 (dd, *J* = 14.1, 8.9 Hz, 1H), 2.93 (ddd, *J* = 18.7, 12.9, 6.2 Hz, 3H), 2.21 (s, 3H), 1.50 (p, *J* = 5.0 Hz, 4H), 1.24 (s, 8H), 1.16 (t, *J* = 3.5 Hz, 2H).

^13^C NMR (101 MHz, DMSO) δ 171.69, 170.72, 166.11, 155.92, 153.21, 149.51, 143.31, 141.15, 140.02, 136.75, 134.90, 131.74, 127.99, 127.02, 124.17, 121.36, 112.70, 111.70, 110.59, 109.18, 56.18, 53.53, 31.82, 29.54, 28.95, 26.93, 19.18, 16.15.

MS (ESI, positive) m/z calculated for C_57_H_59_F_2_N_7_O_8_ [M+H]^+^ : 1006.43, found: 1006.41.

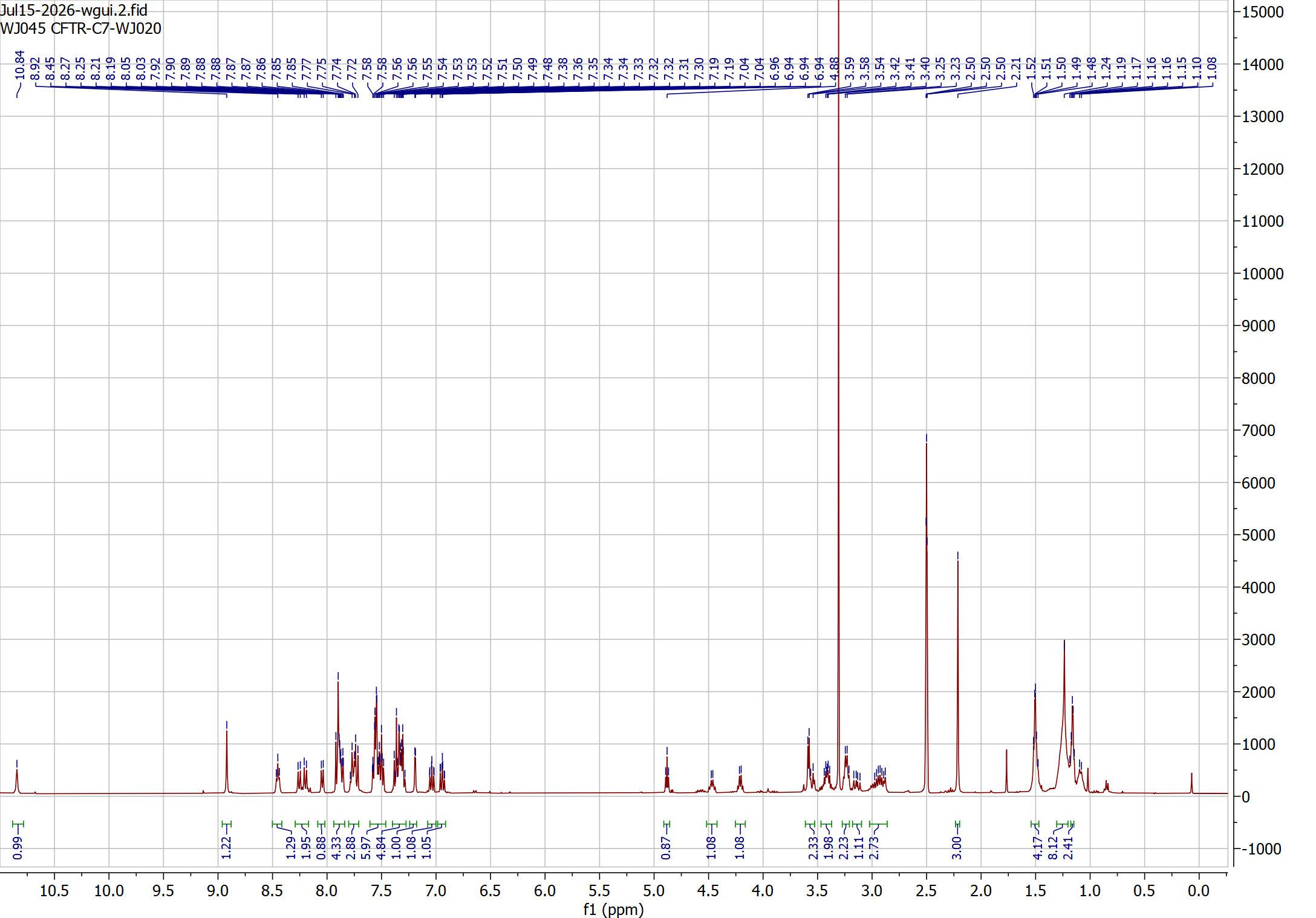

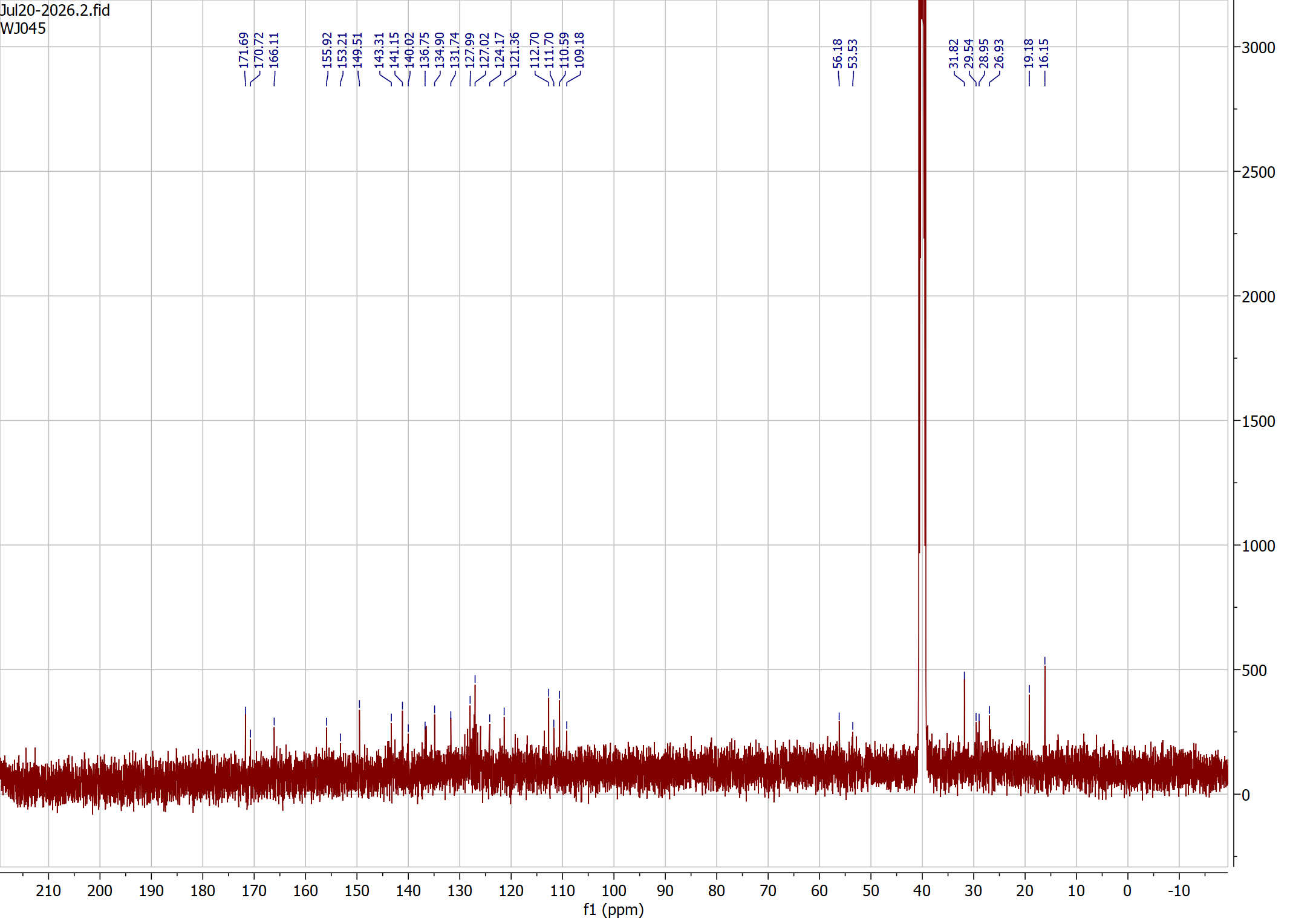

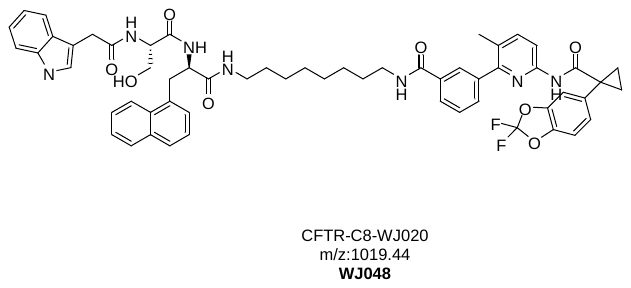

**WJ048**

^1^H NMR (400 MHz, DMSO) δ 10.85 (d, *J* = 2.5 Hz, 1H), 8.92 (s, 1H), 8.45 (t, *J* = 5.7 Hz, 1H), 8.26 (d, *J* = 8.3 Hz, 1H), 8.20 (d, *J* = 8.3 Hz, 1H), 8.05 (d, *J* = 7.0 Hz, 1H), 7.94 – 7.83 (m, 4H), 7.81 – 7.71 (m, 3H), 7.60 – 7.48 (m, 6H), 7.39 – 7.30 (m, 5H), 7.19 (d, *J* = 2.3 Hz, 1H), 7.04 (ddd, *J* = 8.1, 7.0, 1.2 Hz, 1H), 6.94 (ddd, *J* = 8.0, 7.0, 1.1 Hz, 1H), 4.88 (t, *J* = 5.4 Hz, 1H), 4.52 – 4.43 (m, 1H), 4.21 (d, *J* = 6.2 Hz, 1H), 3.59 (d, *J* = 4.6 Hz, 2H), 3.42 (dt, *J* = 8.2, 5.5 Hz, 2H), 3.24 (q, *J* = 6.7 Hz, 2H), 3.14 (dd, *J* = 13.9, 8.8 Hz, 1H), 2.94 (dd, *J* = 13.0, 6.4 Hz, 2H), 2.21 (s, 3H), 1.50 (p, *J* = 4.4 Hz, 4H), 1.29 – 1.14 (m, 10H), 1.09 (d, *J* = 7.4 Hz, 2H).

^13^C NMR (101 MHz, DMSO) δ 171.68, 170.73, 170.68, 166.11, 155.91, 149.51, 143.31, 142.59, 141.15, 140.01, 136.74, 136.54, 134.90, 134.28, 133.86, 132.00, 131.73, 129.03, 128.45, 127.99, 127.89, 127.73, 127.43, 127.22, 127.02, 126.78, 126.51, 125.97, 125.74, 124.16, 121.36, 119.21, 118.72, 113.54, 112.70, 111.70, 110.58, 109.18, 61.89, 56.19, 54.27, 35.18, 31.81, 29.55, 29.19, 26.97, 26.74, 19.19, 16.15.

MS (ESI, positive) m/z calculated for C_58_H_61_F_2_N_7_O_8_ [M+H]^+^ : 1020.45, found: 1020.51.

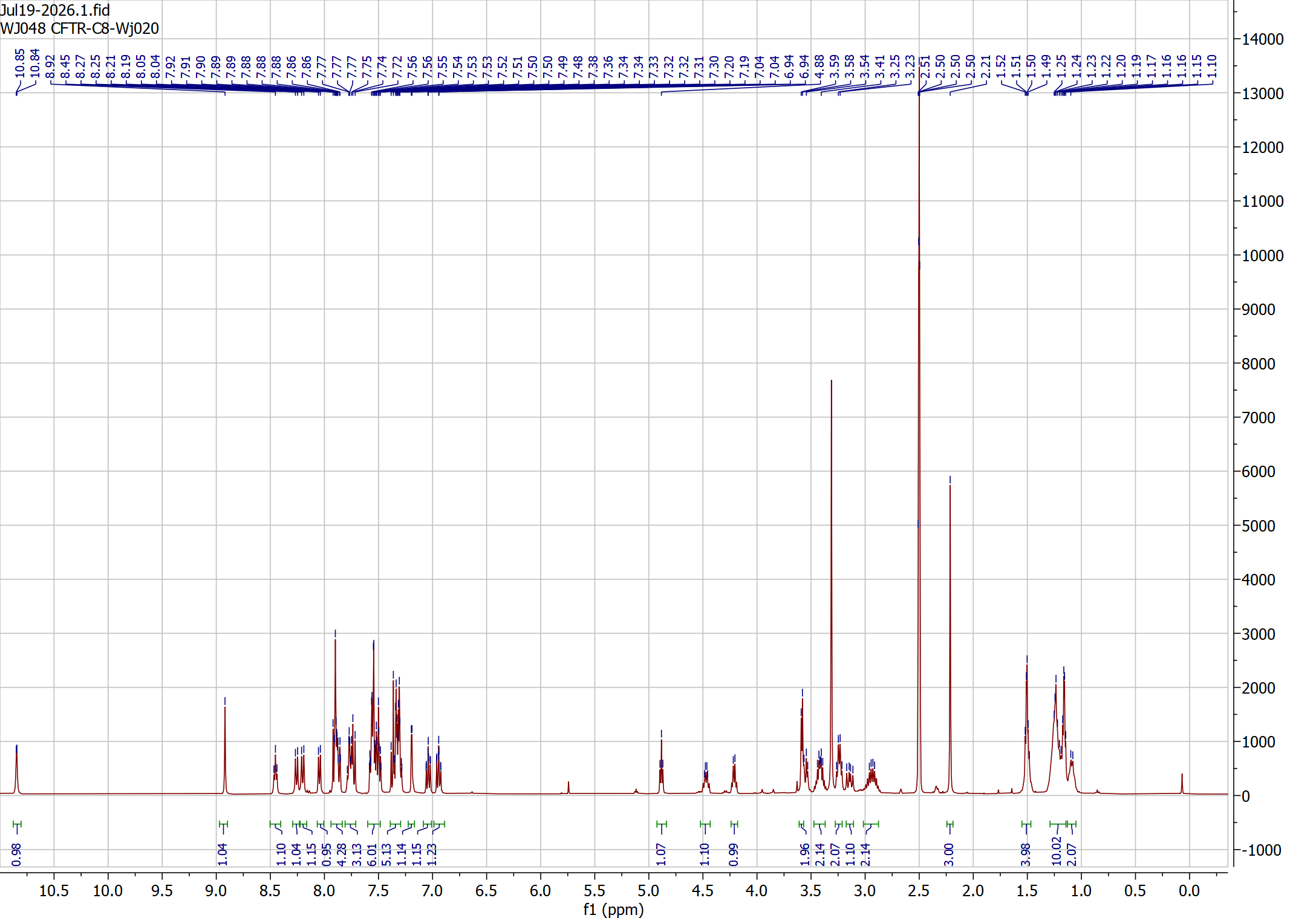

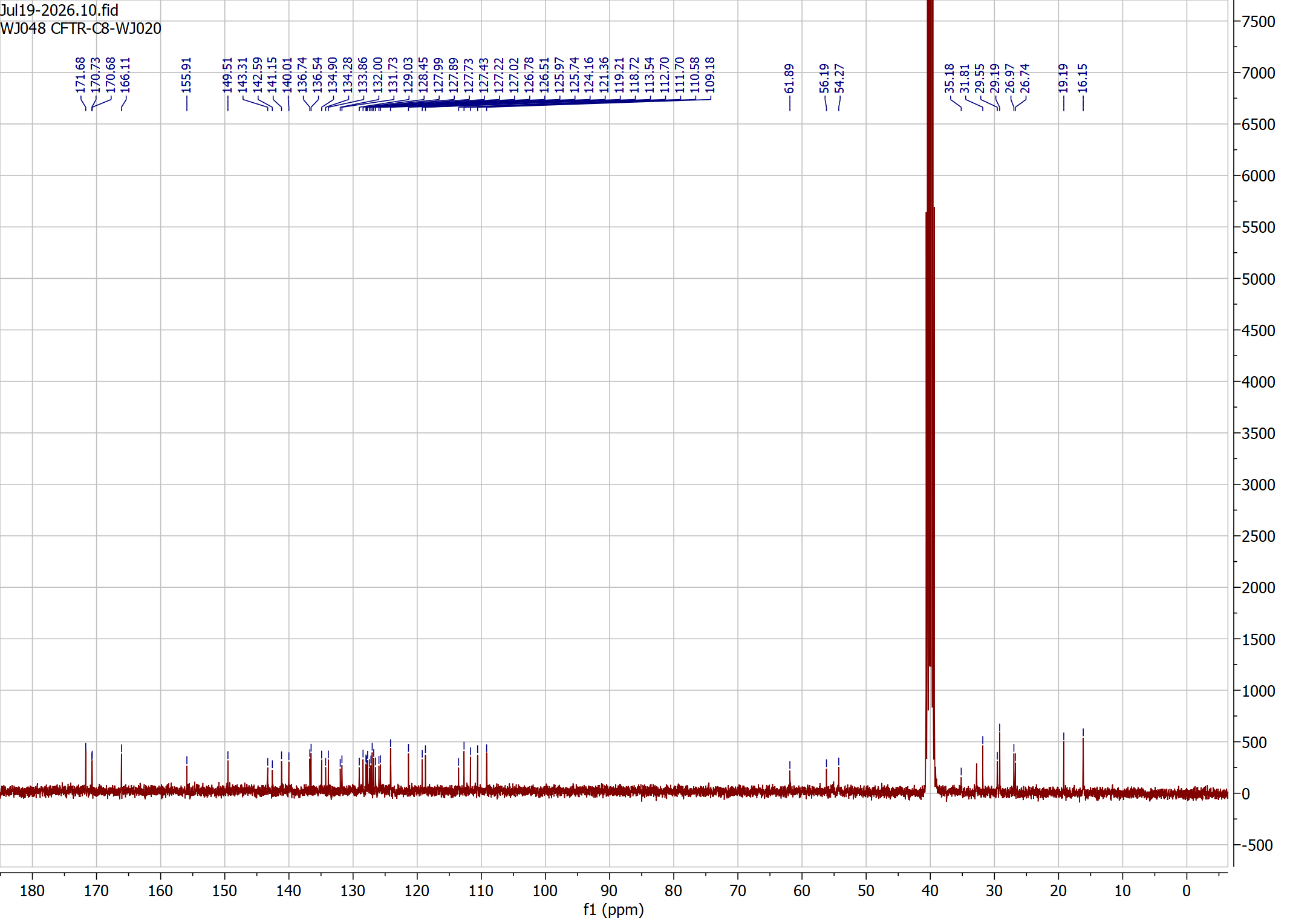

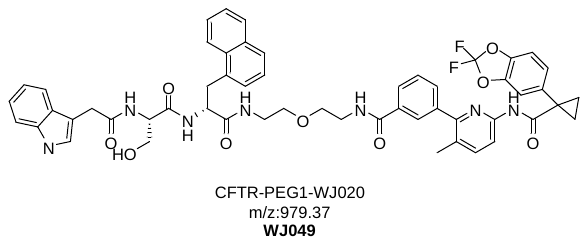

**WJ049**

^1^H NMR (400 MHz, DMSO) δ 10.84 (d, *J* = 2.3 Hz, 1H), 8.94 (s, 1H), 8.49 (t, *J* = 5.3 Hz, 1H), 8.28 (d, *J* = 8.3 Hz, 1H), 8.19 (d, *J* = 8.4 Hz, 1H), 8.04 (d, *J* = 7.0 Hz, 1H), 7.95 – 7.84 (m, 5H), 7.74 (dd, *J* = 18.3, 8.2 Hz, 2H), 7.60 – 7.46 (m, 6H), 7.41 – 7.27 (m, 5H), 7.19 (d, *J* = 2.5 Hz, 1H), 7.04 (ddd, *J* = 8.2, 7.0, 1.2 Hz, 1H), 6.94 (ddd, *J* = 8.0, 7.0, 1.0 Hz, 1H), 4.87 (t, *J* = 5.4 Hz, 1H), 4.52 (dt, *J* = 8.6, 4.2 Hz, 1H), 4.22 (d, *J* = 6.3 Hz, 1H), 3.58 (d, *J* = 2.3 Hz, 3H), 3.40 (dd, *J* = 13.8, 6.2 Hz, 6H), 3.26 (s, 2H), 3.20 – 3.12 (m, 2H), 2.20 (s, 3H), 1.50 (d, *J* = 2.7 Hz, 2H), 1.16 (d, *J* = 2.8 Hz, 2H).

^13^C NMR (101 MHz, DMSO) δ 171.68, 171.23, 170.67, 166.42, 155.86, 149.52, 143.30, 142.58, 141.14, 140.06, 136.75, 136.54, 134.58, 134.16, 133.85, 132.00, 131.90, 129.03, 128.48, 128.08, 127.92, 127.73, 127.45, 127.23, 127.00, 126.78, 126.50, 125.97, 125.76, 124.20, 124.09, 121.37, 119.22, 118.74, 113.60, 112.68, 111.71, 110.58, 109.18, 69.08, 68.87, 61.92, 56.12, 54.14, 35.13, 32.77, 31.82, 19.17, 16.14.

MS (ESI, positive) m/z calculated for C_54_H_52_F_2_N_7_O_9_ [M+H]^+^ : 980.38, found: 980.41.

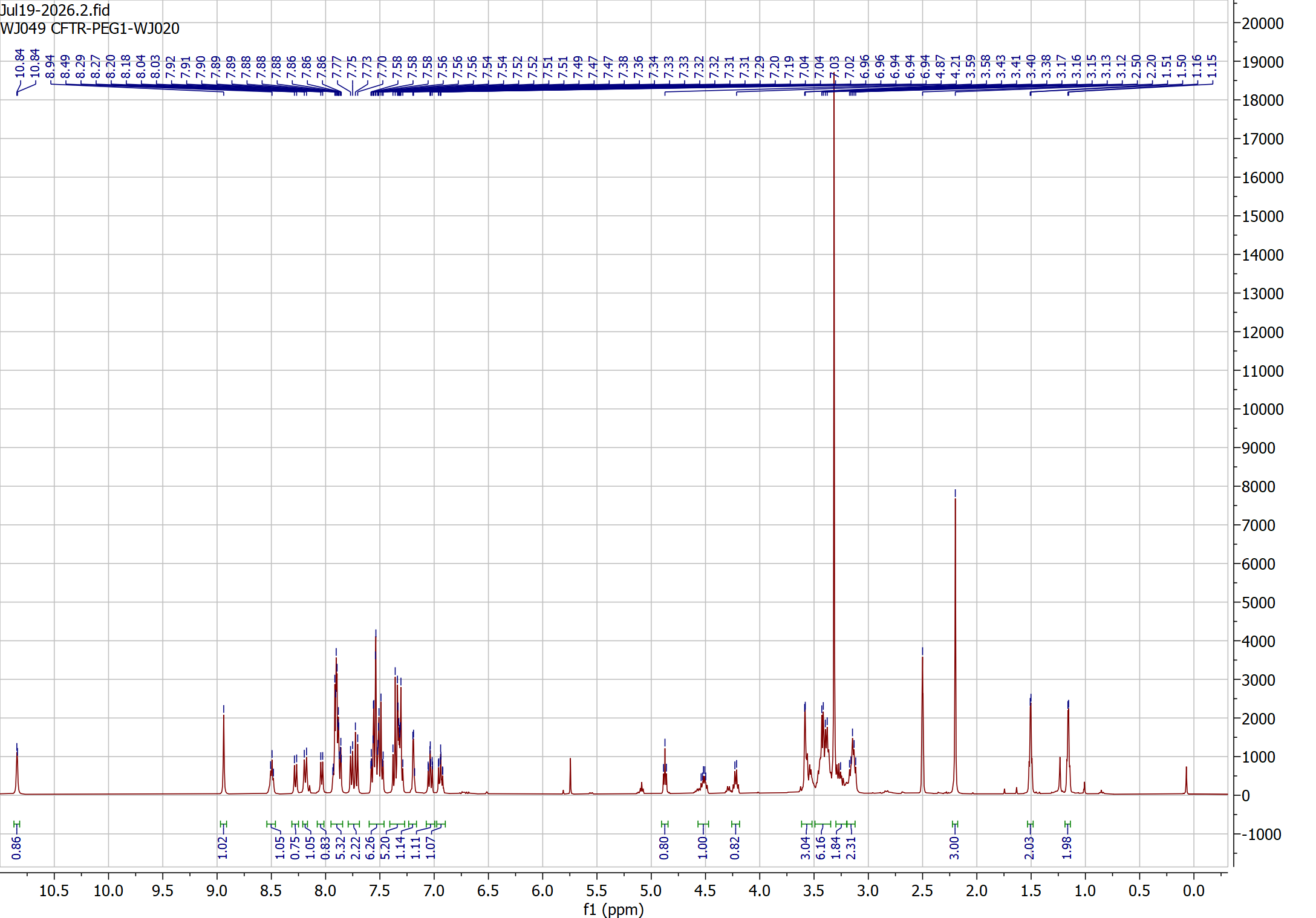

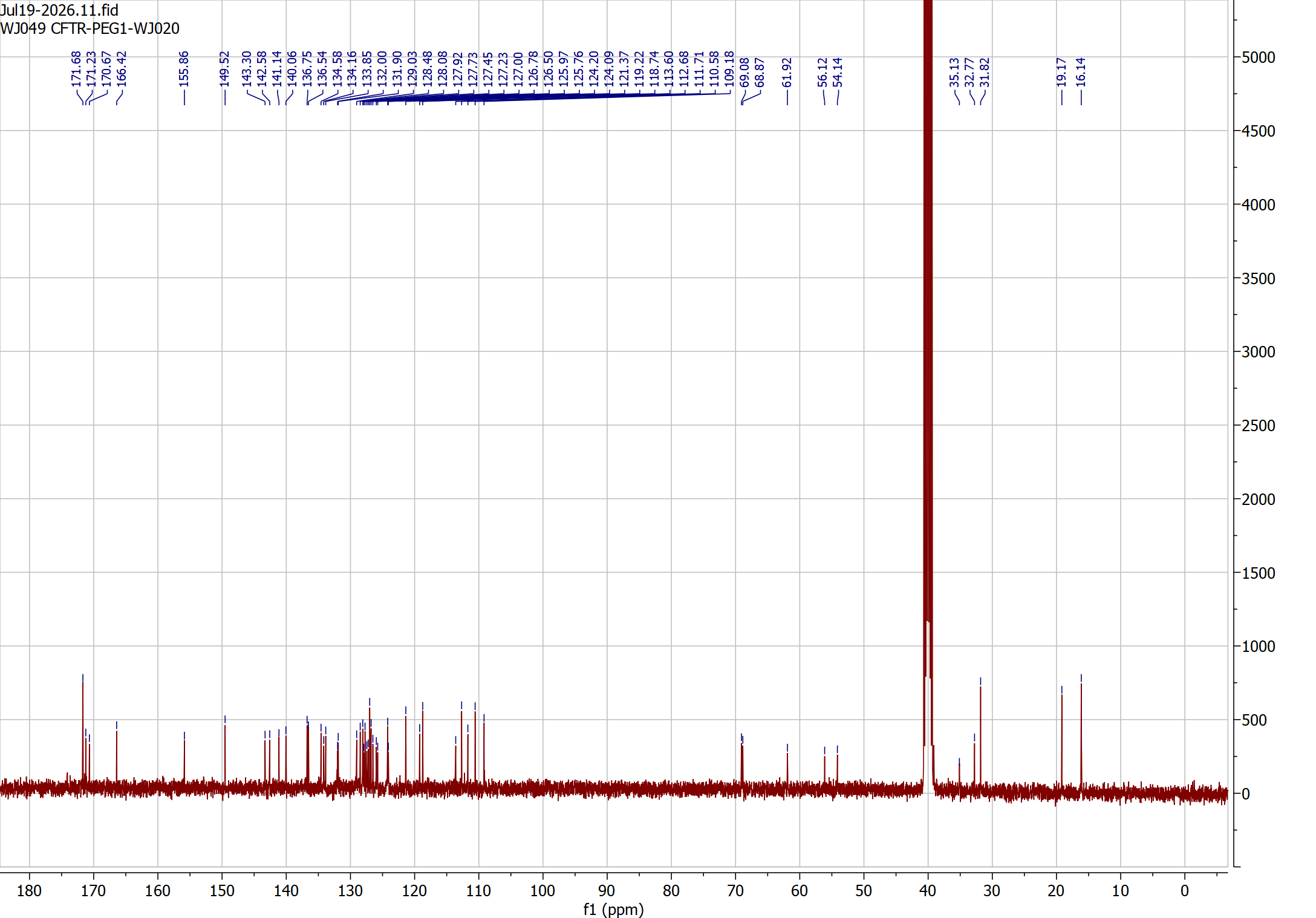

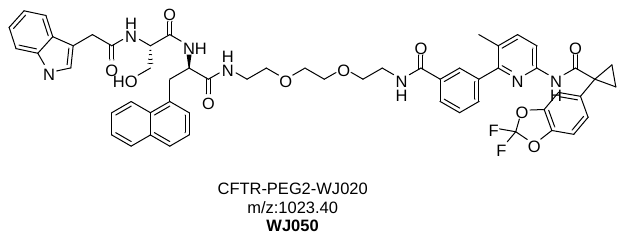

**WJ050**

^1^H NMR (400 MHz, DMSO) δ 10.84 (d, *J* = 2.5 Hz, 1H), 8.94 (s, 1H), 8.53 (t, *J* = 5.6 Hz, 1H), 8.27 (d, *J* = 8.3 Hz, 1H), 8.19 (d, *J* = 8.5 Hz, 1H), 8.03 (d, *J* = 7.1 Hz, 1H), 7.89 (qd, *J* = 7.1, 3.2 Hz, 5H), 7.74 (dd, *J* = 14.2, 8.1 Hz, 2H), 7.63 – 7.47 (m, 6H), 7.42 – 7.27 (m, 5H), 7.19 (dd, *J* = 4.7, 2.4 Hz, 1H), 7.04 (ddd, *J* = 8.1, 7.0, 1.2 Hz, 1H), 7.00 – 6.89 (m, 1H), 4.87 (t, *J* = 5.4 Hz, 1H), 4.61 – 4.47 (m, 1H), 4.23 (d, *J* = 6.4 Hz, 1H), 3.61 – 3.36 (m, 13H), 3.29 – 3.08 (m, 5H), 2.21 (s, 3H), 1.51 (q, *J* = 4.2 Hz, 2H), 1.16 (q, *J* = 4.2 Hz, 2H).

^13^C NMR (101 MHz, DMSO) δ 171.69, 171.63, 171.21, 170.64, 170.61, 166.37, 155.87, 149.53, 143.30, 142.58, 141.14, 140.06, 136.75, 136.54, 134.58, 134.16, 133.85, 132.01, 131.90, 129.02, 128.49, 128.05, 127.95, 127.73, 127.45, 127.23, 127.00, 126.78, 126.50, 125.97, 125.76, 124.26, 124.19, 124.10, 121.37, 119.21, 118.74, 113.59, 112.68, 111.71, 110.58, 109.20, 109.13, 69.92, 69.30, 69.14, 69.06, 61.94, 56.10, 54.13, 35.17, 32.78, 31.82, 19.17, 16.14.

MS (ESI, positive) m/z calculated for C_56_H_56_F_2_N_7_O_10_ [M+H]^+^ : 1024.41, found: 1024.41.

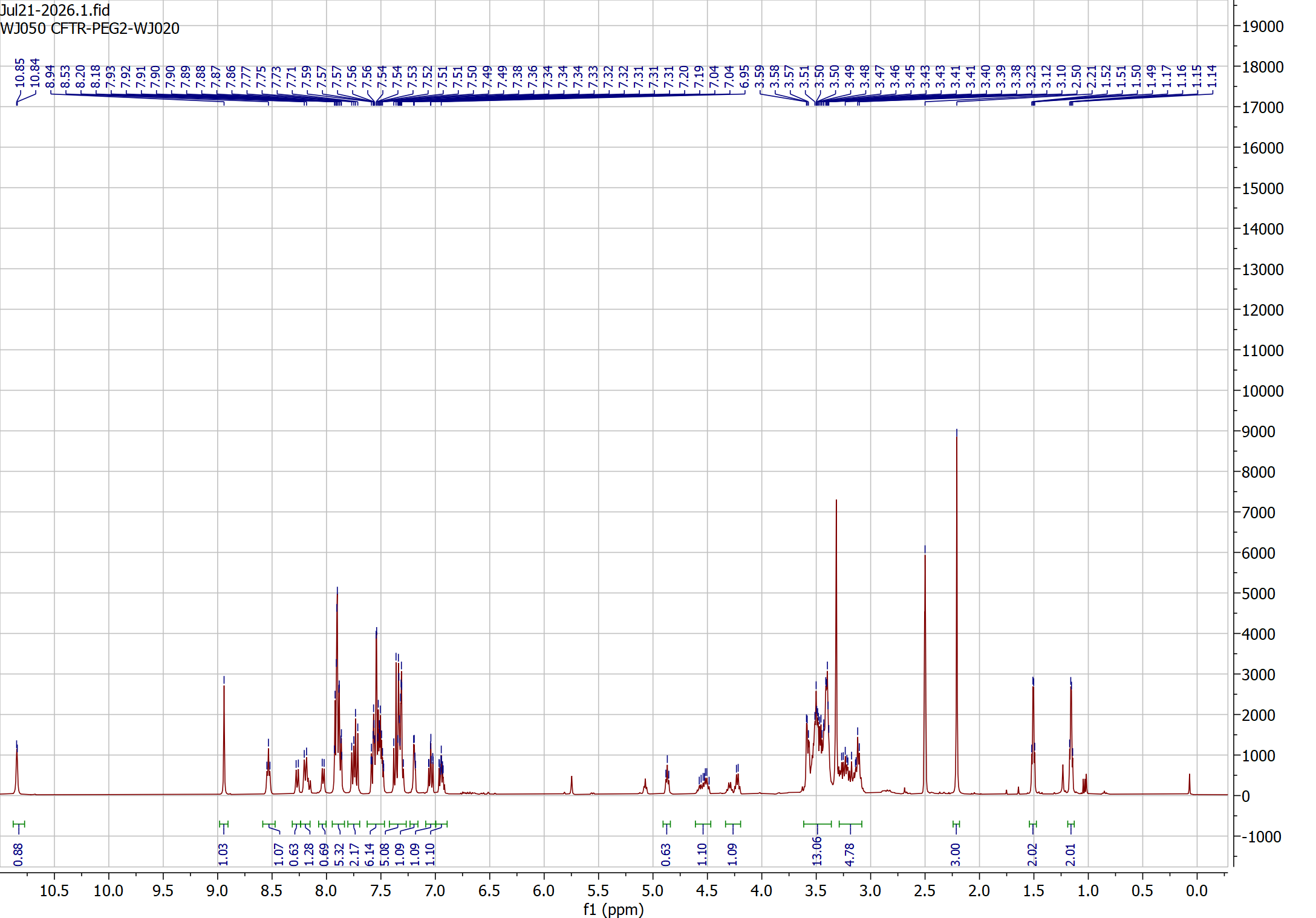

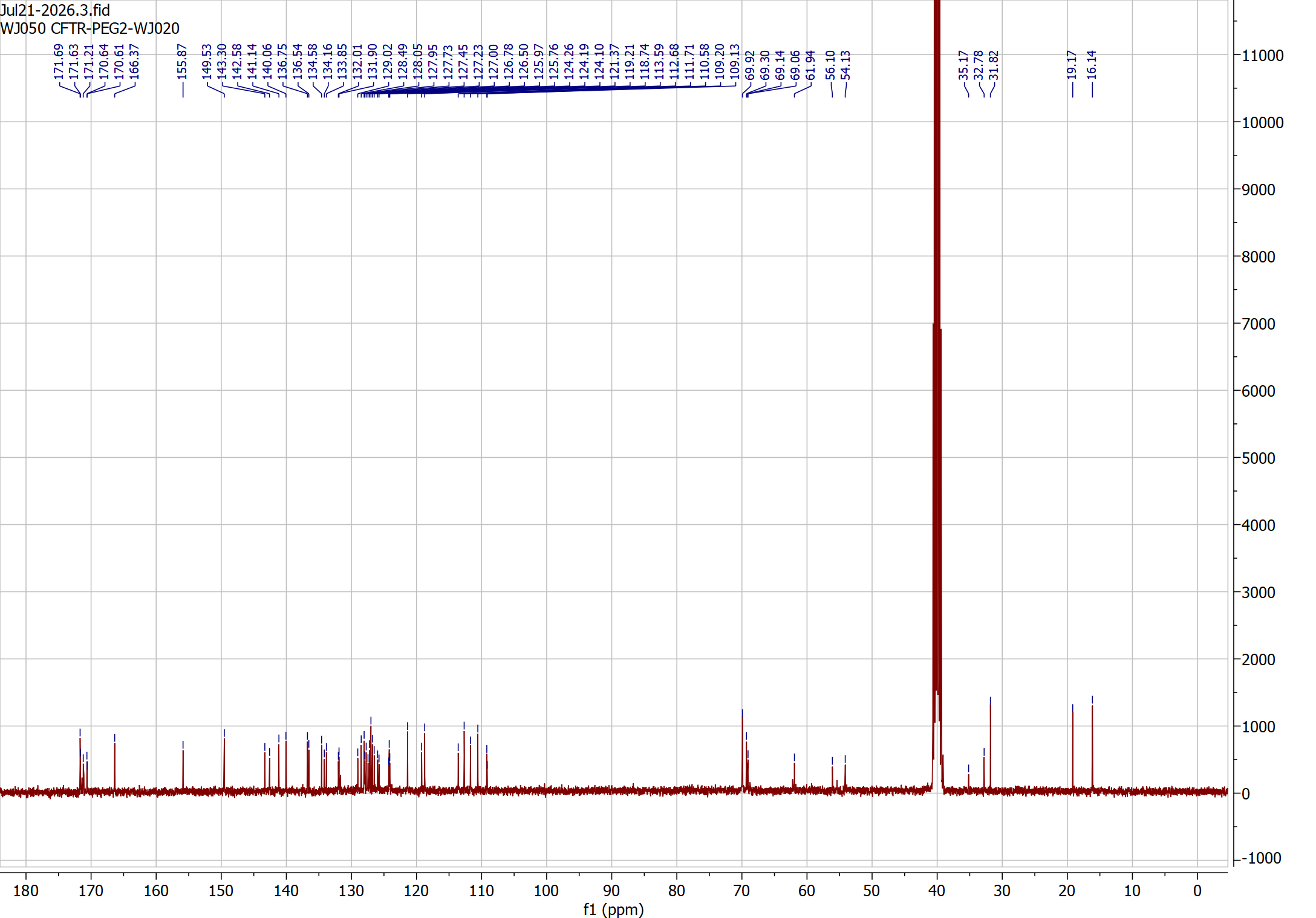

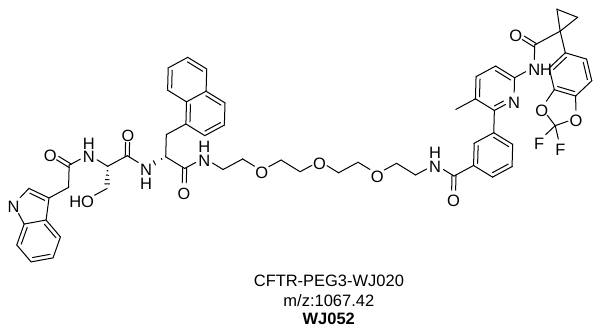

**WJ052**

^1^H NMR (400 MHz, DMSO) δ 10.84 (d, *J* = 2.2 Hz, 1H), 8.94 (s, 1H), 8.53 (t, *J* = 5.6 Hz, 1H), 8.26 (d, *J* = 8.3 Hz, 1H), 8.19 (d, *J* = 8.3 Hz, 1H), 8.02 (d, *J* = 7.1 Hz, 1H), 7.89 (td, *J* = 8.7, 6.9 Hz, 5H), 7.74 (dd, *J* = 14.6, 8.1 Hz, 2H), 7.62 – 7.46 (m, 6H), 7.40 – 7.28 (m, 5H), 7.20 (d, *J* = 2.4 Hz, 1H), 7.04 (ddd, *J* = 8.1, 7.0, 1.2 Hz, 1H), 6.95 (ddd, *J* = 8.0, 7.0, 1.0 Hz, 1H), 4.86 (t, *J* = 5.4 Hz, 1H), 4.56 – 4.47 (m, 1H), 4.22 (d, *J* = 6.4 Hz, 1H), 3.59 (d, *J* = 2.7 Hz, 2H), 3.51 (q, *J* = 5.0 Hz, 6H), 3.47 – 3.35 (m, 8H), 3.30 – 3.08 (m, 5H), 2.21 (s, 3H), 1.51 (q, *J* = 4.2 Hz, 2H), 1.16 (q, *J* = 4.3 Hz, 2H).

^13^C NMR (101 MHz, DMSO) δ 171.69, 171.62, 171.20, 170.63, 166.35, 155.87, 149.53, 143.31, 142.58, 141.14, 140.06, 136.75, 136.54, 134.60, 134.16, 133.85, 132.01, 131.89, 129.03, 128.49, 128.05, 127.95, 127.73, 127.45, 127.23, 127.01, 126.78, 126.51, 125.97, 125.76, 124.19, 124.10, 121.37, 119.21, 118.74, 113.59, 112.68, 111.71, 110.58, 109.20, 70.16, 70.11, 70.02, 69.94, 69.29, 69.05, 61.94, 56.09, 54.12, 35.18, 32.78, 31.82, 19.17, 16.14.

MS (ESI, positive) m/z calculated for C_58_H_60_F_2_N_7_O_11_ [M+H]^+^ : 1068.43, found: 1068.41.

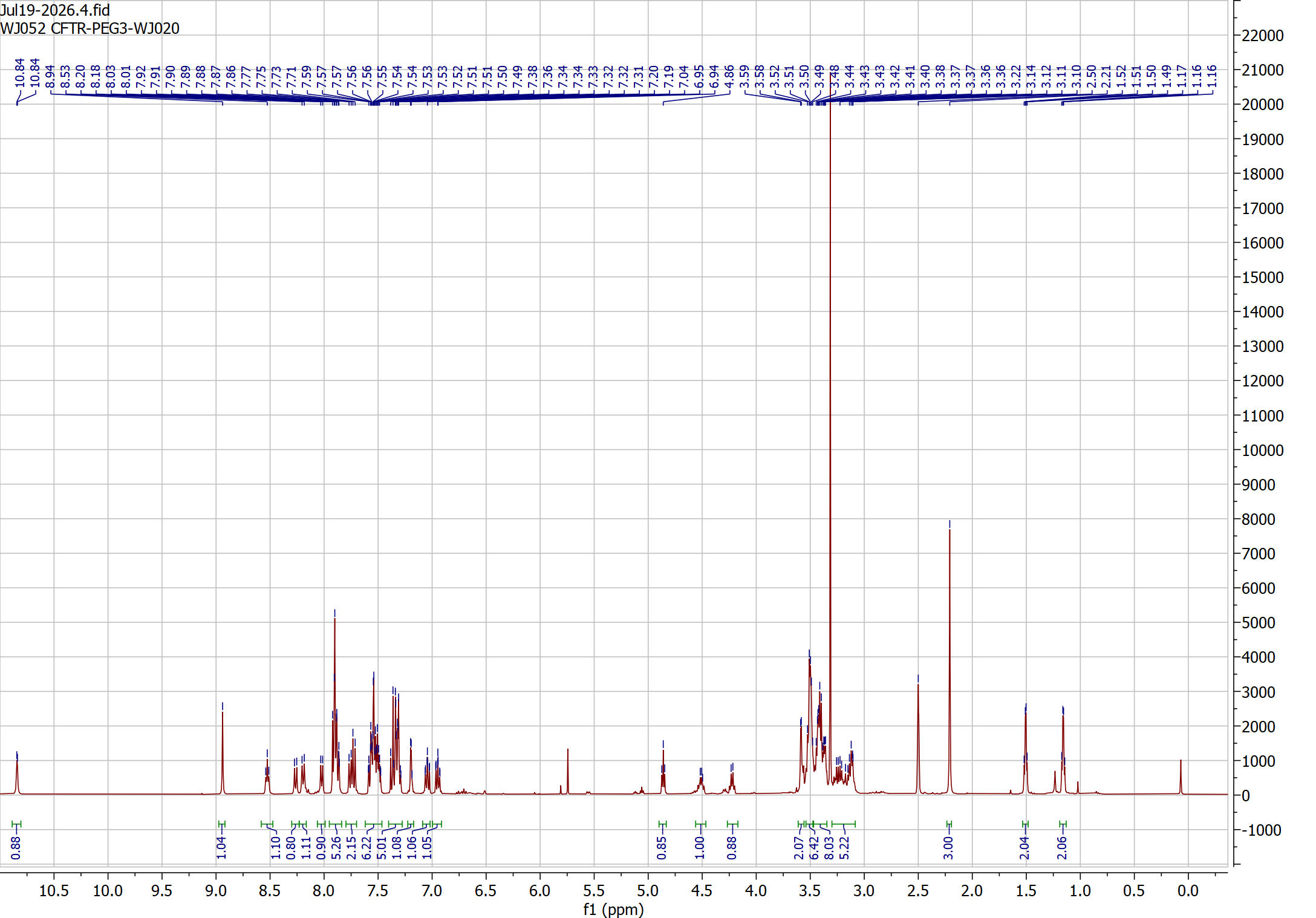

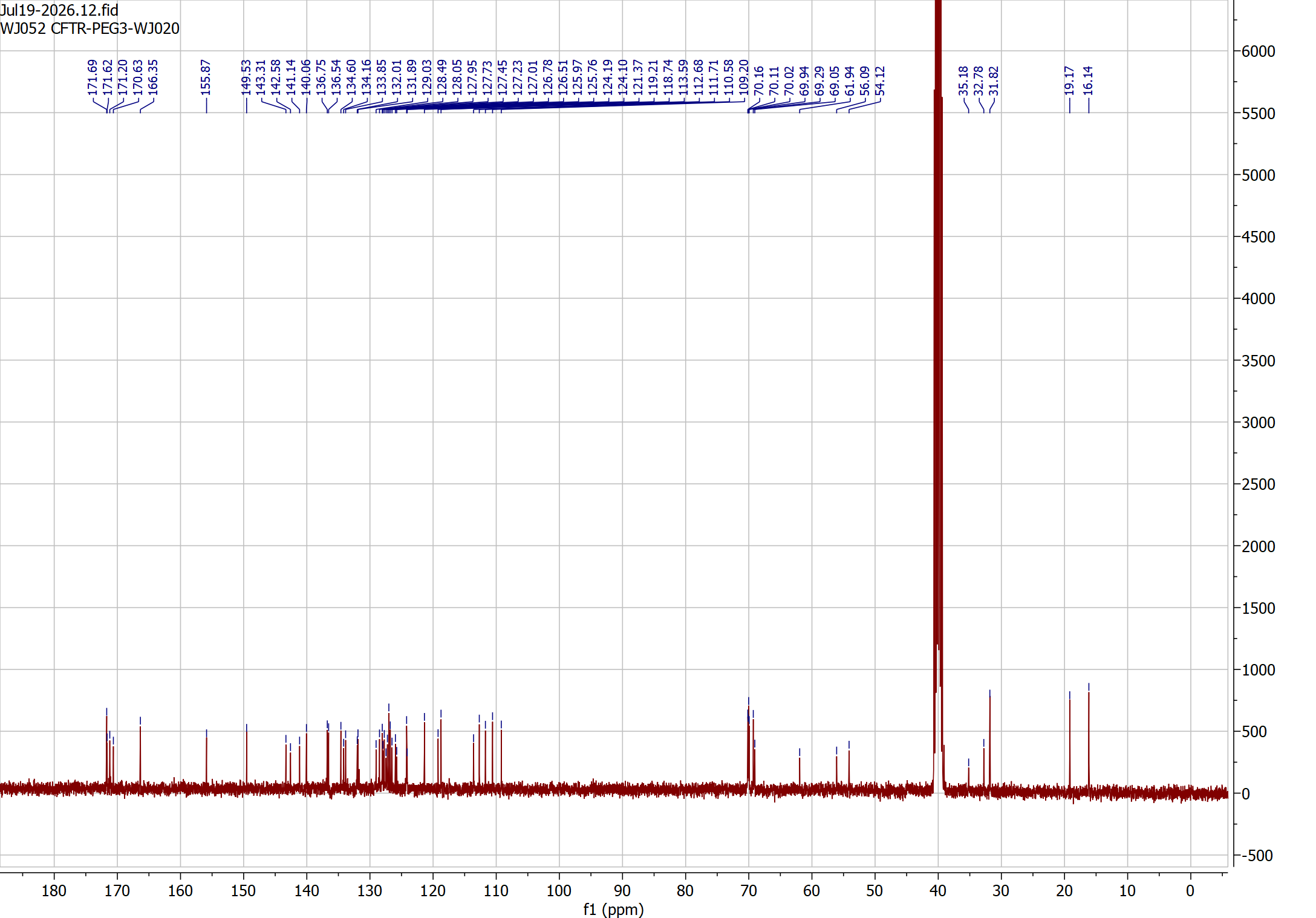

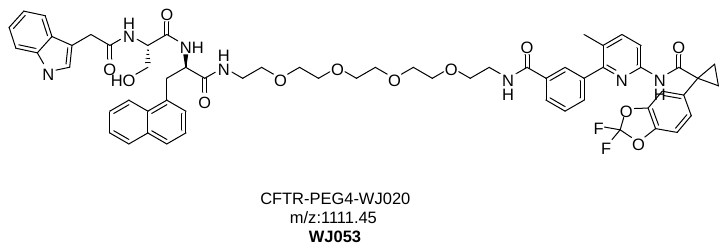

**WJ053**

^1^H NMR (400 MHz, DMSO) δ 10.84 (d, *J* = 2.4 Hz, 1H), 8.94 (s, 1H), 8.52 (t, *J* = 5.6 Hz, 1H), 8.26 (d, *J* = 8.2 Hz, 1H), 8.23 – 8.14 (m, 1H), 8.02 (d, *J* = 7.1 Hz, 1H), 7.89 (td, *J* = 8.8, 6.8 Hz, 5H), 7.74 (dd, *J* = 15.0, 8.1 Hz, 2H), 7.62 – 7.47 (m, 6H), 7.42 – 7.28 (m, 5H), 7.19 (dd, *J* = 5.4, 2.4 Hz, 1H), 7.04 (ddd, *J* = 8.2, 6.9, 1.2 Hz, 1H), 6.95 (ddd, *J* = 8.0, 5.6, 1.3 Hz, 1H), 4.86 (t, *J* = 5.4 Hz, 1H), 4.60 – 4.47 (m, 1H), 4.33 – 4.18 (m, 1H), 3.62 – 3.55 (m, 2H), 3.55 – 3.36 (m, 18H), 3.30 – 3.06 (m, 5H), 2.21 (s, 3H), 1.51 (q, *J* = 4.2 Hz, 2H), 1.16 (q, *J* = 4.3 Hz, 2H).

^13^C NMR (101 MHz, DMSO) δ 171.69, 171.62, 171.20, 170.63, 166.34, 155.88, 149.53, 143.31, 142.58, 141.14, 140.05, 136.75, 136.54, 134.61, 134.17, 133.85, 132.01, 131.89, 129.03, 128.48, 128.04, 127.96, 127.74, 127.46, 127.23, 127.01, 126.78, 126.51, 125.97, 125.76, 124.19, 124.11, 121.37, 119.21, 118.74, 113.59, 112.68, 111.71, 110.58, 109.20, 70.21, 70.17, 70.11, 70.03, 69.94, 69.29, 69.05, 61.94, 56.09, 54.12, 35.19, 32.78, 31.82, 19.17, 16.14.

MS (ESI, positive) m/z calculated for C_62_H_64_F_2_N_7_O_12_ [M+H]^+^ : 1112.46, found: 1112.41.

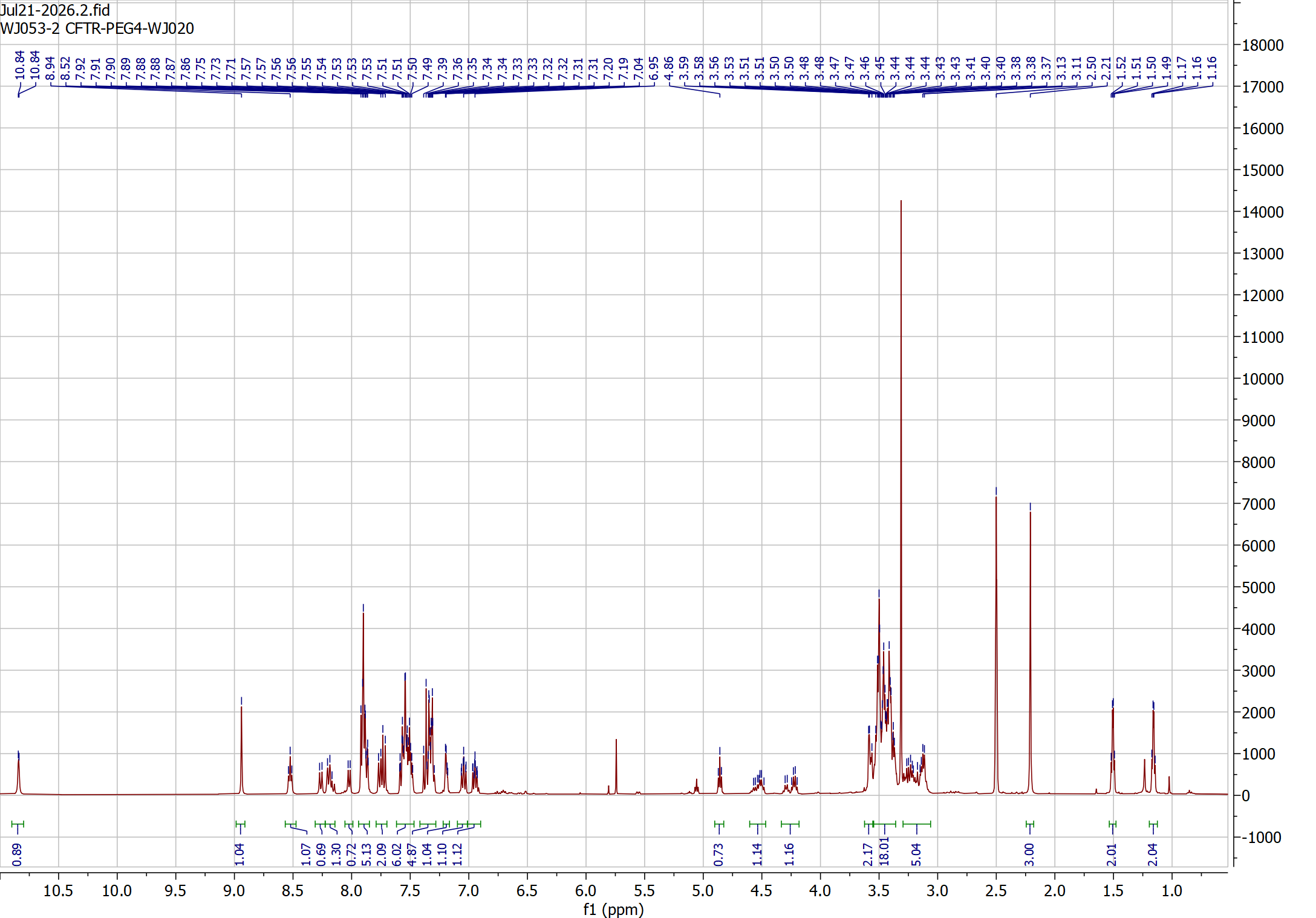

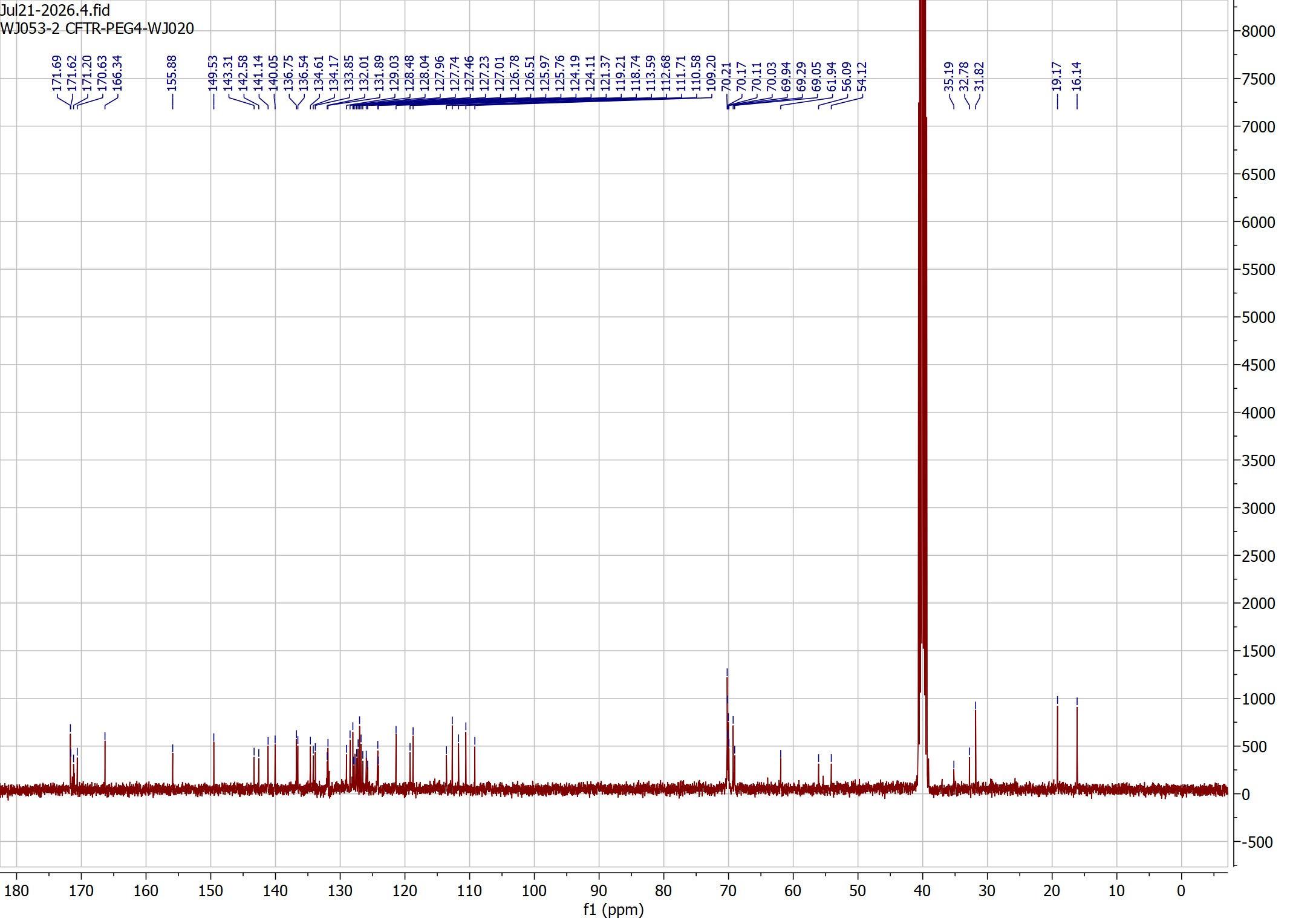

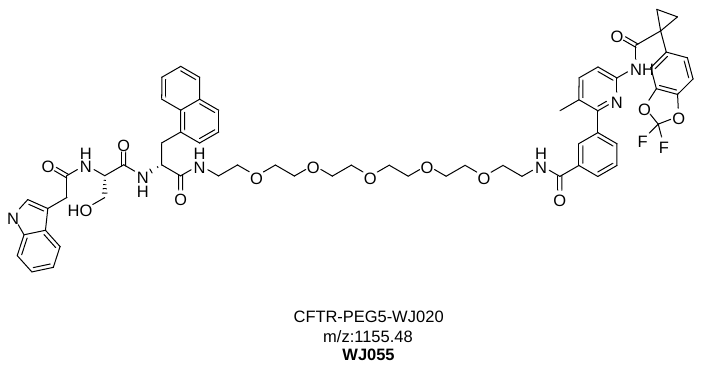

**WJ055**

^1^H NMR (400 MHz, DMSO) δ 10.84 (d, *J* = 2.3 Hz, 1H), 8.94 (s, 1H), 8.52 (t, *J* = 5.6 Hz, 1H), 8.26 (d, *J* = 8.3 Hz, 1H), 8.20 (d, *J* = 8.2 Hz, 1H), 8.02 (d, *J* = 7.1 Hz, 1H), 7.89 (td, *J* = 8.8, 6.7 Hz, 5H), 7.74 (dd, *J* = 15.1, 8.1 Hz, 2H), 7.61 – 7.47 (m, 6H), 7.41 – 7.28 (m, 5H), 7.19 (dd, *J* = 5.6, 2.4 Hz, 1H), 7.05 (ddd, *J* = 8.2, 6.9, 1.3 Hz, 1H), 6.99 – 6.92 (m, 1H), 4.86 (t, *J* = 5.3 Hz, 1H), 4.51 (d, *J* = 5.2 Hz, 1H), 4.22 (d, *J* = 6.4 Hz, 1H), 3.61 – 3.35 (m, 25H), 3.29 – 3.21 (m, 2H), 3.13 (dd, *J* = 9.9, 4.4 Hz, 2H), 2.21 (s, 3H), 1.51 (q, *J* = 4.2 Hz, 2H), 1.16 (q, *J* = 4.3 Hz, 2H).

^13^C NMR (101 MHz, DMSO) δ 171.69, 170.63, 166.34, 155.88, 149.53, 143.31, 142.58, 141.14, 140.05, 136.76, 136.54, 134.61, 134.17, 131.89, 129.03, 128.48, 128.05, 127.23, 127.01, 126.78, 126.51, 125.98, 125.76, 124.19, 121.37, 119.21, 118.74, 113.58, 112.69, 111.71, 110.59, 109.20, 70.21, 70.11, 70.04, 69.29, 56.09, 54.13, 32.78, 31.82, 19.17, 16.14.

MS (ESI, positive) m/z calculated for C_62_H_69_F_2_N_7_O_13_ [M+H]^+^ : 1156.48, found: 1156.51.

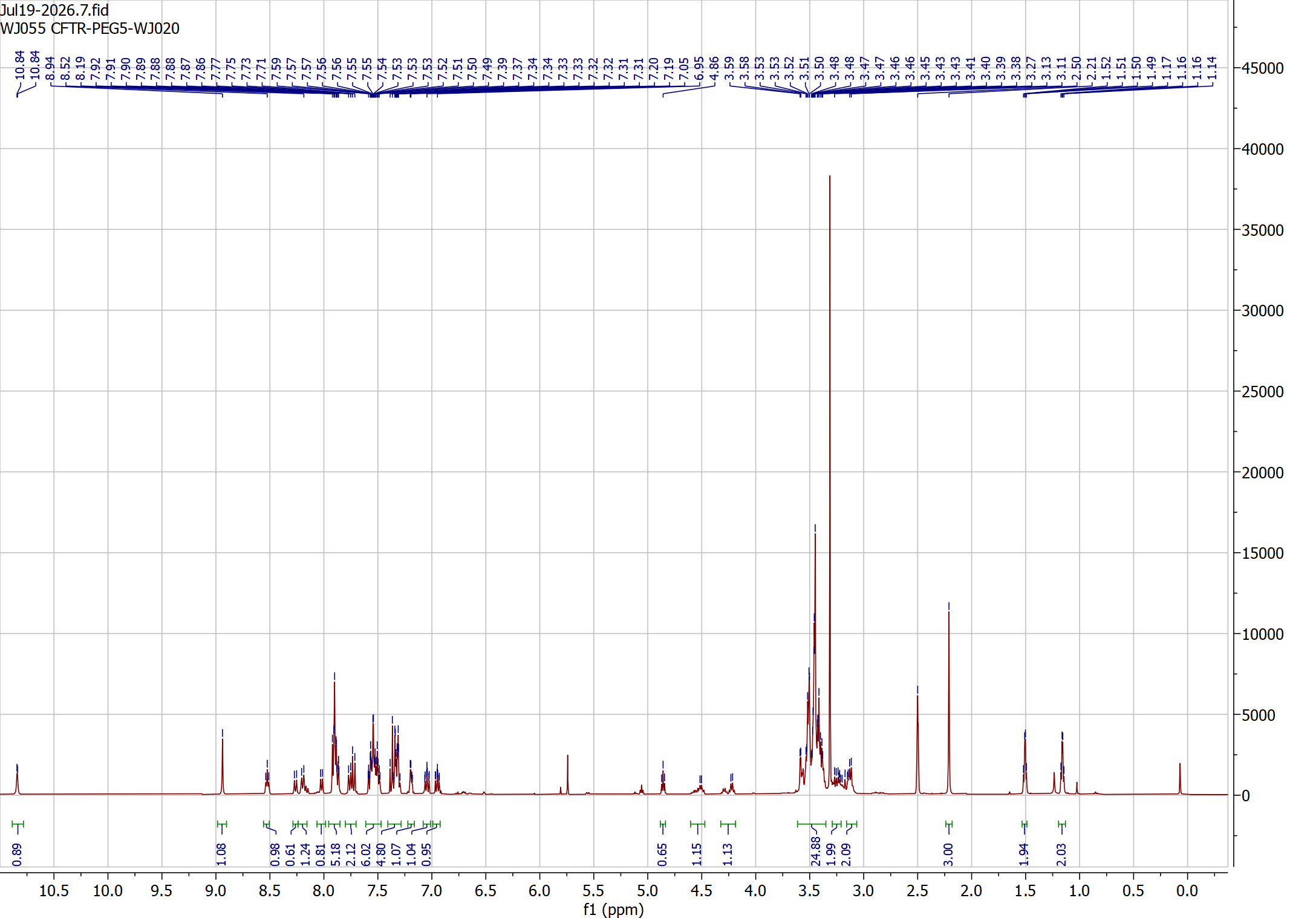

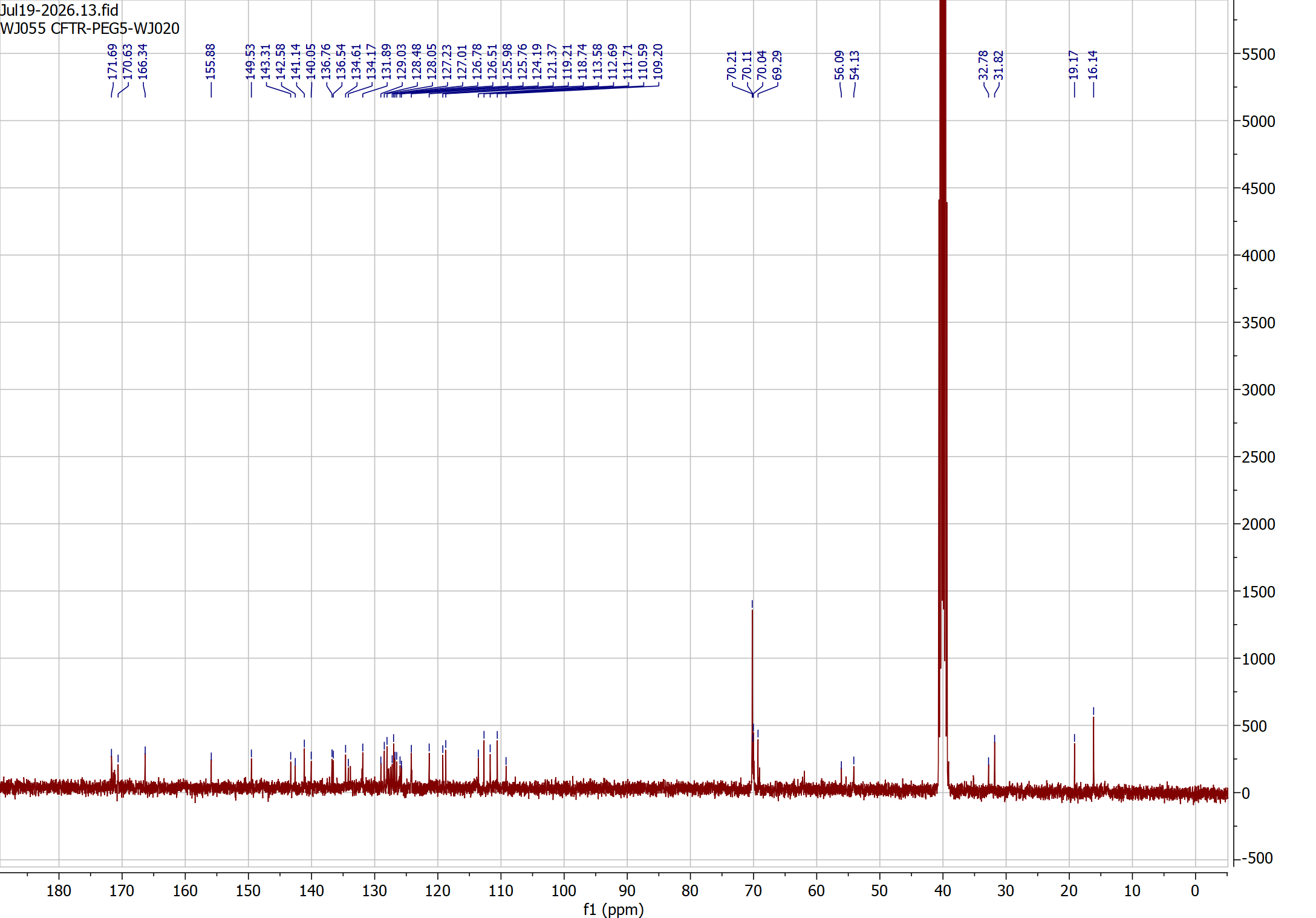
